# Accounting for Gene Flow from Unsampled ‘Ghost’ Populations while Estimating Evolutionay History under the Isolation with Migration Model

**DOI:** 10.1101/733600

**Authors:** Arun Sethuraman, Melissa Lynch

## Abstract

Unsampled or extinct ‘ghost’ populations leave signatures on the genomes of individuals from extant, sampled populations, especially if they have exchanged genes with them over evolutionary time. This gene flow from ‘ghost’ populations can introduce biases when estimating evolutionary history from genomic data, often leading to data misinterpretation and ambiguous results. Here we assess these biases while accounting, or not accounting for gene flow from ‘ghost’ populations under the Isolation with Migration (IM) model. We perform extensive simulations under five scenarios with no gene flow (Scenario A), to extensive gene flow to- and from- an unsampled ‘ghost’ population (Scenarios B, C, D, and E). Estimates of evolutionary history across all scenarios A-E (effective population sizes, divergence times, and migration rates) indicate consistent a) under-estimation of divergence times between sampled populations, (b) over-estimation of effective population sizes of sampled populations, and (c) under-estimation of migration rates between sampled populations, with increased gene flow from the unsampled ‘ghost’ population. Without accounting for an unsampled ‘ghost’, summary statistics like *F*_*ST*_ are under-estimated, and *π* is over-estimated with increased gene flow from the‘ghost’. To show this persistent issue in empirical data, we use a 355 locus dataset from African Hunter-Gatherer populations and discuss similar biases in estimating evolutionary history while not accounting for unsampled ‘ghosts’. Considering the large effects of gene flow from these ‘ghosts’, we propose a multi-pronged approach to account for the presence of unsampled ‘ghost’ populations in population genomics studies to reduce erroneous inferences.

## Introduction

Studies that apply population genetics methods to infer evolutionary history, including those in the fields of conservation, agriculture, evolution, and anthropology often begin with a sampling strategy. As a rule of thumb, we would expect that the more extensive the sampling of individuals across their geographical range, or according to the biological question at hand, the better the resolution of population genetic analyses. However, in any such study, genomic data collected harbors signatures of evolutionary processes of drift, selection, and gene flow between unsampled ‘ghost’ and sampled populations (Beerli, 2004). A ‘ghost’ population can be defined as one or more extant or extinct populations that have, or continue to exchange genes with sampled populations. Such ‘ghost’ populations are ubiquitous, and have increasingly been discovered across numerous species complexes. For example, several studies of the evolutionary history of African hunter-gatherers (Lachance et al., 2012), (Hey et al., 2018), (Durvasula & Sankararaman, 2019) identify significant gene flow from an unsampled archaic population (diverged from modern humans around the same time as Neanderthals) into multiple modern Hunter-Gatherer lineages. Similar studies in black rats (Konečný et al., 2013), *Brachypodium sylvaticum* bunchgrass (Rosenthal, Ramakrishnan, & Cruzan, 2008), bonobos (Kuhlwilm, Han, Sousa, Excoffier, & Marques-Bonet, 2019) and modern humans (Nielsen et al., 2017) also describe the occurrence of unsampled ‘ghost’ populations in their respective systems. The presence of such unsampled genomic variation in sampled genomic data is a result of either (a) incomplete population sampling on the part of the researcher, (b) population extinction or decline, or (c) the amount of gene flow from such unsampled, or extinct populations over evolutionary time.

Importantly, not accounting for such unsampled ‘ghost’ population gene flow into extant sampled populations can lead to erroneous estimates from summary statistics, or phylogenetic/mutation model-based methods, and thus, erroneous conclusions about the evolutionary history of the species. Surprisingly, the studies that account for unsampled ‘ghosts’ are still part of the minority in population genomics research and publications.

For example, under an evolutionary scenario where two populations of a species don’t directly exchange genes between each other but exchange genes bidirectionally with an unsampled ‘ghost’, one would expect that measures of population differentiation (*F*_*ST*_) between sampled extant populations would be much lower than expected under a model where gene flow from the ‘ghost’ is unaccounted for. Similarly, other summary statistics, such as the allele frequency distribution (Tajima’s D), heterozygosity, nucleotide diversity (*π*), effective population size (*N*_*e*_), and estimates of time since most recent common ancestor (*t*_*MRCA*_) can all be biased depending on the degree of gene flow from unsampled ‘ghost’ populations.

Model-based methods for estimating population structure and gene flow, including Bayesian or maximum likelihood approaches, as implemented in software such as `structure` (Pritchard, Stephens, & Donnelly, 2000), `ADMIXTURE` (Alexander, Novembre, & Lange, 2009), `MIGRATE-n` (Beerli & Felsenstein, 2001), and `IMa2p` (Sethuraman & Hey, 2016) are also not infallible to the effects of ‘ghost’ gene flow, in that identical estimates of phylogenetic and mutational history may be achieved despite a number of unique demographic histories (Lawson, Van Dorp, & Falush, 2018). For example, (Lawson et al., 2018) describe three scenarios of evolutionary history - one of recent admixture from four divergent populations, one of gene flow from a ‘ghost’ population, and one of recent bottlenecks in sampled, extant populations. In the three different scenarios, the same population structure and admixture proportions are inferred using the programs `structure` or `ADMIXTURE`. Correspondingly, not accounting for ‘ghost’ gene flow has been known to bias estimates of migration rates (Hey et al., 2018) and divergence times (Lachance et al., 2012) while using other model-based estimators of evolutionary history.

Biases in estimates of effective population sizes and migration rates in the presence of the unsampled ‘ghost’ populations were previously investigated under an Island Model (Beerli, 2004; Slatkin, 2005). Using an ‘n’-island model (Slatkin, 1985), where each of ‘n’ populations of constant size can exchange genes at constant rates with the ‘n-1’ other populations, Beerli estimated the effect of the magnitude and direction of migration in the presence of a ‘ghost’ population. They simulated three populations of identical effective population size and per generation mutation rates under this ‘n’ island model, such that each set of populations either had high or low magnitude of unidirectional or bidirectional migration with the ‘ghost’ population. Beerli also estimated the effect of the number of ‘ghost’ populations by sampling two populations out of a larger, varied set of unsampled populations. Each of these datasets was then analyzed using the `MIGRATE-n` software. Beerli’s study identified several effects of the ‘ghost’ population gene flow, including (1) a higher migration rate, both unidirectional and bidirectional to and from the ‘ghost’ population, led to an overestimation of migration rates between the sampled populations, (2) increasing the number of ‘ghost’ populations increased the bias in estimates of population sizes, but had little effect on the migration rate estimates, and (3) increasing the number of sampled loci did not improve or affect the estimation of migration rates.

Here we extend Beerli’s study to a more complex model of evolutionary history, popularly termed the Isolation with Migration (IM) model (Hey & Nielsen, 2004, 2007; Nielsen & Wakeley, 2001). This class of models is widely used to model divergence with gene flow, where two sampled populations or subpopulations have diverged from the ancestor, and maintain gene flow via exchange of migrants postdivergence. Numerous tools have been developed to estimate evolutionary history under the IM model, including the IM suite - `IMa2p`, and `IMa3` (Sethuraman & Hey, 2016; Hey et al., 2018), `MIST` (Chung & Hey, 2017), and `dadi` (Gutenkunst, Hernandez, Williamson, & Bustamante, 2009). The IM suite of tools are genealogy samplers that use a Metropolis Coupled Markov Chain Monte Carlo (MCMCMC) to explore the parameter space of evolutionary demographic parameters, propose updates to parameters and genealogies, and examine the posterior density distribution of the set of estimable parameters given genomic data from two or more sampled populations. These programs estimate divergence times, migration rates, and effective population sizes of all populations included in the model.

Importantly, these programs also have the ability to allow for the presence of a single ‘ghost’ population, where a population is added to the model and assumed to be the outgroup to all sampled populations (this aligns with Beerli’s (2001) finding that the addition of more than one ‘ghost’ population doesn’t significantly affect estimates of migration rates), and estimate parameters of the IM model under various phylogenetic models. This allows us to compare and contrast how accounting or not-accounting for the presence of a ‘ghost’ population could potentially bias estimates of divergence times, effective population sizes, and migration rates between sampled extant populations.

Our study utilizes an extensive set of simulations under the IM model, with (1) varying degrees of unidirectional and bidirectional gene flow from unsampled ‘ghost’ populations, (2) varying the number of sampled genomic loci, to quantify the biases in (a) widely-used summary statistics in population genomics, and (b) estimates of evolutionary history under the IM model.

Additionally, as a proof of concept, we use 355 nuclear genomic loci under four demographic IM models to infer the demographic history of African Hunter-Gatherer populations. The evolutionary history of African Hunter-Gatherers has been previously summarized by (Lachance et al., 2012), wherein a genome wide study of the Hadza and Sandawe indicated signals of (a) population bottlenecks in the Hadza, indicated by proportion of polymorphic sites, (b) ancient admixture determined by the *S*∗ statistic. Here we utilize a model-based approach to understand these observations from the Hadza, Sandawe, Baka, and Yoruba to show how the addition of an unsampled ‘ghost’ population aids in accurate estimates of demographic parameters using `IMa2p`, comparable to the results of (Hey et al., 2018).

## Methods

### Simulations

The `ms` software (Hudson, 2002) was used to generate genomic data under IM model ‘versions’ of the Beerli (Beerli, 2004) simulations (Fig. 1). This software uses coalescent methods to generate genomic data by sampling haplotypes from populations evolving under various models. In sum, three populations were assumed in all models: *N*_0_, and *N*_1_ (extant, sampled), and *N*_*ghost*_ (unsampled ‘ghost’). All populations were assumed to have a constant effective population size (mutation rate per locus per generation scaled population size, *θ* = 4*N*_*e*_*μ* = 0.01). Populations *N*_0_ and *N*_1_ were assumed to have diverged from their common ancestor (*N*_*a*_) at *t*_0_=0.5 (divergence time is scaled by mutation rate per locus per generation as t/*μ*). Population *N*_*ghost*_ diverged from *N*_*a*_ at a time *t*_1_=2. Ten individuals were sampled per population.

**Figure 1:**
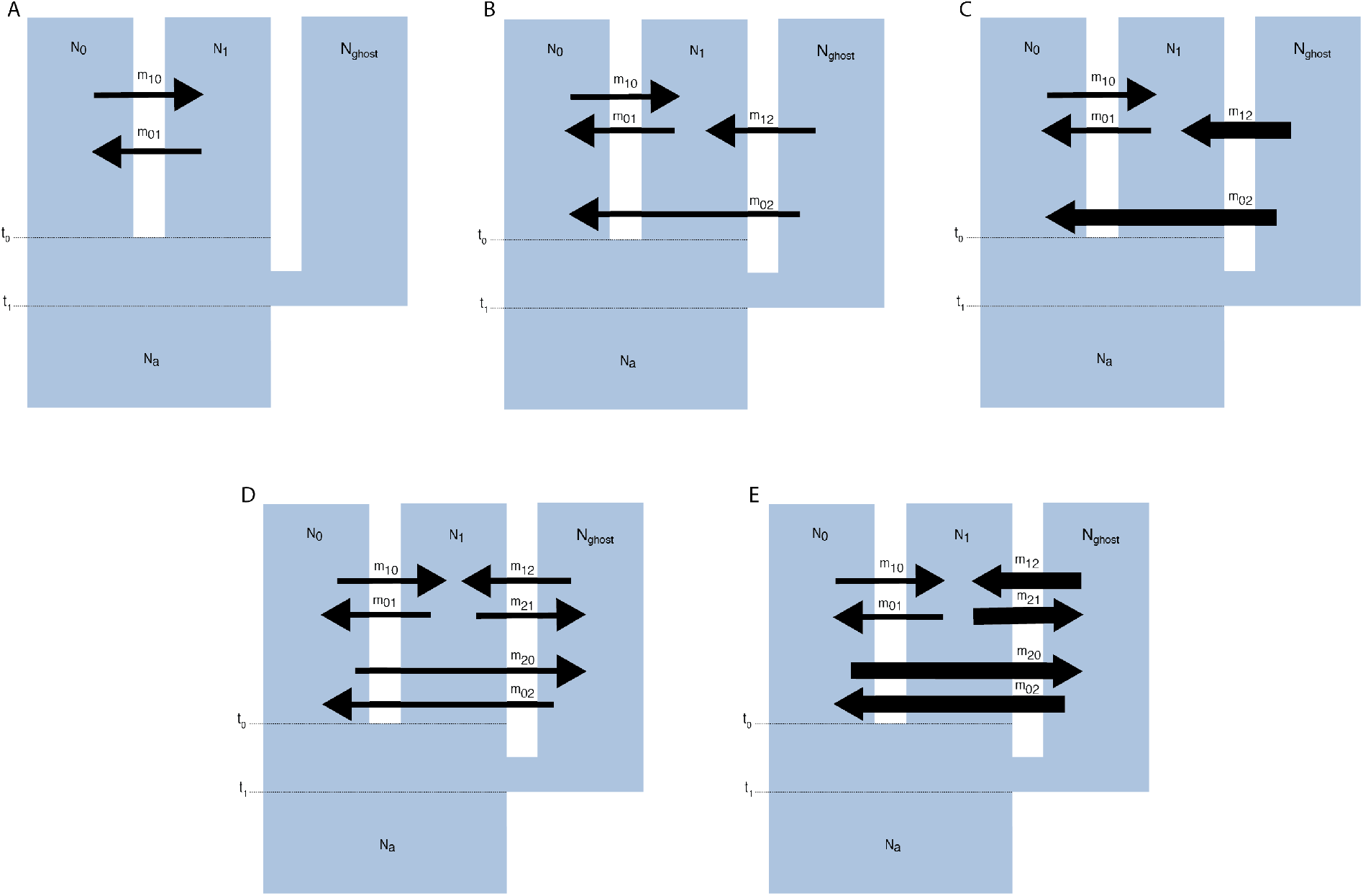
Models A-E under the Isolation with Migration Model. *N*_0_ and *N*_1_ are the two sampled populations (which diverged at time *t*_0_), *N*_*ghost*_ is the unsampled ‘ghost’ population, and *N*_*a*_ is the common ancestor of *N*_0_, *N*_1_ and ‘ghost’ (which diverged at time *t*_1_). The arrows indicate either high (bold arrow) or low (small arrow) levels of gene flow, as well as direction. Model A is the null model, under which there is no gene flow from the ‘ghost’ into the two sampled populations, while models B and C represent low and high unidirectional gene flow from the ‘ghost’ into the two sampled populations, respectively. Models D and E represent models of low and high bi-directional gene flow to and from the ‘ghost’ into the two sampled populations, respectively.

There were five different migration models (Fig. 1) simulated under this IM model. Under model A, only populations *N*_0_ and *N*_1_ exchange genes since divergence from D at a rate of 4*N*_0_*m*_0→1_ = 4*N*_1_*m*_1→0_ = 1.0. This is equivalent to one migrant every fourth generation. Neither *N*_0_ nor *N*_1_ exchange any migrants with the ‘ghost’ population *N*_*ghost*_. Under model B, *N*_0_ and *N*_1_ exchange migrants at a rate of 4*N*_0_*m*_0→1_ = 4*N*_1_*m*_1→0_ = 1.0 as in model A, but individuals also emigrate out of the ‘ghost’ *N*_*ghost*_ into *N*_0_ and *N*_1_ at the same rate (4*N*_*ghost*_*m*_1→2_ = 4*N*_*ghost*_*m*_0→2_ = 1.0). Under model C, the ‘ghost’ population *N*_*ghost*_ exchanges 4*N*_*ghost*_*m*_1→2_ = 4*N*_*ghost*_*m*_0→2_ = 10.0 migrants every fourth generation, while *N*_0_ and *N*_1_ continue to exchange migrants at the same low rate (4*N*_0_*m*_0→1_ = 4*N*_1_*m*_1→0_ = 1.0). Under model D, all three populations exchange migrants at the same low rate (4*N*_*e*_*m* = 1.0). Under the last model E, the ghost *N*_*ghost*_ exchanges migrants bidirectionally at a high rate (4*N*_*e*_*m* = 10.0), while the two sampled populations *N*_0_ and *N*_1_ still exchange migrants at a low rate of 4*N*_0_*m*_0→1_ = 4*N*_1_*m*_1→0_ = 1.0. Under all simulated models, ten individuals were sampled from each population, and separate data-sets were constructed with two or five genomic loci. Ten replicate datasets were simulated under each model, to construct confidence intervals around estimates.

### Summary Statistics

The R PopGenome (Pfeifer, Wittelsbuerger, Ramos-Onsins, & Lercher, 2014) package was used to calculate measures of population differentiation measured as Weir and Cockerham’s *F*_*ST*_, Tajima’s D, nucleotide diversity (*π*), effective population size (*N*_*e*_) measured as the Watterson estimator of genetic diversity (*θ*), and the number of segregating sites (*S*). The PopGenome package was chosen owing to its implementation of numerous summary statistics by invoking a single function.

### Estimates of evolutionary history - Simulations

Evolutionary history was then estimated with the `IMa2p` (Sethuraman and Hey, 2014) software under three separate models: (1) a two population model, wherein the simulated genomic data was down-sampled to only include populations *N*_0_ and *N*_1_ (2) a three population model, wherein all three populations, *N*_0_, *N*_1_, and *N*_*ghost*_ were included in a population model with *N*_0_ and *N*_1_ sharing a more recent common ancestor, and *N*_*ghost*_ as the outgroup and (3) a two population model (same as model 1), but with the addition of an outgroup ‘ghost’ population. Model (3) is the ideal scenario, wherein users can utilize the option of adding an unsampled ‘ghost’ population to their estimation of evolutionary history. This option has been conveniently written into `IMa2p` (Sethuraman & Hey, 2016) and `IMa3` (Hey et al., 2018).

Prior values for effective population sizes, divergence times, and migration rates were set using estimates of Watterson’s *θ*. Upper bounds on *θ* estimates were set to be five times the geometric mean of Watterson’s *θ* across all loci, the upper bound on the divergence times were set to two times the geometric mean of Watterson’s *θ* across all loci, and the upper bound on migration rates was set to 2 divided by the geometric mean of Watterson’s *θ* across all loci, according to the recommendations of (Hey, 2011a). All runs were performed using 10 chains distributed across 56 processors (total of 560 chains), discarding 1 × 10^6^ MCMC iterations as burn-in, followed by sampling 100,000 genealogies. It was ensured that all chains were sufficiently mixed by swapping genealogies across chains (swap rates > 0.5) and that the chains had converged prior to sampling genealogies (inferred by observing the autocorrelations between parameter estimates, across iterations, and in effective sample size values.

All sampled genealogies were then used to estimate marginal posterior densities of parameters, and the modes of these marginal distributions and 95% confidence intervals around the modes were computed, and compared to the ‘true’ simulated parameters used in `ms` (Hudson, 2002).

### Empirical Data

African Hunter-Gatherer populations (Hadza, Sandawe) are a culturally diverse group of indigenous populations who are hypothesized to have diverged from other ancient African modern human lineages (particularly the Pygmy (Baka) and Yoruba (agricultural) populations). The Hadza and Sandawe are both historically Hunter-Gatherer populations from Central Tanzania, and are known to share a complex cultural and demographic history. While the Hadza are hypothesized to have little known cultural confluence with other native populations, the Sandawe, on the other hand, have historically admixed with other Northern populations, including the Baka (pygmy), and Yoruba (pastoral/agricultural). (Lachance et al., 2012) used summary statistics to determine signals of (a) population bottleneck in the Hadza (using the relative proportion of polymorphic sites), and (b) archaic admixture with an unknown ancestral species or population, using the *S*∗ statistic.

We filtered the genomes of 20 individuals (5 each of Hadza, Sandawe, Baka, and Yoruba) with filters used by (Gronau, Hubisz, Gulko, Danko, & Siepel, 2011), based on removing recombination hotspots, duplications, syntenic regions with *P. troglodytes*. The filtered diploid SNP loci were then phased with `fastPHASE` to obtain haplotypes across populations (Scheet & Stephens, 2006). The haplotypes were then filtered further to remove possible recombining segments using a four-gamete test (Hudson & Kaplan, 1985). This produced a total of 355 random, unlinked, putatively neutral loci, which were used in demographic analyses described below.

Demography was then inferred under different models of population history - (1) 2 population IM models with all pairs of populations (Hadza-Sandawe, Sandawe-Baka, Baka-Yoruba, Baka-Hadza, Sandawe-Yoruba, Hadza-Yoruba) assuming that there is no ‘ghost’ population (Fig. 6A), and (2) 2 population IM models with all pairs of populations, assuming that there is an outgroup ‘ghost’ population (Fig. 6B).

The ‘ghost’ population is assumed to be an unsampled outgroup lineage that exchanges genes with sampled populations at constant migration rates, estimated under the IM model. All priors on population sizes, divergence times, and migration rates were set based on the guidelines of (Hey, 2011b), by using the harmonic mean of Watterson’s estimator of *N*_*e*_ across loci (see Supplementary Table 6).Parallel runs of MCMC were then performed with 100, 000 iterations of burn-in, followed by a total run time of 48 hours. Mixing, and convergence were assessed by observing swap rates over runs, acceptance rates of parameter updates, effective sample sizes, and autocorrelations. Convergence of the MCMC was then assessed using `Tracer` (Rambaut, Drummond, Xie, Baele, & Suchard, 2018). Sampled genealogies were used to estimate marginal posterior density distributions of demographic parameters and scaled with a human generation time of 29 years. Likelihood ratio tests were used to determine statistical significance of migration estimates.

## Results

### Effects on the estimation of summary statistics

Estimates of Tajima’s D (Fig. 2A) did not vary with increased migration among populations. The number of segregating sites (S), Watterson’s *θ*, and nucleotide diversity (*pi*) all had similar patterns across models (Fig. 2B, D, E) such that populations with increased migration from the ‘ghost’ show greater estimates of all diversity statistics. Estimates of differentiation between populations (*F*_*ST*_) showed different patterns depending on sampling strategy. A 3 population model reflected higher differentiation among populations where there was no migration between ‘ghost’, with differentiation among populations decreasing as migration rates increased (models B-E) (Fig. 2C). A 2 population model did not show any differences in differentiation between sampled populations. *F*_*ST*_ estimates were higher in models of unidirectional gene flow from ‘ghost’ (models B, C) and lower *F*_*ST*_ in the model with highest bidirectional gene flow (model E) (Fig. 2C). Increasing the number of sampled loci did not change estimated summary statistics, although the differences between high and low migration models were minimized when more loci were sampled (Supplementary Fig. 1).

**Figure 2:**
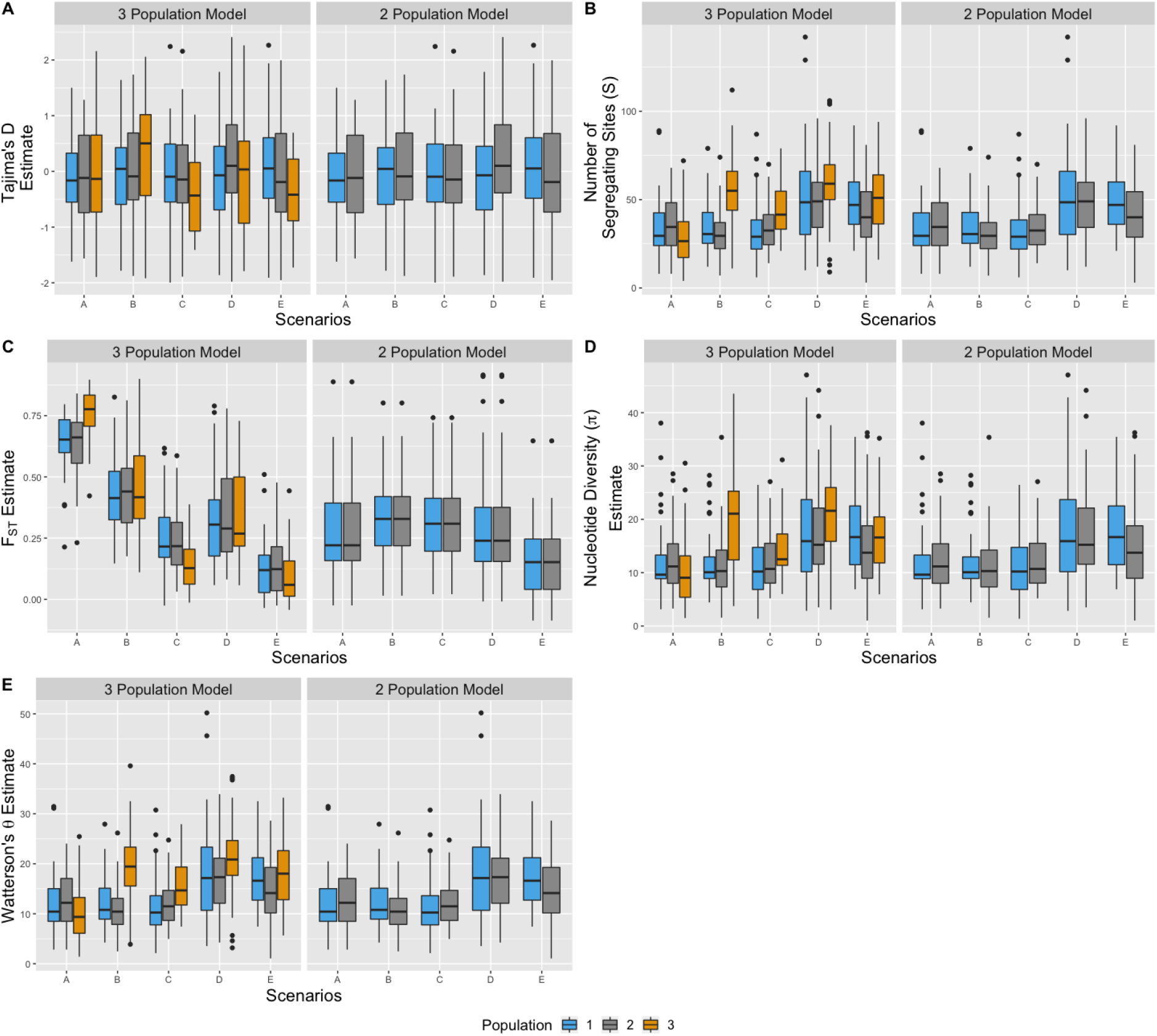
Summary Statistic Estimates under models A-E while using 5 genomic loci. Under each model, we estimate (A) Tajima’s D, (B) Number of segregating sites, (C) Genetic Differentiation (*F*_*st*_, (D) Nucleotide Diversity (*π*), and (E) Effective population size (Watterson’s *θ*) under either a 3 population sampling strategy (left panels), or a 2 population sampling strategy (right panels). We would expect that all estimates of Tajima’s D would be close to 0 (indicating putatively neutral evolution), estimates of differentiation between the two sampled populations to be smaller with greater gene flow from the ‘ghost’ (i.e. under models D, E, compared to model A), and estimates of genetic diversity to be greater with greater gene flow from the ‘ghost’. This is reflected in all estimated summary statistics under the 2 population sampling strategy. The 3 population sampling strategy serves as a positive control, wherein in an ideal scenario, we would sample all three populations prior to estimating summary statistics.

### Effects on the estimation of divergence times

Divergence times (*t*; 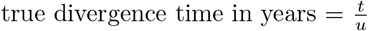; *u* is the mutation rate per locus per generation) between sampled populations were consistently under-estimated when migration increased from the unsampled ‘ghost’ population (Figs. 3, Supplementary Figs. 2, 3, 32), compared to the true value (*t*_0_ = 5) across all our simulations. Nonetheless, across a majority of our simulations, the simulated divergence time was included in the estimated 95% confidence interval. However, sampling more genomic loci led to a reduction in the confidence intervals around divergence time estimates (Supplementary Fig. 32. The 2-population model with high bi-directional ‘ghost’ migration (model E) was the only model to underestimate divergence times without the confidence interval including the true value (simulated divergence time) while sampling either 2 or 5 loci.

**Figure 3:**
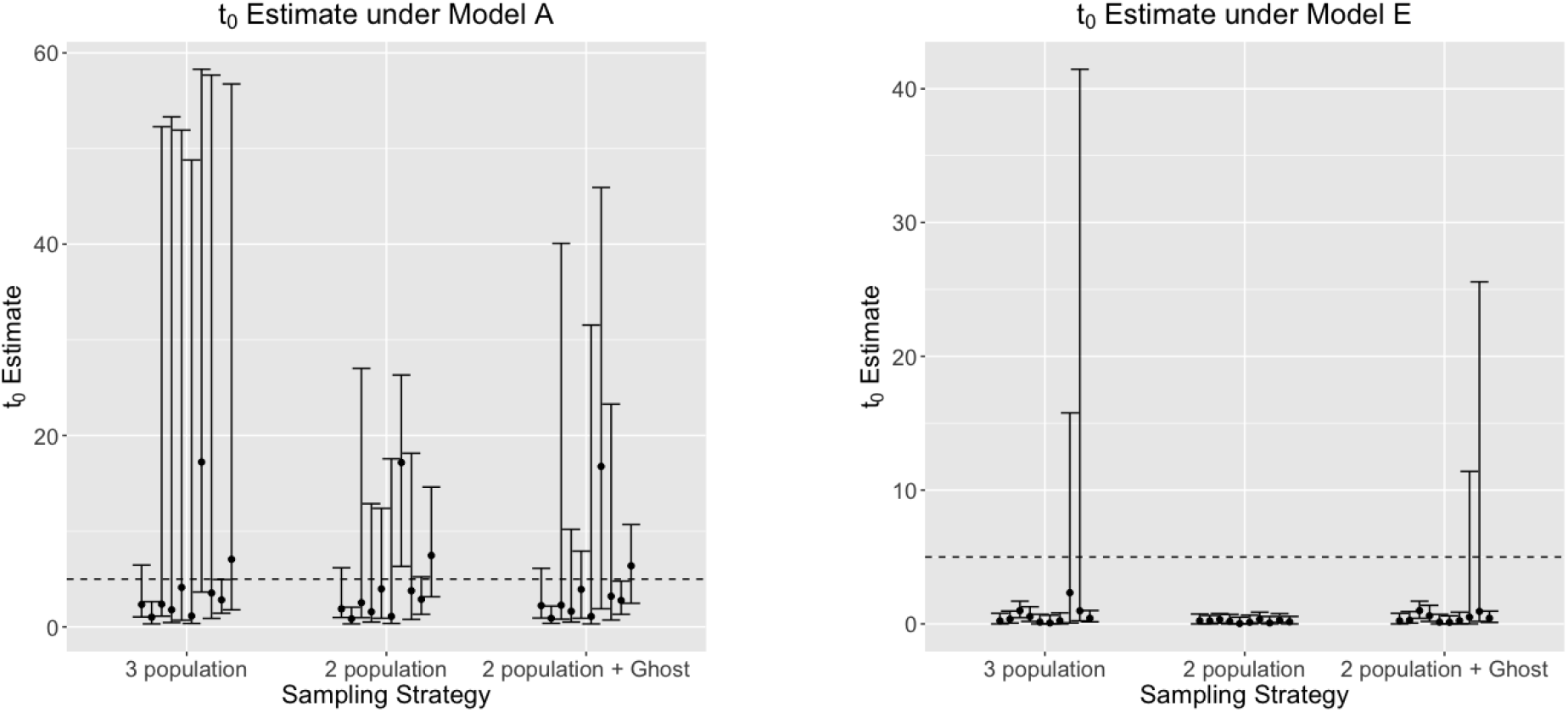
Divergence time (*t*_0_) estimates between sampled populations, using 5 genomic loci under models A and E, estimated using a 3-population model, 2-population model (without a ‘ghost’), and a 2-population model with a ‘ghost’ outgroup. The dotted line indicates the true (simulated) divergence time (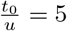, where *u* is the mutation rate per site per generation. Modes, and 95% confidence intervals from 10 replicate runs are shown per model. Compared to the null model A, divergence times are consistently under-estimated with large unidirectional, or bidirectional gene flow from the ‘ghost’.

Divergence time estimates between the two sampled populations and the ‘ghost’ population were consistent with the true simulated divergence time (*t*_1_ = 20) except for the model E (Supplementary Figs. 4, 5) Including a ‘ghost’ population in model led to underestimation in the no migration model (A), but led to better estimates with smaller confidence interval widths in most other models, reiterating the importance of including a ‘ghost’ population while estimating evolutionary history using `IMa2p`.

### Effects on the estimation of effective population sizes

While estimating scaled effective population sizes (*θ* = 4*N*_*e*_*u*; *N*_*e*_ is the effective population size; *u* is the mutation rate per locus per generation) of sampled populations, the inclusion of a ‘ghost’ population resulted in estimates similar to having sampled all three populations, while the two population models had wider confidence intervals across all estimates. Nonetheless, greater migration rates (under models C and E) resulted in overestimation of effective population sizes in both sampled populations (Figs. 4, Supplementary Figs. 6-15, 33-37).

**Figure 4:**
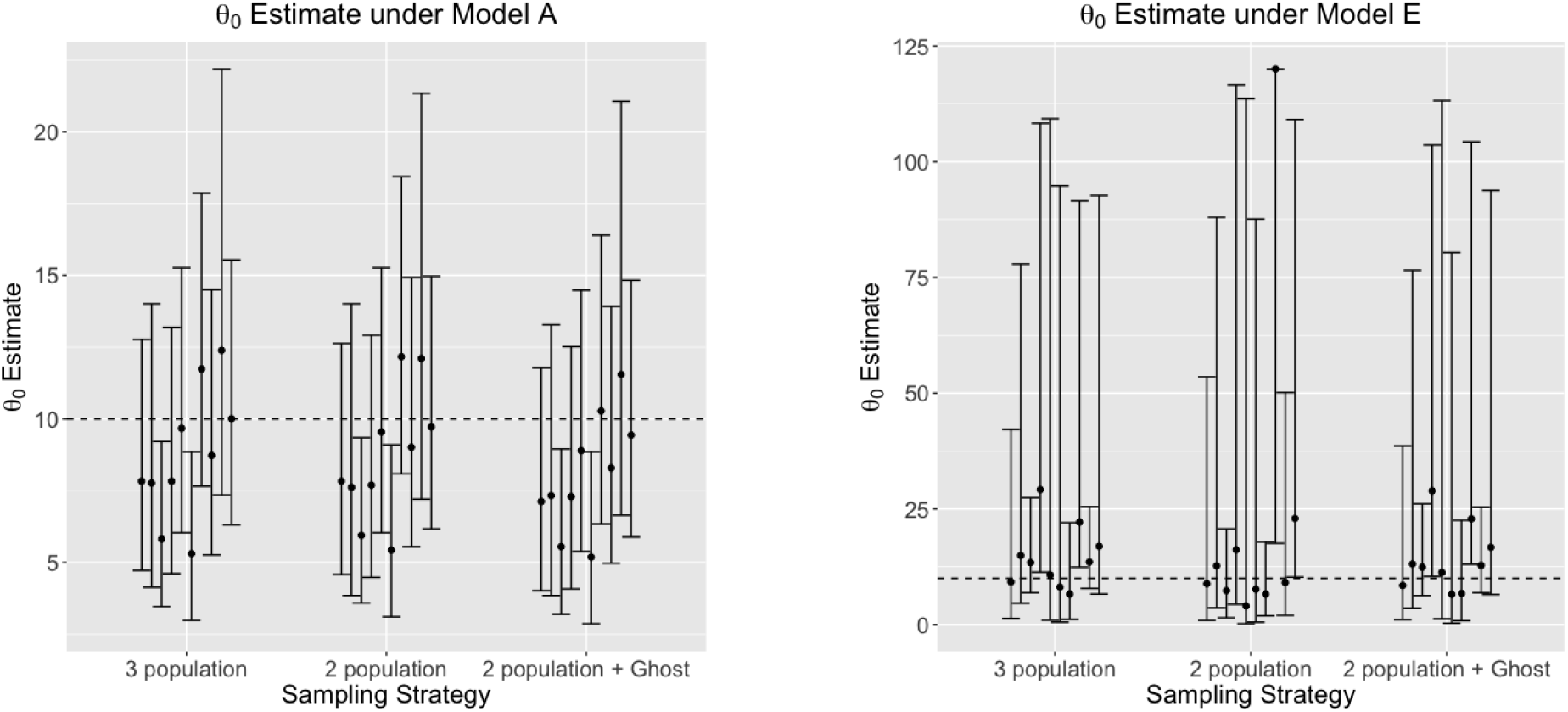
Scaled effective population size (*θ*_0_ = 4*N*_0_*u*) estimate of the first sampled populations, using 5 genomic loci under models A and E, estimated using a 3-population model, 2-population model (without a ‘ghost’), and a 2-population model with a ‘ghost’ outgroup. True simulated *θ*_0_ = 10, and is shown with the dotted line. *θ*_0_ is consistently over-estimated, as seen by the inflated confidence intervals around the mode, with increased bi-directional gene flow from the ‘ghost’ (models E), compared to the null model of no gene flow from the ‘ghost’ (A).

The level of gene flow from the ‘ghost’ population greatly affects effective population size estimates of the common ancestor of the two sampled populations. Estimates from the two population model have smaller confidence intervals when there is no or low migration with the ‘ghost’ (models A, B), but in models with high migration (C and E), the effective population size was consistently overestimated by ≈ 2 − 10 times the ‘true’ value (*θ* = 10).

When estimating the effective population sizes of the common ancestor of the two sampled populations and the ‘ghost’ population, the two population model which included the ‘ghost’ performed as well as the three population model in models of low or no migration (A and B). However in high migration models (C, E), the three population model obtained better estimates, while the model with the inclusion of a ‘ghost’ mostly underestimated *θ*_4_. Increasing the number of loci did not affect estimates of effective population sizes in any of our models.

### Effects on the estimation of migration rates between sampled populations

Migration rates (scaled as *m* = *M/u*; *M* is the population migration rate per locus per generation; *u* is the mutation rate per generation per locus) between sampled populations were relatively accurately estimated under both 2 and 5 locus sampling schemes (Figs. 5, Supplementary Figs. 16-31, 38, 39). The inclusion of a ‘ghost’ population did not affect these estimates in two and three population models. Regardless of direction, models with highest true migration rates (C and E) have consistent overestimates of migration rates between sampled populations for all models (Supplementary Figs. 40-45). Estimates of migration rates in models with lower true migration rates (B and D) are more accurate. The model with no migration to or from ‘ghost’ (A) has greatest under-estimate of any model. Estimates of bidirectional migration rates between the ‘ghost’ and sampled populations were greatly under-estimated when ‘true’ migration was high, with estimates approximating ten times below the ‘true’ value. Additionally, increasing the number of sampled loci did not improve estimates of migration rates.

**Figure 5:**
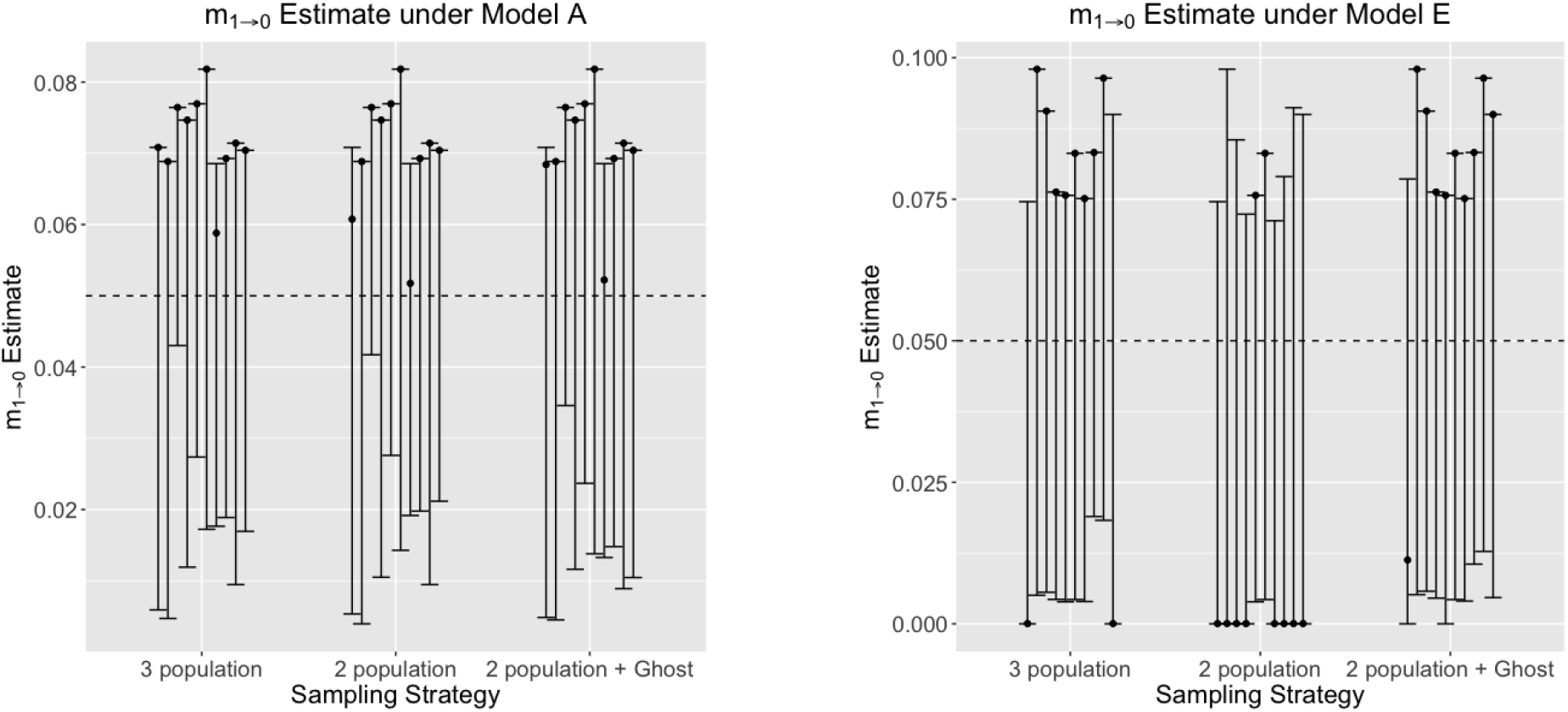
Scaled migration rate (*m*_1→0_ = 4*N*_1_*u*, where *N*_1_ is the effective population size of one of the sampled populations, and *u* is the mutation rate per site per generation) estimate between the two sampled populations, using 5 genomic loci under models A and E, estimated using a 3-population model, 2-population model (without a ‘ghost’), and a 2-population model with a ghost outgroup. True simulated *m*_1→0_ = 0.05 and is shown with a dotted line. Models with increased gene flow from the ghost consistently lead to over-estimation of *m*_1→0_, as observed by the inflated confidence intervals around the mode.

### Effect of ‘ghost’ gene flow on African hunter-gatherer Evolutionary History

The Hadza was estimated to have the smallest size across models (≈ 5, 000), while Yoruba was estimated to have the largest population size (≈ 25, 000; Fig. 6C-H, Supplementary Tables 2,5). All ancestral population sizes were estimated to be around 15, 000, consistent with previous estimates (Hey et al., 2018). Importantly, no significant migration was detected in all pairwise models by using the LLR test, indicating that there is insufficient information in the data provided to the pairwise model to glean migration estimates (Supplementary Table 1). Divergence time estimates indicated that the earliest split occurred between the Baka and Sandawe populations (≈ 67, 000 ybp), and the most recent split occurred between the Hadza and Sandawe (≈ 24, 000 ybp, Fig. 6C-H, Supplementary Tables 5,6).

**Figure 6:**
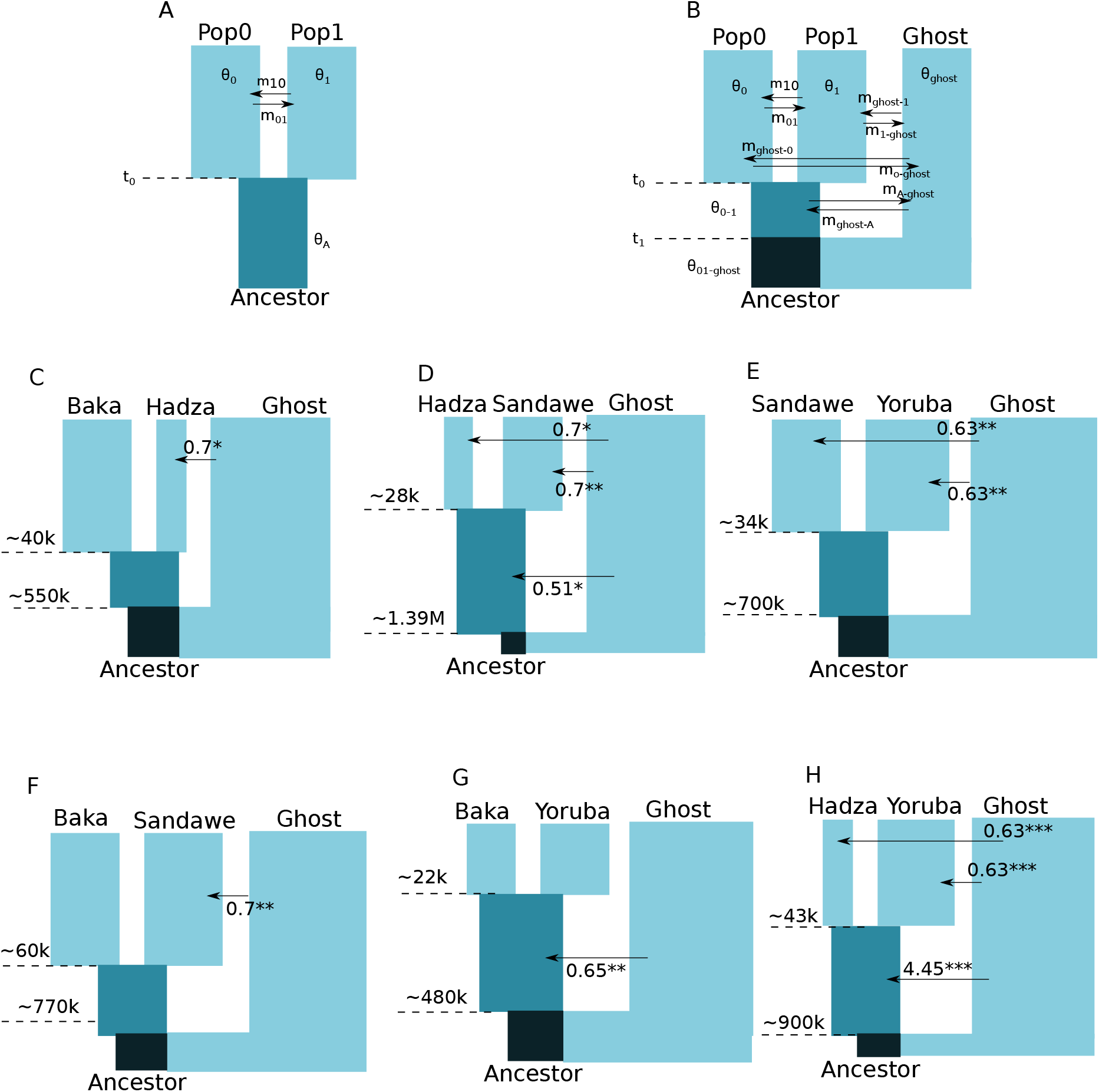
(A) 2-population IM model, showing all the estimated parameters - Effective population sizes in demographic scales, *θ*_0_, *θ*_1_, *θ*_*A*_, migration rates (2*N*_*e*_*m*), *m*_10_, *m*_01_, and divergence time *t*_0_ in years before present. (B) 2-population IM model with an outgroup unsampled ‘ghost’ population, with all estimated parameters (effective population sizes, divergence times, and migration rates), (C-H) Estimated divergence times in years before present under the 2-population with ‘ghost’ models between all pairs of populations (Hadza, Sandawe, Baka, and Yoruba). Width of population blocks is proportional to their estimated sizes, while only statistically significant migration rates > 0 are shown (∗*p* < 0.05, ∗∗*p* < 0.01, ∗∗∗*p* < 0.001). Statistical significance of migration rates were estimated using the LLR test of (Nielsen & Wakeley, 2001). All parameters were scaled by a human generation time of 29 years.

However, with the addition of an unsampled ‘ghost’ population to these models, significant migration (assessed using the LLR test of (Nielsen & Wakeley, 2001)) was detected into the extant Hadza, Sandawe, and Yoruba populations (Fig. 6C-H, Supplementary Tables 3,4). Common ancestral populations of Hadza-Sandawe, Baka-Yoruba, and Hadza-Yoruba were also estimated to have significant migration from unsampled ‘ghost’ populations.

Divergence times of common ancestral populations of all tested pairs with the unsampled ‘ghost’ population was several folds larger than divergence times estimated under the 2-population models (≈ 483000 ybp - ≈ 1393800 ybp). ‘Ghost’ populations were also considerably larger than sampled populations (≈ 250000), except between Baka and Sandawe (≈ 92, 000, Fig. 6C-H, Supplementary Table 5).

All two population and two populations with ‘ghost’ models converged within 48 hours, with 100, 000 steps of burn-in, which was assessed by similarity of parameter estimates across duplicate runs, as well as using `Tracer` (Rambaut et al., 2018).

## Discussion

Gene flow from unsampled ‘ghost’ populations has affected species’ population histories, yet its effect on current evolutionary history models and summary statistics has scarcely been quantified. To quantify these effects, we simulate genomic data under the Isolation with Migration (IM) model with different levels of gene flow from unsampled ‘ghost’ populations. Our models reveal that with increased gene flow from unsampled ‘ghosts’, differentiation between sampled populations (*F*_*st*_) is underestimated (Fig. 2C) and genomic diversity (Watterson’s *θ*, nucleotide diversity, *π*) is overestimated. Sampling more genomic loci may improve precision, but it will not eliminate errors or bias. We acknowledge that our simulations have caveats, including sampling a fixed number of individuals from two populations of constant size, primarily owing to the enormity of simulation space. While this limits the generality of our conclusions, we contend that the effects of gene flow from unsampled ‘ghost’ populations are far from negligible. While we only demonstrate these biases using the IM suite of tools, we contend that similar biases are bound to affect estimates using other tools alike, if ‘ghost’ populations are not accounted for.

For instance, consider an allopatric speciation scenario (where geographic separation makes two species genomically distinct). If there is more gene flow from an unsampled ‘ghost’ population, differentiation between the sampled extant populations would be lower than expected under a scenario of no gene flow from the ‘ghost’. Investigators may then wrongly conclude that the species are not genetically distinct and that there is gene flow between them when in fact the gene flow occurred in the past between the extant populations and the ‘ghost’.

Similarly, not accounting for an unsampled ‘ghost’ in a model-based estimation of evolutionary history can lead to incorrect conclusions about the true evolutionary history. Our simulations consistently show that when there is an unknown ‘ghost’ population, current statistical methods will underestimate divergence times, overestimate effect population sizes, and underestimate migration rates between sampled populations.

This affirms previous findings which also showed that not accounting for an unsampled ‘ghost’ can skew estimates of evolutionary history (Beerli, 2004).

Estimates of unidirectional migration rate from one sampled population into the unsampled ‘ghost’ were always over-estimated when there was no gene flow into the ‘ghost’, while under-estimated when there was a larger degree of gene flow into the ‘ghost’ (e.g. Supplementary Figs. 20, 21). This could due to the fact that the divergence time between the common ancestor of the sampled populations, and the unsampled ‘ghost’ population is as yet too small, leading to erroneous conclusions of migration rates between recently diverged populations (Hey, Chung, & Sethuraman, 2015).

Increasing the number of sampled genomic loci generally improved the confidence intervals around all estimates. This could be an ideal strategy, especially in this age of next generation sequencing, and the ability to obtain long haplotypic segments at lower costs.

Estimates of effective population sizes were largely robust to the evolutionary model. For instance, in a model with no migration from an unsampled ‘ghost’ population, the effective population size of the common ancestor of the sampled populations is accurately estimated (Fig. 4).

Across all our simulations, including an unsampled ‘ghost’ population in the model improved all estimated evolutionary parameters. This should be an ideal strategy across all population genetics studies, where the true population tree or history is unknown.

For instance, in a recent study (Hey et al., 2018) of African hunter-gatherers (Hadza, Sandawe) and agriculturalists (Yoruba, Baka), significant unidirectional migration (‘ghost’ into Baka, Yoruba, Sandawe, and the four populations’ common ancestor) was only detected when a ‘ghost’ population was included in their model. Including the ‘ghost’ population also produced more accurate estimates of the small effective population sizes in the sampled populations. Both of these patterns also hold in our study.

Our analyses of the demographic history of 20 diploid African Hunter-Gatherer genomes across 355 loci revealed some interesting patterns of population size change, and ancestral introgression.The Hadza-Sandawe split (Figure 6D) was determined to be very recent (as recent as 4500 ybp, 95% c.i of 4505 − 49, 560). This concurs with the prior neighbor-joining analyses of (Lachance et al., 2012), which indicated the Baka divergence before the Hadza-Sandawe split. Also in agreement with (Lachance et al., 2012), all our models determined that the Hadza underwent a recent population bottleneck (with *N*_*e*_ = 128 − 4300), although estimates are lower than those determined from polymorphism levels (*N*_*e*_ = 9200 − 20, 900 (Lachance et al., 2012)). Significant ancestral introgression with an unsampled ‘ghost’ population was also detected, but was largely unidirectional (from the ‘ghost’ into extant populations), suggesting that introgression events were more ancient than divergence from recent common ancestors of extant populations. Divergence times with this ancestral ‘ghost’ population were as high as 1.7 million years ago, which was also previously detected by (Lachance et al., 2012) using TMRCA estimates on introgressed regions. Importantly, if a researcher were to only utilize 2-population IM models to infer the evolutionary history of the Hadza, Sandawe, Yoruba, and Baka, their inference would be at odds with the estimated population structure and admixture proportions of (Lachance et al., 2012). The addition of an unsampled ‘ghost’ however to the 2-population IM model in `IMa2p` correctly infers their evolutionary history, congruent with the findings of (Hey et al., 2018).

Our study re-iterates the key points made by (Beerli, 2004), in that while it is impossible to always sample all species or large number of informative genomic loci, it is imperative to account for the presence of unsampled ‘ghost’ populations while estimating both summary statistics and the evolutionary history of sampled species.

Detecting the presence of a ‘ghost’ population in genomic data should ideally be a multi-pronged approach: (a) computing summary statistics from population genetic data, and reconciling observed summary statistics with the species’ known biological history, (b) estimating the population genetic structure of sampled populations via a model-based approach like `STRUCTURE` or `ADMIXTURE` (Pritchard et al., 2000; Alexander et al., 2009), (c) if available, using genome-wide estimates of summary statistics like differentiation (e.g. *F*_*st*_) and divergence (*D*_*xy*_) between populations to identify regions of low differentiation or divergence, and hence putatively introgressed from an unsampled ‘ghost’, (d) computing TMRCA (Time to Most Recent Common Ancestry) distribution across the genome, `sensu` (Lachance et al., 2012), to visualize differences in coalescent time distributions of putatively introgressed (from archaic/‘ghost’ populations) regions, versus non-introgressed regions, and finally, (e) testing a variety of evolutionary models, both including and excluding the presence of an unsampled ‘ghost’ under a model-based framework, like ABC (Approximate Bayesian Computation (Beaumont, Zhang, & Balding, 2002)), or IM (Hey & Nielsen, 2004) to find the best fitting model that explains observed population genomic data.

Alternately, in the absence of genome-size data, for e.g. performing analyses with a few microsatellite or nuclear loci), our recommendation would still be to begin with steps (a) and (b) above, and then, estimate evolutionary history using a variety of models with and without ‘ghost’ populations using the LLR tests implemented in `IMa2p` and `IMa3`. Additionally, both programs allow for conveniently testing of significant non-zero bidirectional migration estimates from an outgroup ‘ghost’ using the LLR test of (Nielsen & Wakeley, 2001).

## Acknowledgments

This work was supported by an NSF ABI Grant 1564659 to Arun Sethuraman and Jody Hey. This research includes calculations carried out on Temple University’s HPC resources and thus was supported in part by the National Science Foundation through major research instrumentation grant number 1625061 and by the US Army Research Laboratory under contract number W911NF-16-2-0189. We would like to thank two anonymous reviewers, Robin Waples, Jody Hey, Roxane Saisho, and Dylan Steinecke for their helpful suggestions and constructive feedback. We graciously acknowledge the Joe Lachance and Sarah Tishkoff for providing us access to the African hunter-gatherer genomic data.

## Data Accessibility

All simulation scripts, `IMa2p` datasets will be made available on the Author’s GitHub page at www.github.com/arunsethuraman.

## Author Contributions

ML conceptualized the study, performed all the simulations, ran all the IM analyses, and AS and ML wrote the paper.

## References

Alexander, D. H., Novembre, J., & Lange, K. (2009). Fast model-based estimation of ancestry in unrelated individuals. Genome research, 19 (9), 1655–1664.

Beaumont, M. A., Zhang, W., & Balding, D. J. (2002). Approximate bayesian computation in population genetics. Genetics, 162 (4), 2025–2035.

Beerli, P. (2004). Effect of unsampled populations on the estimation of population sizes and migration rates between sampled populations. Molecular Ecology, 13 (4), 827–836.

Beerli, P., & Felsenstein, J. (2001). Maximum likelihood estimation of a migration matrix and effective population sizes in n subpopulations by using a coalescent approach. Proceedings of the National Academy of Sciences, 98 (8), 4563–4568.

Chung, Y., & Hey, J. (2017, 02). Bayesian Analysis of Evolutionary Divergence with Genomic Data under Diverse Demographic Models. Molecular Biology and Evolution, 34 (6), 1517–1528. doi: 10.1093/molbev/msx070

Durvasula, A., & Sankararaman, S. (2019). Recovering signals of ghost archaic introgression in african populations. bioRxiv, 285734.

Gronau, I., Hubisz, M. J., Gulko, B., Danko, C. G., & Siepel, A. (2011). Bayesian inference of ancient human demography from individual genome sequences. Nature genetics, 43 (10), 1031.

Gutenkunst, R. N., Hernandez, R. D., Williamson, S. H., & Bustamante, C. D. (2009). Inferring the joint demographic history of multiple populations from multidimensional snp frequency data. PLoS genetics, 5 (10), e1000695.

Hey, J. (2011a). Documentation for ima2. Department of Genetics, Rutgers University: Piscataway, NJ, USA.

Hey, J. (2011b). Documentation for ima2. New Brunswick, NJ: Rutgers Uni.

Hey, J., Chung, Y., & Sethuraman, A. (2015). On the occurrence of false positives in tests of migration under an isolation-with-migration model. Molecular ecology, 24 (20), 5078–5083.

Hey, J., Chung, Y., Sethuraman, A., Lachance, J., Tishkoff, S., Sousa, V. C., & Wang, Y. (2018). Phylogeny estimation by integration over isolation with migration models. Molecular biology and evolution, 35 (11), 2805–2818.

Hey, J., & Nielsen, R. (2004). Multilocus methods for estimating population sizes, migration rates and divergence time, with applications to the divergence of drosophila pseudoobscura and d. persimilis. Genetics, 167 (2), 747–760.

Hey, J., & Nielsen, R. (2007). Integration within the felsenstein equation for improved markov chain monte carlo methods in population genetics. Proceedings of the National Academy of Sciences, 104 (8), 2785–2790.

Hudson, R. R. (2002). Generating samples under a wright–fisher neutral model of genetic variation. Bioinformatics, 18 (2), 337–338.

Hudson, R. R., & Kaplan, N. L. (1985). Statistical properties of the number of recombination events in the history of a sample of dna sequences. Genetics, 111 (1), 147–164.

Konečný, A., Estoup, A., Duplantier, J.-M., Bryja, J., Bâ, K., Galan, M., … Cosson, J.-F. (2013). Invasion genetics of the introduced black rat (rattus rattus) in senegal, west africa. Molecular Ecology, 22 (2), 286–300.

Kuhlwilm, M., Han, S., Sousa, V. C., Excoffier, L., & Marques-Bonet, T. (2019). Ancient admixture from an extinct ape lineage into bonobos. Nature ecology & evolution, 3 (6), 957.

Lachance, J., Vernot, B., Elbers, C. C., Ferwerda, B., Froment, A., Bodo, J.-M., … others (2012). Evolutionary history and adaptation from high-coverage whole-genome sequences of diverse african hunter-gatherers. Cell, 150 (3), 457–469.

Lawson, D. J., Van Dorp, L., & Falush, D. (2018). A tutorial on how not to over-interpret structure and admixture bar plots. Nature Communications, 9 (1), 3258.

Nielsen, R., Akey, J. M., Jakobsson, M., Pritchard, J. K., Tishkoff, S., & Willerslev, E. (2017). Tracing the peopling of the world through genomics. Nature, 541 (7637), 302.

Nielsen, R., & Wakeley, J. (2001). Distinguishing migration from isolation: a markov chain monte carlo approach. Genetics, 158 (2), 885–896.

Pfeifer, B., Wittelsbuerger, U., Ramos-Onsins, S. E., & Lercher, M. J. (2014). Popgenome: An efficient swiss army knife for population genomic analyses in r. Molecular Biology and Evolution, 31, 1929–1936. doi: 10.1093/molbev/msu136

Pritchard, J. K., Stephens, M., & Donnelly, P. (2000). Inference of population structure using multilocus genotype data. Genetics, 155 (2), 945–959.

Rambaut, A., Drummond, A. J., Xie, D., Baele, G., & Suchard, M. A. (2018). Posterior summarization in bayesian phylogenetics using tracer 1.7. Systematic biology, 67 (5), 901.

Rosenthal, D. M., Ramakrishnan, A. P., & Cruzan, M. B. (2008). Evidence for multiple sources of invasion and intraspecific hybridization in brachypodium sylvaticum (hudson) beauv. in north america. Molecular Ecology, 17 (21), 4657–4669.

Scheet, P., & Stephens, M. (2006). A fast and flexible statistical model for large-scale population genotype data: applications to inferring missing genotypes and haplotypic phase. The American Journal of Human Genetics, 78 (4), 629–644.

Sethuraman, A., & Hey, J. (2016). Im a2p–parallel mcmc and inference of ancient demography under the isolation with migration (im) model. Molecular ecology resources, 16 (1), 206–215.

Slatkin, M. (1985). Gene flow in natural populations. Annual review of ecology and systematics, 16 (1), 393–430.

Slatkin, M. (2005). Seeing ghosts: the effect of unsampled populations on migration rates estimated for sampled populations. Molecular Ecology, 14 (1), 67–73.

