## Supplemental Tables and Figures for "Accounting for Gene Flow from Unsampled ‘Ghost’ Populations while Estimating Evolutionay History under the Isolation with Migration Model"

### Supplementary Material - The Effects of Gene Flow from Unsampled 'Ghost' Populations on the Estimation of Evolutionary History under the Isolation with Migration Model

Arun Sethuraman*[1] and Melissa Lynch†[1]

[1]Department of Biological Sciences, California State University San Marcos

May 22, 2020

#### Supplementary Tables and Figures

| Pop 1 | Pop 2 | Genealogies | $M_{2\to1}$ | $M_{1\to2}$ |
|---|---|---|---|---|
| Baka | Hadza | 24055 | 0.693(ns) | 0.700(ns) |
| Hadza | Sandawe | 18919 | 0.700(ns) | 0.700(ns) |
| Sandawe | Yoruba | 14805 | 0.163(ns) | 0.001(ns) |
| Baka | Sandawe | 23164 | 0.700(ns) | 0.419(ns) |
| Baka | Yoruba | 18079 | 0.000(ns) | 0.551(ns) |
| Hadza | Yoruba | 18919 | 0.000(ns) | 0.005(ns) |

**Table 1:** African populations, number of genealogies sampled over parallel MCMC runs, bidirectional migration estimates under the 2 population IM model. Significance of the LLR test for migration are indicated in parentheses. *ns* indicates not significant ($p > 0.05$.

*
†

| Pop 1 | Pop 2 | $\Theta_1$ | $\Theta_2$ | $\Theta_A$ | $t$ |
|---|---|---|---|---|---|
| Baka | Hadza | 15952(11081-31052) | 5723(4262-7672) | 13760(12055-15465) | 52264(38139-72040) |
| Hadza | Sandawe | 3707(2272-5620) | 14708(8968-47232) | 14947(13273-16382) | 23580(15258-37450) |
| Sandawe | Yoruba | 15700(10951-25727) | 24936(17811-44198) | 15700(14381-17020) | 38309(26050-50568) |
| Baka | Sandawe | 19326(13664-34590) | 15633(11940-22034) | 14156(12679-15879) | 67112(49977-87102) |
| Baka | Yoruba | 14318(9326-28242) | 21674(15369-35335) | 15106(13267-16682) | 41167(28970-56415) |
| Hadza | Yoruba | 5754(4416-7627) | 28768(20740-51247) | 15120(14050-16726) | 45068(32636-57501) |

**Table 2:** Demographic scale effective population sizes of extant sampled ($\theta_1$, $\theta_2$) and ancestors ($\theta_A$), divergence times ($t$) in years under the 2 population IM model. Also shown are 95% confidence intervals in parentheses. All estimates were scaled by a human generation time of 29 years.

| Pop 1 | Pop 2 | Genealogies | $M_{2\to 1}$ | $M_{1\to 2}$ | $M_{ghost\to 1}$ | $M_{1\to ghost}$ |
|---|---|---|---|---|---|---|
| Baka | Hadza | 19490 | 0.700(ns) | 0.301(ns) | 0.232(ns) | 0.120(ns) |
| Hadza | Sandawe | 19019 | 0.700(ns) | 0.000(ns) | 0.700(*) | 0.000(ns) |
| Sandawe | Yoruba | 13524 | 0.286(ns) | 0.054(ns) | 0.63(**) | 0.000(ns) |
| Baka | Sandawe | 19340 | 0.479(ns) | 0.297(ns) | 0.283(ns) | 0.000(ns) |
| Baka | Yoruba | 15836 | 0.318(ns) | 0.446(ns) | 0.65(ns) | 0.000(ns) |
| Hadza | Yoruba | 13334 | 0.306(ns) | 0.63(ns) | 0.63(***) | 0.000(ns) |

**Table 3:** Number of genealogies sampled over the parallel MCMC run, bidirectional migration estimates under the 2 population with an unsampled 'ghost'. Also indicated are significance of migration estimates using the LLR test. *ns* indicates not significant, $*$ indicates significant at $p < 0.05$, $**$ at $p < 0.01$, $***$ at $p < 0.001$.

| Pop 1 | Pop 2 | $M_{ghost\to 2}$ | $M_{2\to ghost}$ | $M_{A\to ghost}$ | $M_{ghost\to A}$ |
|---|---|---|---|---|---|
| Baka | Hadza | 0.700(*) | 0.000(ns) | 0.008(ns) | 0.700(ns) |
| Hadza | Sandawe | 0.700(**) | 0.000(ns) | 0.000(ns) | 0.509(*) |
| Sandawe | Yoruba | 0.63(**) | 0.000(ns) | 0.012(ns) | na(ns) |
| Baka | Sandawe | 0.700(**) | 0.000(ns) | 0.000(ns) | 0.013(ns) |
| Baka | Yoruba | 0.65(ns) | 0.000(ns) | 0.042(ns) | 0.65(**) |
| Hadza | Yoruba | 0.63(***) | 0.000(ns) | 0.003(ns) | 4.451(***) |

**Table 4:** Bidirectional migration estimates under the 2 population IM model, with an unsampled ghost population. Also indicated are significance of migration estimates using the LLR test. *ns* indicates not significant, $*$ indicates significant at $p < 0.05$, $**$ at $p < 0.01$, $***$ at $p < 0.001$.

| Pop 1 | Pop 2 | $\Theta_{A1}$ | $\Theta_{A2}$ | $t_1$ | $t_2$ |
|---|---|---|---|---|---|
| Baka | Hadza | 15358(12164-22730) | 10198(3072-12901) | 41332(21379-61286) | 557272(272222-1534992) |
| Hadza | Sandawe | 11493(9868-14279) | 3134(580.5-12190) | 28280(17507-39053) | 1393804(677375-2180259) |
| Sandawe | Yoruba | 14103(11774-20572) | 10739(2458-13327) | 34564(19536-46586) | 692774(377194-1852908) |
| Baka | Sandawe | 12574(10419-20478) | 10658(838.3-13293) | 59733(37507-76402) | 768192(293108-2129544) |
| Baka | Yoruba | 17398(13789-23326) | 10439(7861-12501) | 22438(10471-43381) | 483171(315632-821241) |
| Hadza | Yoruba | 12858(10780-16235) | 8442(2469-11819) | 43745(31677-55813) | 924680(601872-1657788) |

**Table 5:** Ancestral effective population sizes, and divergence times between population 1 and 2 ($t_1$), and the unsampled 'ghost' and their common ancestor ($t_2$ in years before present, estimated using 2-population with ghost models.

| Pop 1 | Pop 2 | $\Theta_1$ | $\Theta_2$ | $\Theta_{ghost}$ |
|---|---|---|---|---|
| Baka | Hadza | 12409(6758-26416) | 4546(2580-6512) | 245610(10689-239958) |
| Hadza | Sandawe | 4295(2670-6153) | 17066(10797-54679) | 232068(26121-227889) |
| Sandawe | Yoruba | 14103(7893-22643) | 23419(13844-45932) | 258644(29888-253727) |
| Baka | Sandawe | 17604(10898-33651) | 13532(9221-19041) | 92330(13772-233638) |
| Baka | Yoruba | 8377(3480-19202) | 13532(6057-25903) | 257615(17140-252202) |
| Hadza | Yoruba | 5585(4026-7143) | 27923(19092-544148) | 259623(34677-254947) |

**Table 6:** Demographic scale effective population size ($N_e$) estimates under the 2 population IM model, with an unsampled ghost population. Also indicated are 95% confidence intervals in parentheses. All estimates were scaled with a human generation time of 29 years.

| Model | Theta | m | t |
|---|---|---|---|
| BH | 14.20 | 0.70 | 1.42 |
| HS | 14.20 | 0.70 | 1.42 |
| SY | 15.98 | 0.63 | 1.60 |
| BS | 14.20 | 0.70 | 1.42 |
| BY | 15.39 | 0.65 | 1.54 |
| HY | 15.98 | 0.63 | 1.60 |
| All | 5.51 | 20.00 | 2.20 |

**Table 7:** Parameter prior upper limits on population sizes ($\Theta$), migration rates ($m$), and divergence times ($t$) used in `IMa2-MPI` analyses of African Hunter-Gatherer demographic history. Populations studied in the model are Baka (B), Hadza (H), Sandawe (S), and Yoruba (Y). Limits are shown for each pairwise 2 population (with and without a "ghost"), and the full 4 population (All populations with and without "ghost") models.

**Figure 1:** Summary Statistic Estimates under models A-E while using 2 genomic loci. Under each model, we estimate (A) Tajima's D, (B) Number of segregating sites, (C) Genetic Differentiation ($F_{st}$, (D) Nucleotide Diversity ($\pi$), and (E) Effective population size (Watterson's $\theta$) under either a 3 population sampling strategy (left panels), or a 2 population sampling strategy (right panels). We would expect that all estimates of Tajima's D would be close to 0 (indicating putatively neutral evolution), estimates of differentiation between the two sampled populations to be smaller with greater gene flow from the 'ghost' (i.e. under models D, E, compared to model A), and estimates of genetic diversity to be greater with greater gene flow from the 'ghost'. This is reflected in all estimated summary statistics under the 2 population sampling strategy. The 3 population sampling strategy serves as a positive control, wherein in an ideal scenario, we would sample all three populations prior to estimating summary statistics.

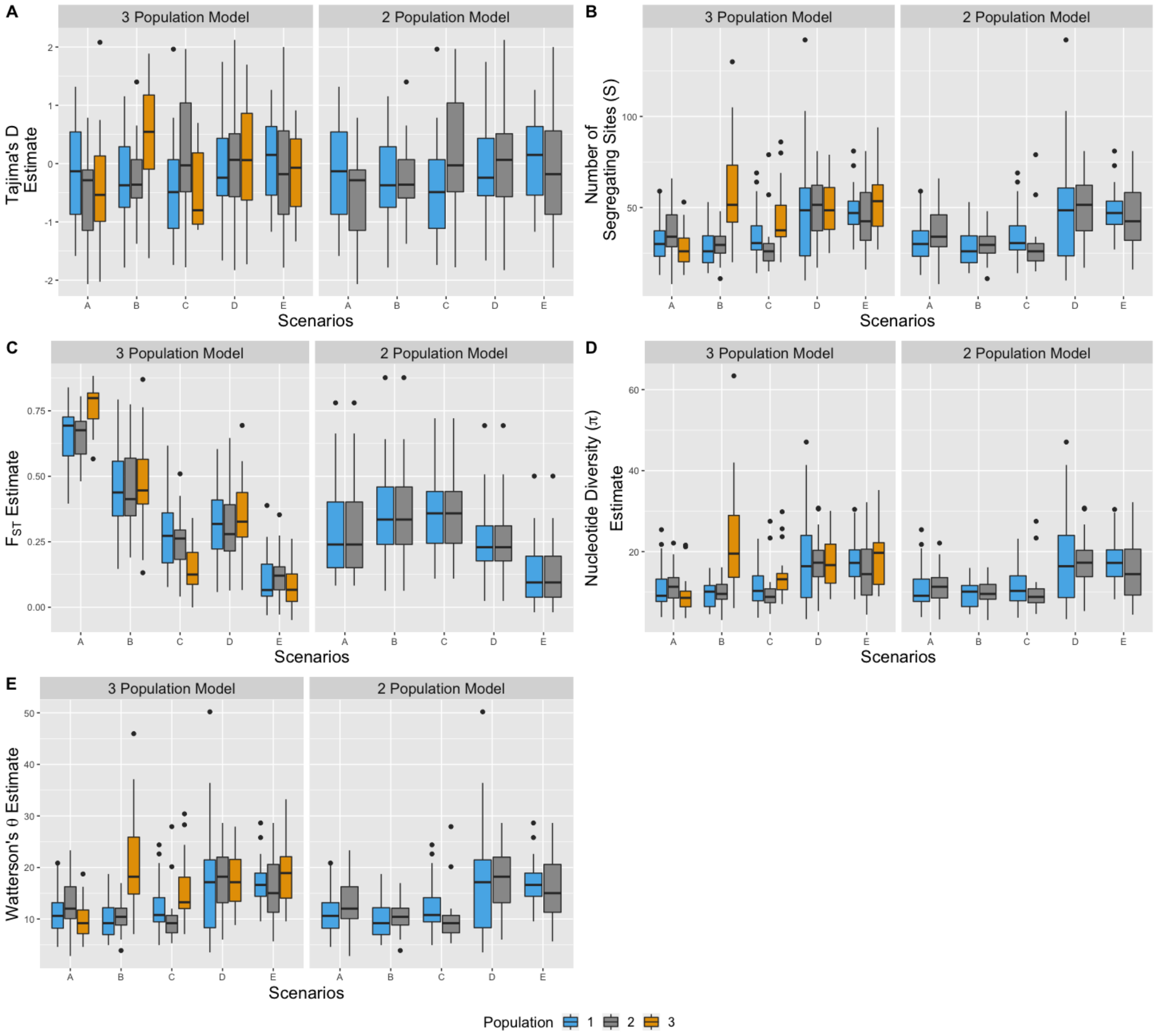

**Figure 2:** Divergence time ($t_0$) estimates between sampled populations, using 2 genomic loci under models A-E, estimated using a 3-population model, 2-population model (without a 'ghost'), and a 2-population model with a 'ghost' outgroup. The dotted line indicates the true (simulated) divergence time ($\frac{t_0}{u} = 5$, where $u$ is the mutation rate per site per generation. Modes, and 95% confidence intervals from 10 replicate runs are shown per model. Compared to the null model A, divergence times are consistently under-estimated with large unidirectional, or bidirectional gene flow from the 'ghost' (models C and E).

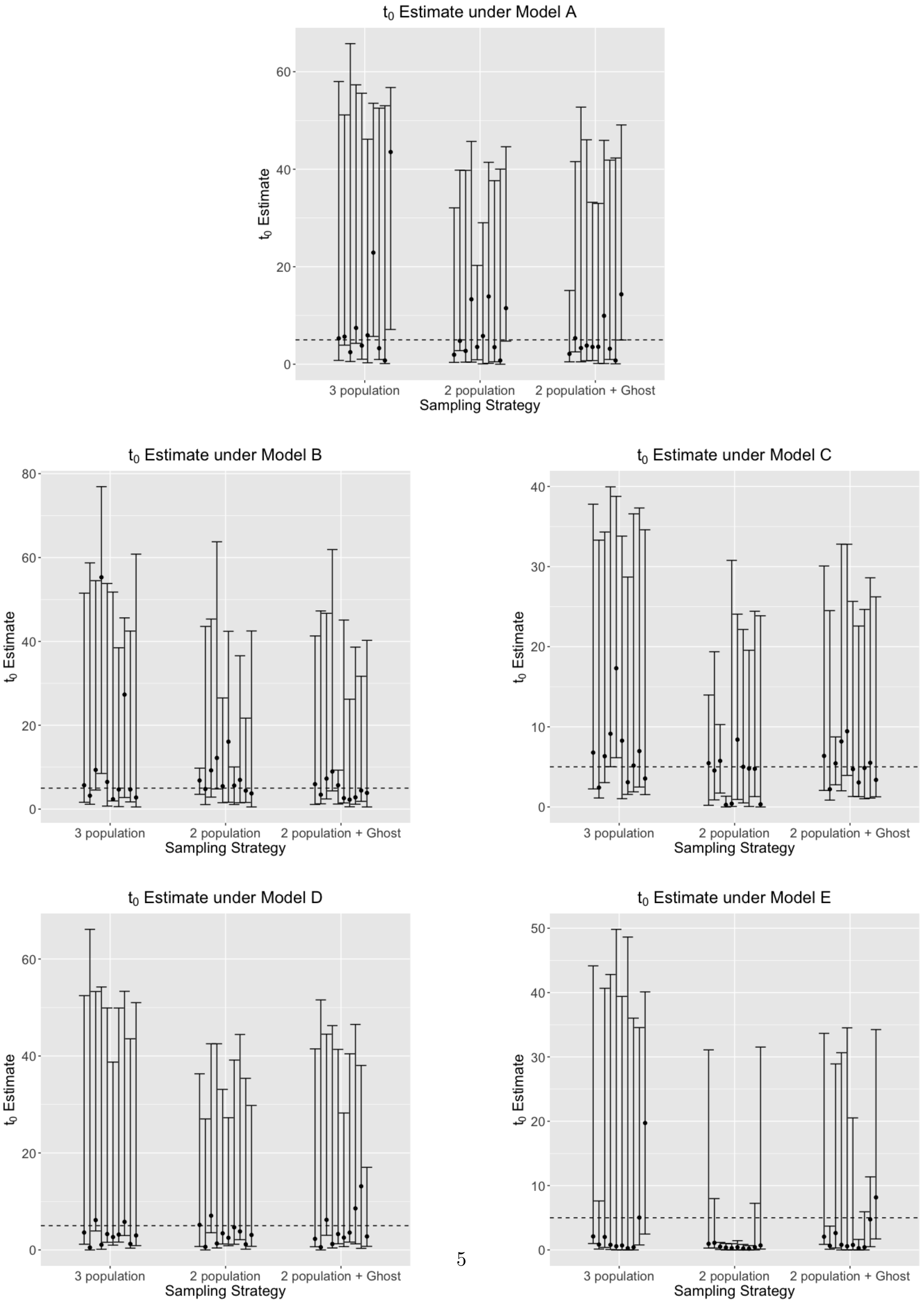

**Figure 3:** Divergence time ($t_0$) estimates between sampled populations, using 5 genomic loci under models A-E, estimated using a 3-population model, 2-population model (without a 'ghost'), and a 2-population model with a 'ghost' outgroup. The dotted line indicates the true (simulated) divergence time ($\frac{t_0}{u} = 5$, where $u$ is the mutation rate per site per generation. Modes, and 95% confidence intervals from 10 replicate runs are shown per model. Compared to the null model A, divergence times are consistently under-estimated with large unidirectional, or bidirectional gene flow from the 'ghost' (models C and E). Compared to estimates with 2 genomic loci (Fig. 2), we observe tighter confidence intervals around the mode while using 5 loci.

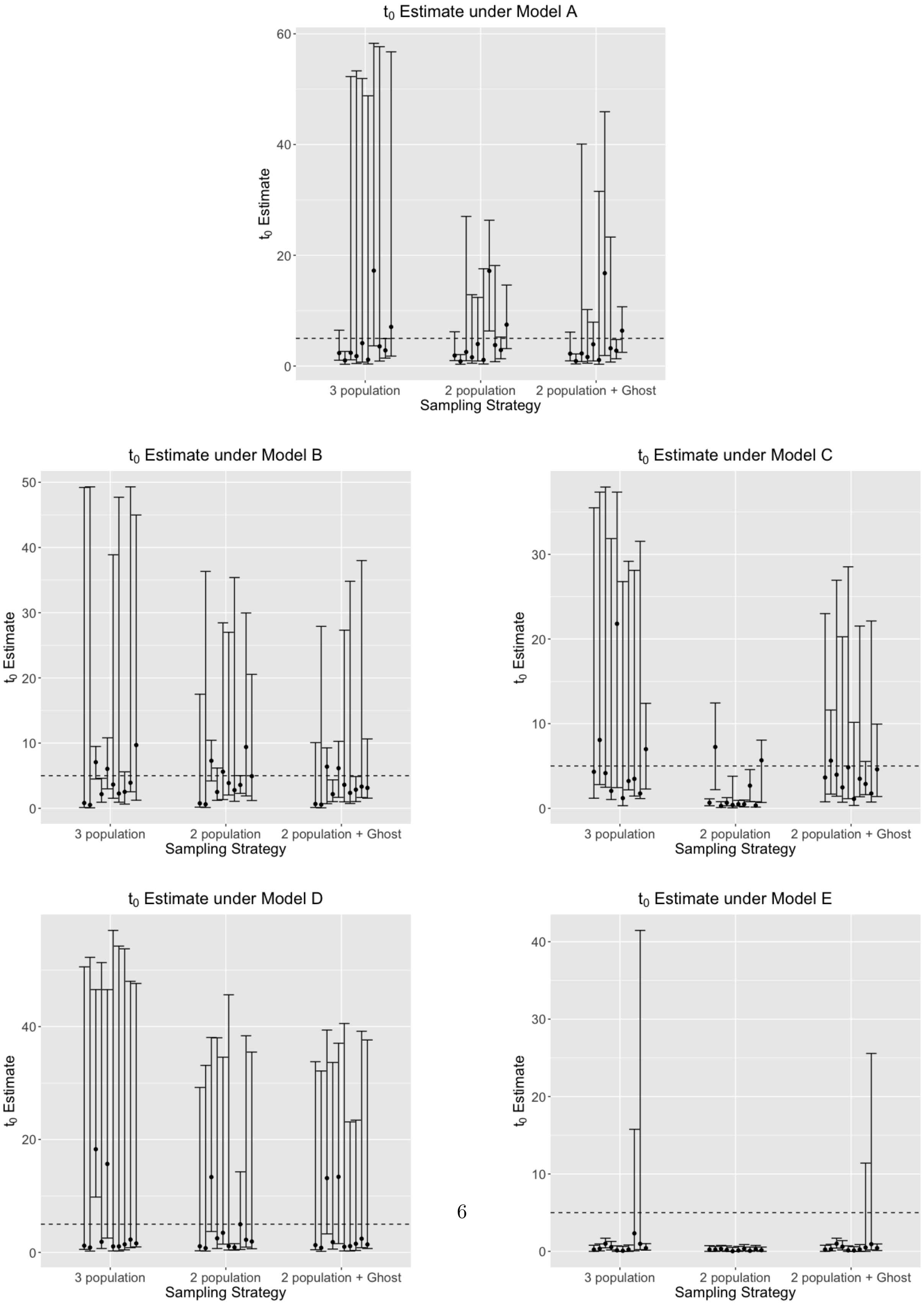

**Figure 4:** Divergence time ($t_1$) estimates between sampled populations, using 2 genomic loci under models A-E, estimated using a 3-population model, 2-population model (without a 'ghost'), and a 2-population model with a 'ghost' outgroup. The dotted line indicates the true (simulated) divergence time ($\frac{t_1}{u} = 20$, where $u$ is the mutation rate per site per generation. Modes, and 95% confidence intervals from 10 replicate runs are shown per model. Compared to the null model A, divergence times are consistently under-estimated with large unidirectional, or bidirectional gene flow from the 'ghost' (models C and E). Compared to estimates with 2 genomic loci (Fig. 4), we observe tighter confidence intervals around the mode while using 5 loci.

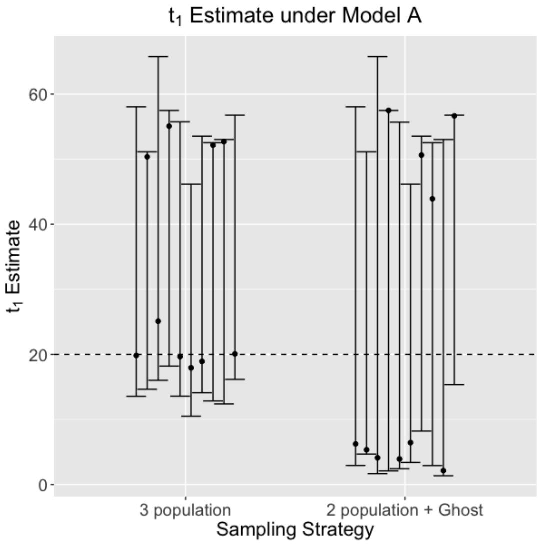


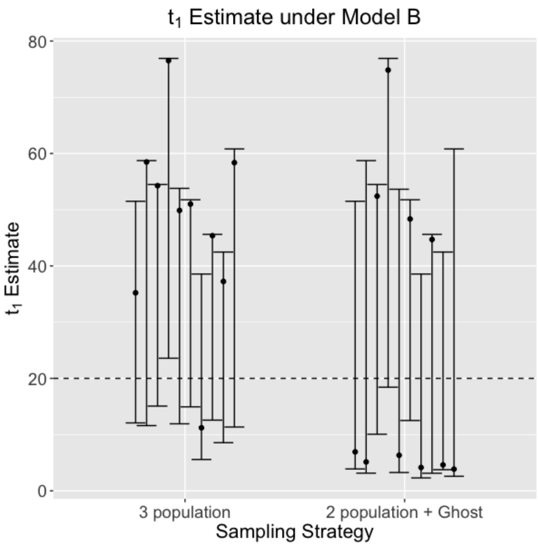


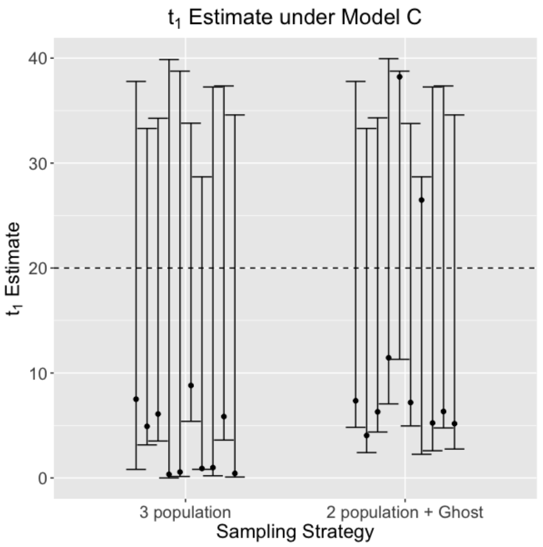


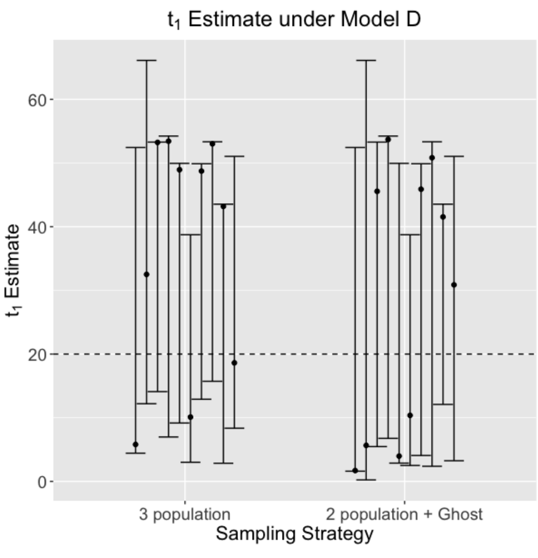


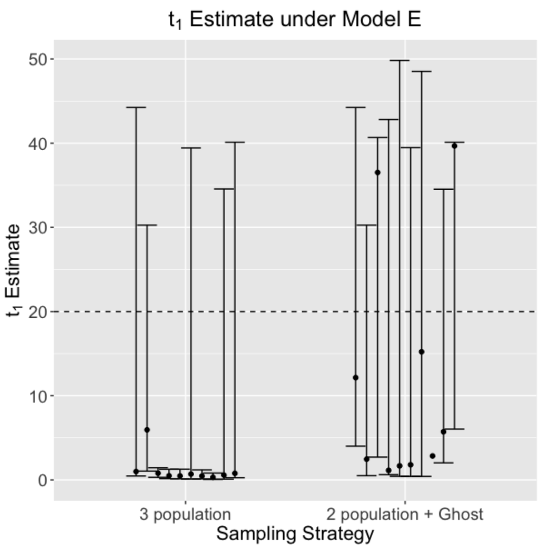

**Figure 5:** Divergence time ($t_1$) estimates between the common ancestor of sampled populations, and the ghost outgroup, using 5 genomic loci under models A-E, estimated using a 3-population model, and a 2-population model with a ghost outgroup.

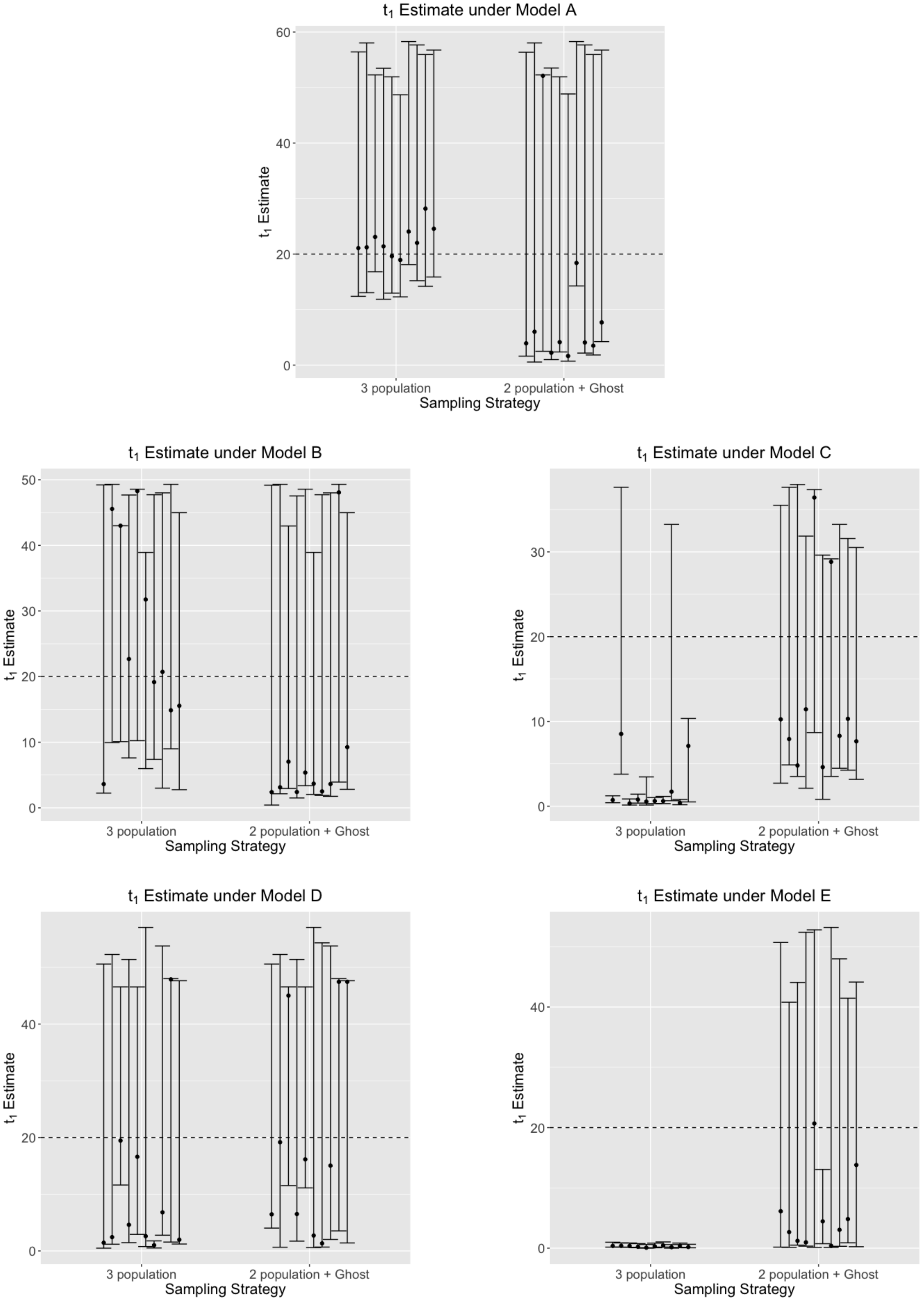

**Figure 6:** Scaled effective population size ($\theta_0 = 4N_0u$) estimate of the first sampled populations, using 2 genomic loci under models A-E, estimated using a 3-population model, 2-population model (without a 'ghost'), and a 2-population model with a 'ghost' outgroup. True simulated $\theta_0 = 10$, and is shown with the dotted line. $\theta_0$ is consistently over-estimated, as seen by the inflated confidence intervals around the mode, with increased bi-directional gene flow from the 'ghost' (models D and E), compared to the null model of no gene flow from the 'ghost' (A).

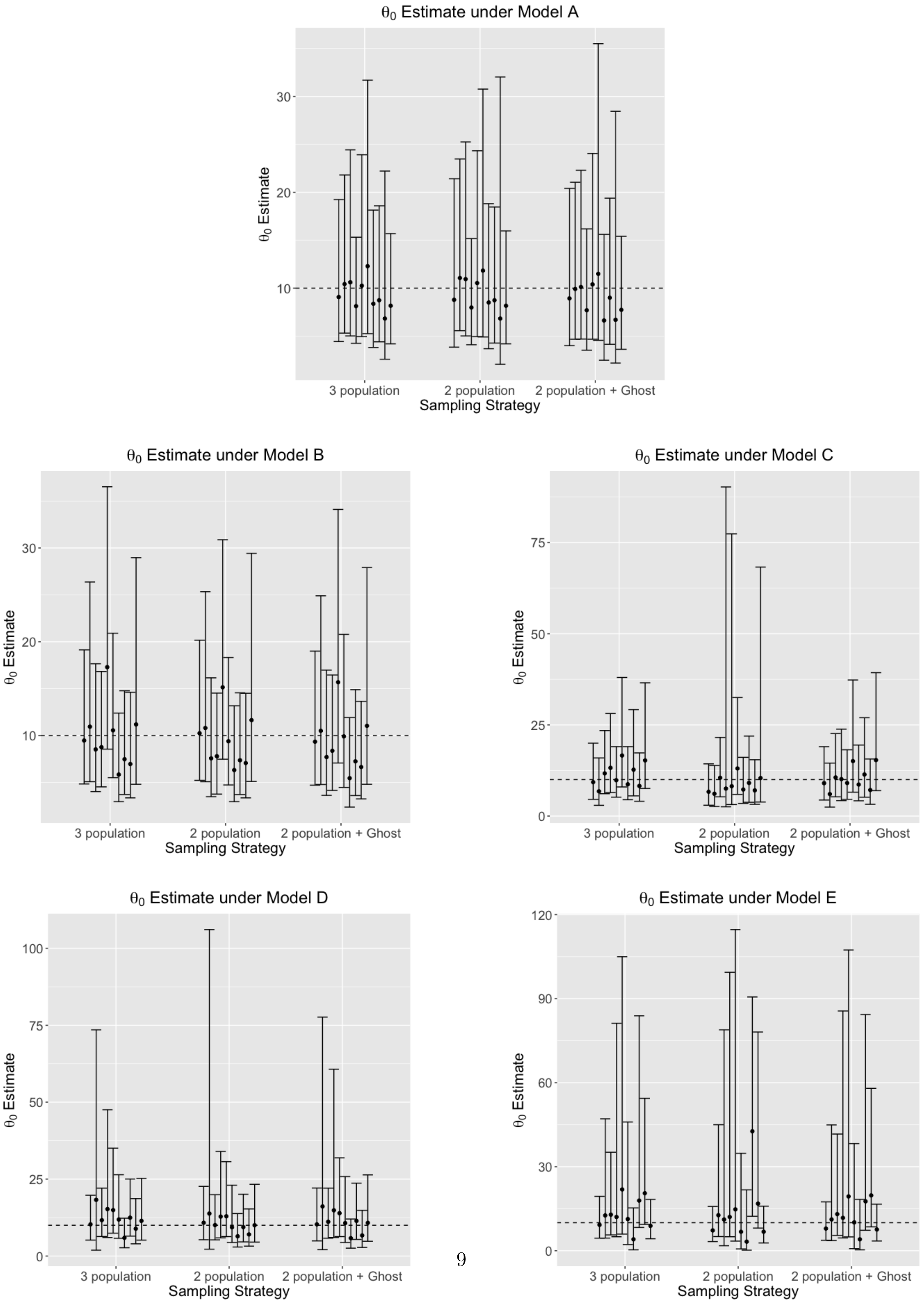

**Figure 7:** Scaled effective population size ($\theta_0 = 4N_0u$) estimate of the first sampled populations, using 5 genomic loci under models A-E, estimated using a 3-population model, 2-population model (without a 'ghost'), and a 2-population model with a 'ghost' outgroup. True simulated $\theta_0 = 10$, and is shown with the dotted line. $\theta_0$ is consistently over-estimated, as seen by the inflated confidence intervals around the mode, with increased bi-directional gene flow from the 'ghost' (models D and E), compared to the null model of no gene flow from the 'ghost' (A). Compared to estimates using 2 genomic loci (Fig. 6), we observe tighter confidence intervals around the mode while using 5 genomic loci across all models (A-E).

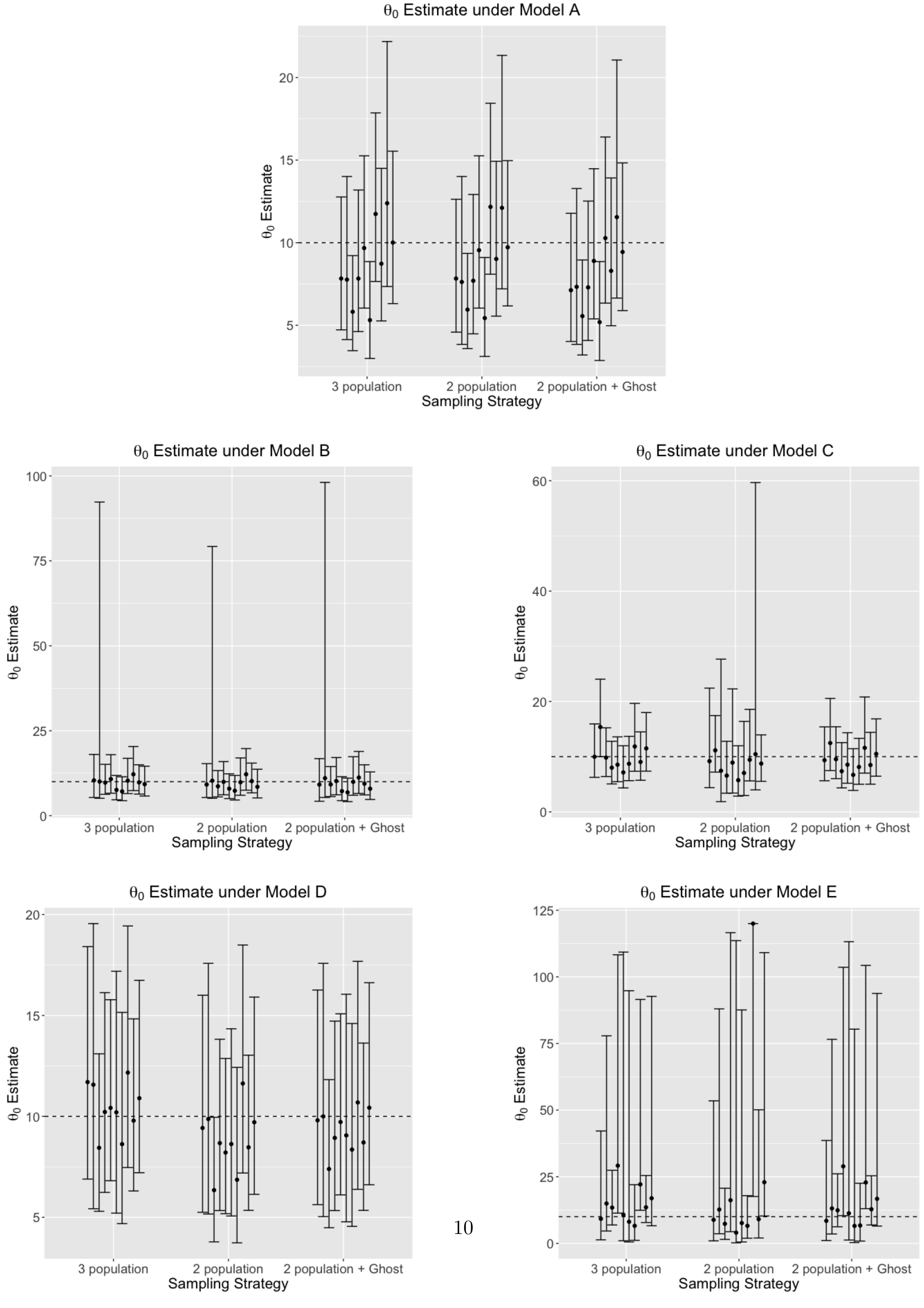

**Figure 8:** Scaled effective population size ($\theta_1 = 4N_1u$) estimate of the second sampled populations, using 2 genomic loci under models A-E, estimated using a 3-population model, 2-population model (without a 'ghost'), and a 2-population model with a 'ghost' outgroup. True simulated $\theta_1 = 10$, and is shown with the dotted line. $\theta_1$ is consistently over-estimated, as seen by the inflated confidence intervals around the mode, with increased bi-directional gene flow from the 'ghost' (models D and E), compared to the null model of no gene flow from the 'ghost' (A).

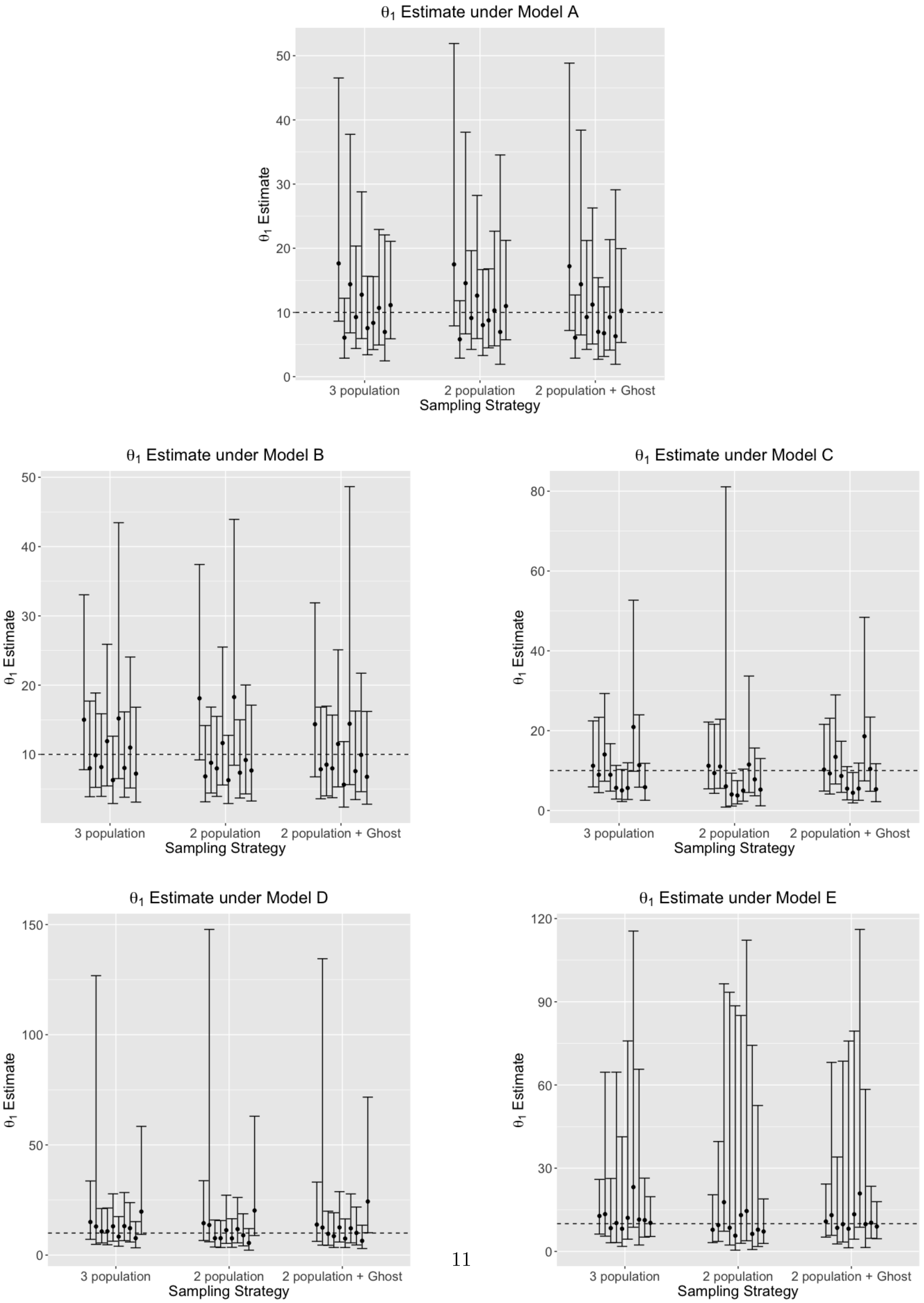

**Figure 9:** Scaled effective population size ($\theta_1 = 4N_1u$) estimate of the first sampled populations, using 2 genomic loci under models A-E, estimated using a 3-population model, 2-population model (without a 'ghost'), and a 2-population model with a 'ghost' outgroup. True simulated $\theta_1 = 10$, and is shown with the dotted line. $\theta_1$ is consistently over-estimated, as seen by the inflated confidence intervals around the mode, with increased bi-directional gene flow from the 'ghost' (models D and E), compared to the null model of no gene flow from the 'ghost' (A). Compared to estimates using 2 genomic loci (Fig. 8), we observe tighter confidence intervals around the mode while using 5 genomic loci across all models (A-E).

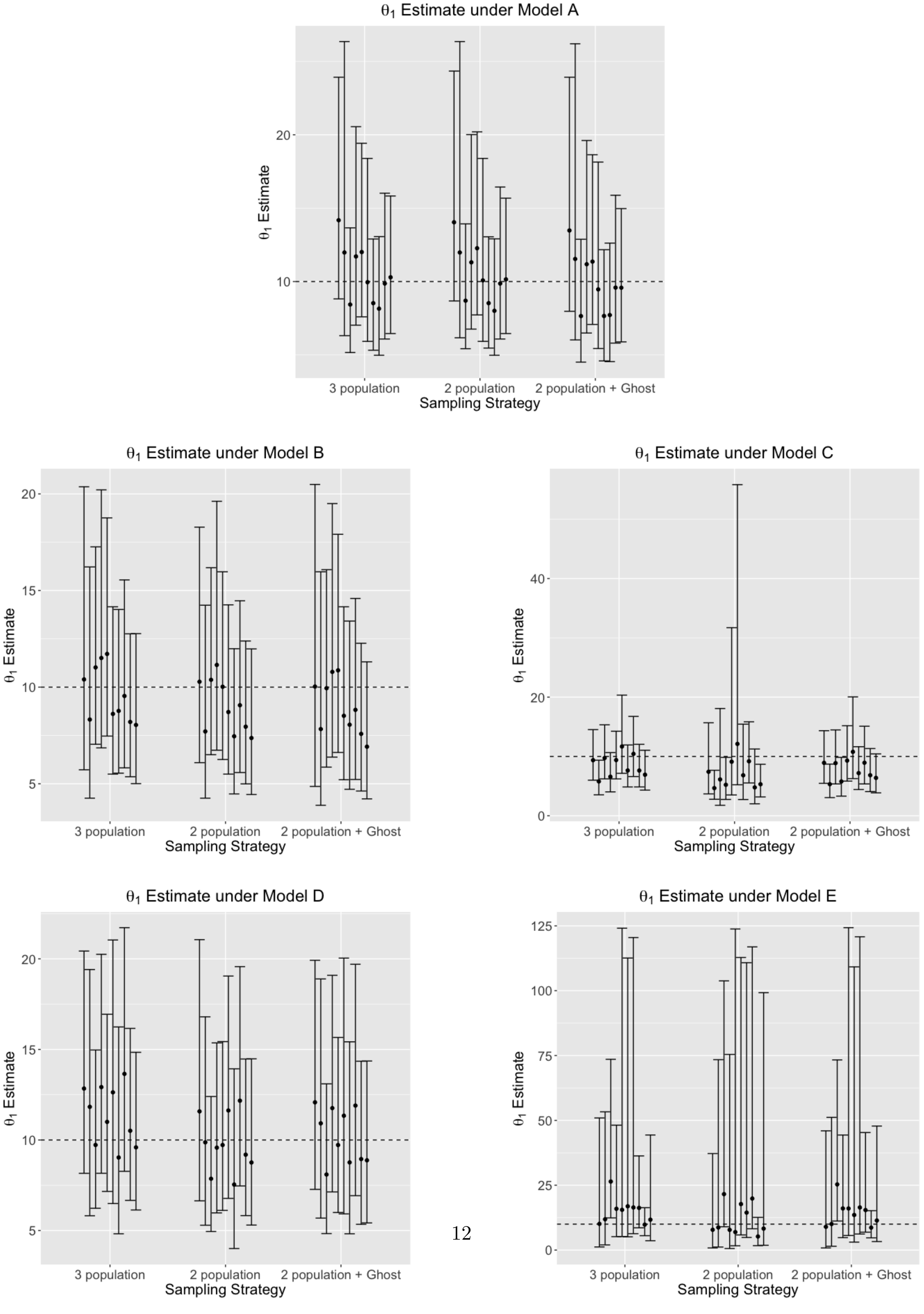

**Figure 10:** Scaled effective population size ($\theta_2 = 4N_2u$) estimate of the common ancestor of the two sampled populations, using 2 genomic loci under models A-E, estimated using a 3-population model, 2-population model (without a 'ghost'), and a 2-population model with a 'ghost' outgroup. True simulated $\theta_2 = 10$, and is shown with the dotted line. $\theta_2$ is consistently over-estimated, as seen by the inflated confidence intervals around the mode, under all models (A-E). The 2-population model (without a 'ghost') however (under models A, B, and D, with none, to little unidirectional gene flow from the 'ghost') show accurate estimates of $\theta_2$. These are nonetheless over-estimated with increased gene flow from the 'ghost' (models C, E).

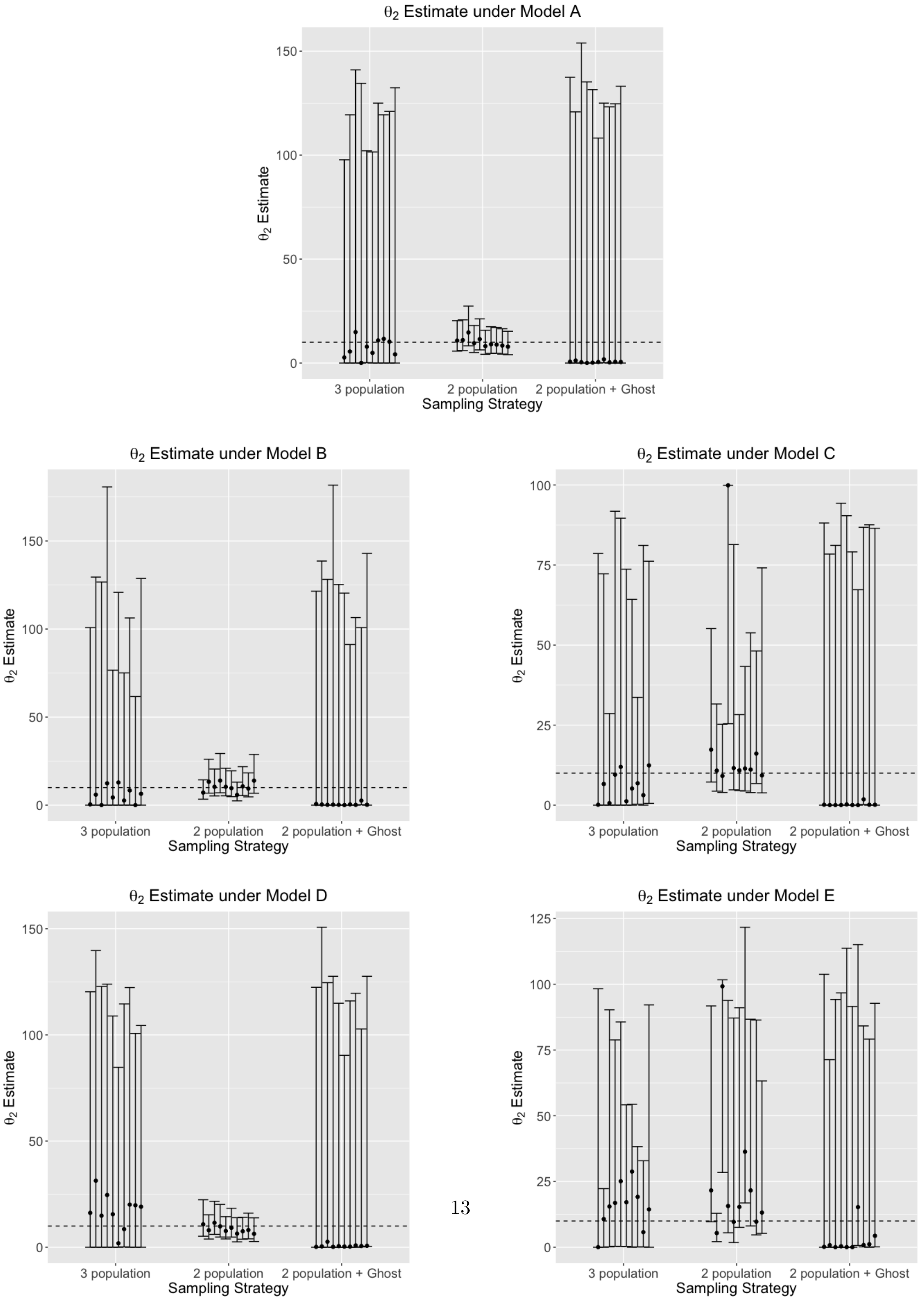

**Figure 11:** Scaled effective population size ($\theta_2 = 4N_2u$) estimate of the common ancestor of the two sampled populations, using 5 genomic loci under models A-E, estimated using a 3-population model, 2-population model (without a 'ghost'), and a 2-population model with a 'ghost' outgroup. True simulated $\theta_2 = 10$, and is shown with the dotted line. $\theta_2$ is consistently over-estimated, as seen by the inflated confidence intervals around the mode, under all models (A-E). The 2-population model (without a 'ghost') however (under models A, B, and D, with none, to little unidirectional gene flow from the 'ghost') show accurate estimates of $\theta_2$. These are nonetheless over-estimated with increased gene flow from the 'ghost' (models C, E). Compared to the 2 locus estimates (Fig. 10), we observe tighter confidence intervals around the mode across all models while using 5 genomic loci.

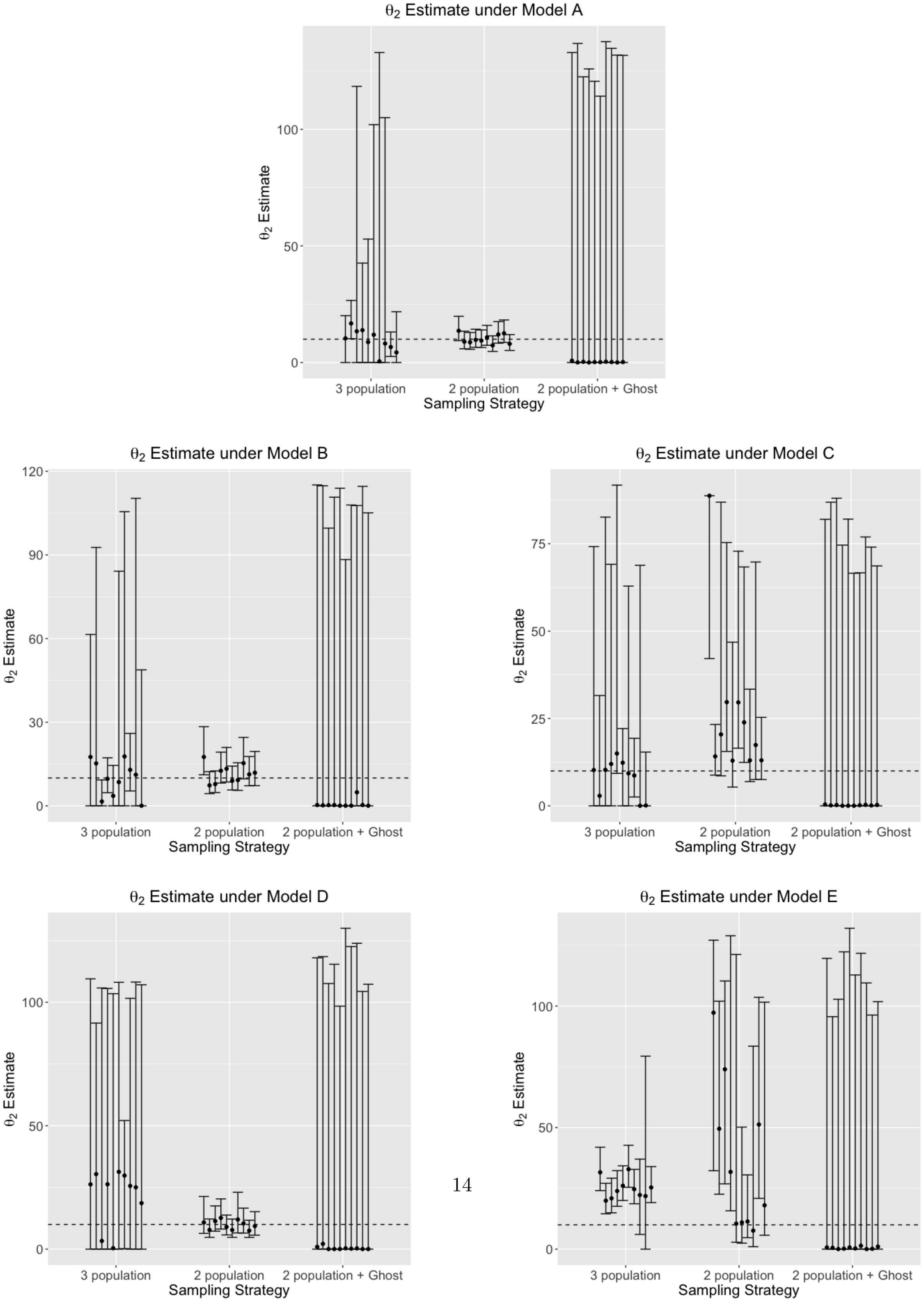

**Figure 12:** Scaled effective population size ($\theta_3 = 4N_3u$) estimate of the unsampled 'ghost', using 2 genomic loci under models A-E, estimated using a 3-population model, and a 2-population model with a 'ghost' outgroup. True simulated $\theta_3 = 10$, and is shown with the dotted line. $\theta_3$ is consistently over-estimated, as seen by the inflated confidence intervals around the mode, under all models (A-E).

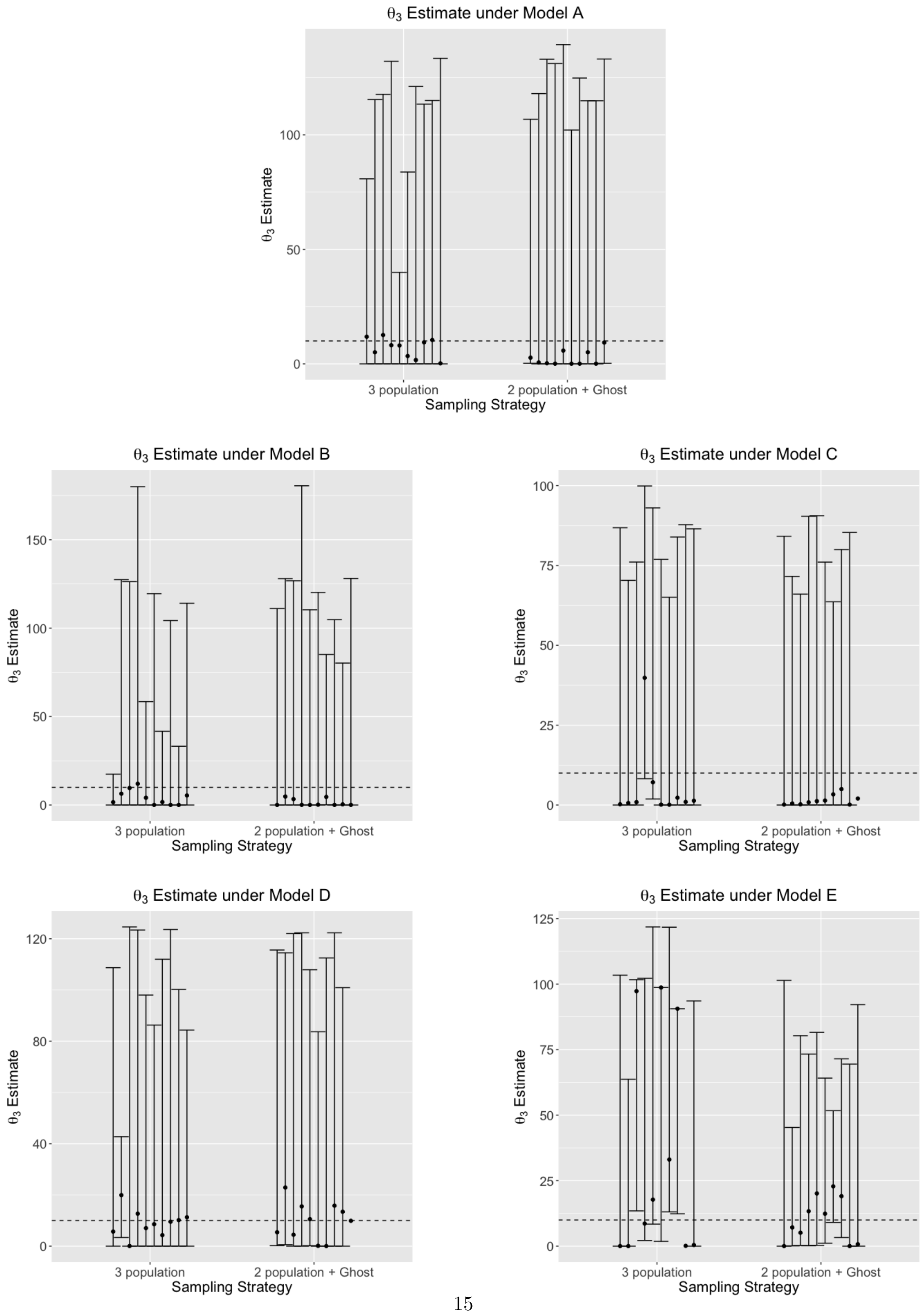

**Figure 13:** Scaled effective population size ($\theta_3 = 4N_3u$) estimate of the unsampled 'ghost', using 5 genomic loci under models A-E, estimated using a 3-population model, and a 2-population model with a 'ghost' outgroup. True simulated $\theta_3 = 10$, and is shown with the dotted line. $\theta_3$ is consistently over-estimated, as seen by the inflated confidence intervals around the mode, under all models (A-E).

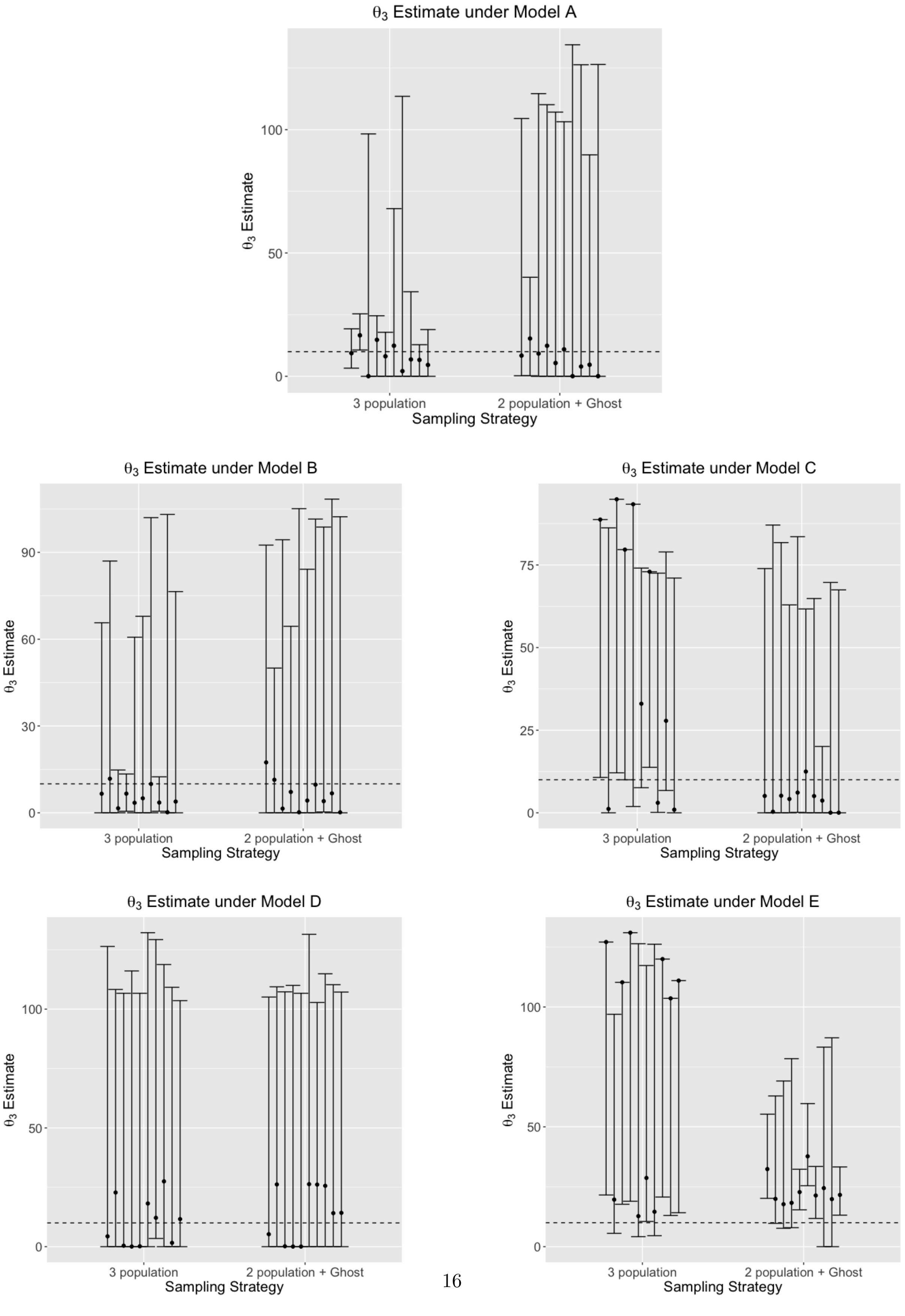

**Figure 14:** Scaled effective population size ($\theta_4 = 4N_4u$) estimate of the common ancestor of unsampled 'ghost', and the common ancestor of the sampled populations using 2 genomic loci under models A-E, estimated using a 3-population model, and a 2-population model with a 'ghost' outgroup. True simulated $\theta_4 = 10$, and is shown with the dotted line. $\theta_4$ is consistently over-estimated, as seen by the inflated confidence intervals around the mode, under all models (A-E).

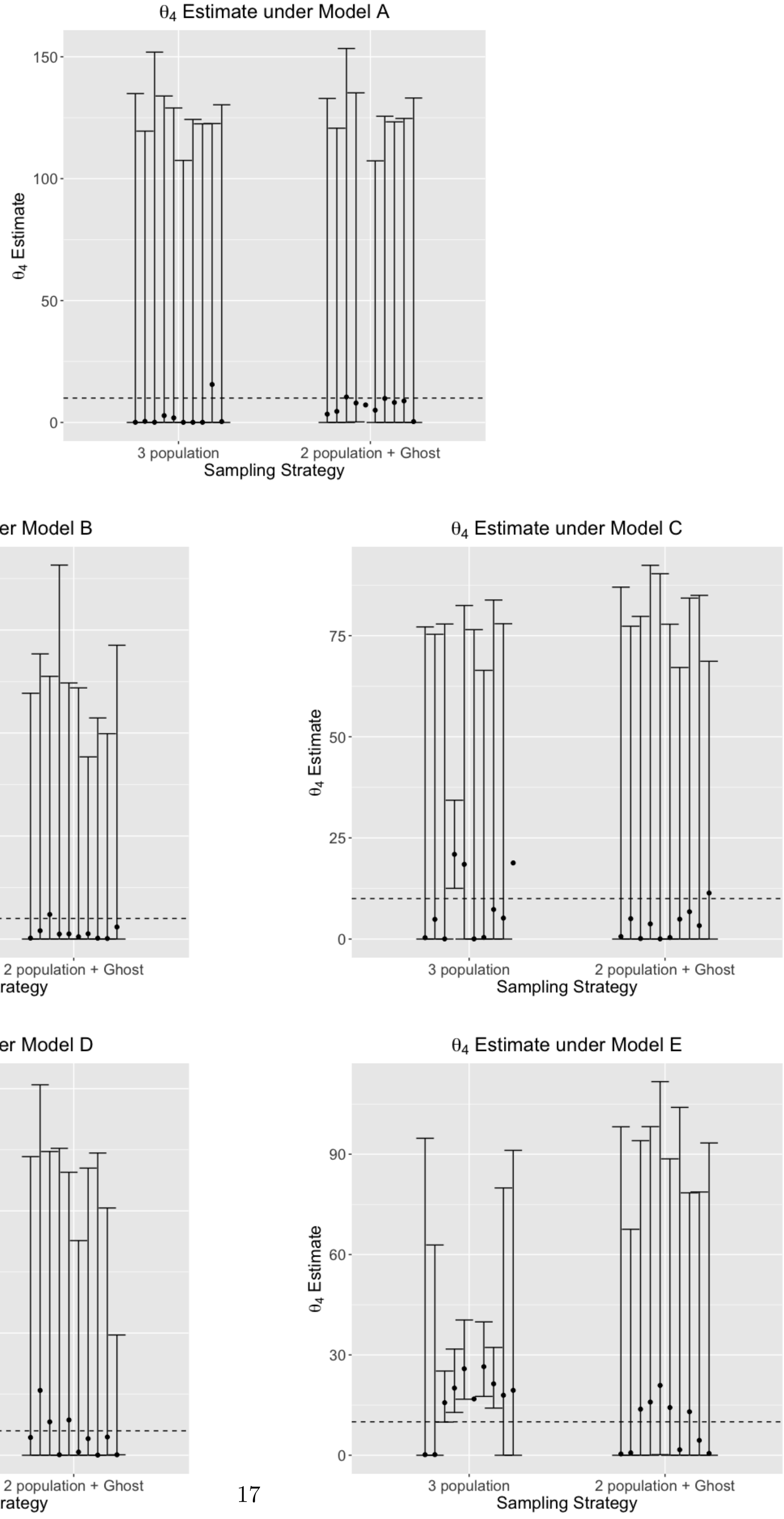

**Figure 15:** Scaled effective population size ($\theta_4 = 4N_4u$) estimate of the common ancestor of unsampled 'ghost', and the common ancestor of the sampled populations using 5 genomic loci under models A-E, estimated using a 3-population model, and a 2-population model with a 'ghost' outgroup. True simulated $\theta_4 = 10$, and is shown with the dotted line. $\theta_4$ is consistently over-estimated, as seen by the inflated confidence intervals around the mode, under all models (A-E). Compared to using 2 loci (Fig. 14), all estimates are more accurate while using 5 loci.

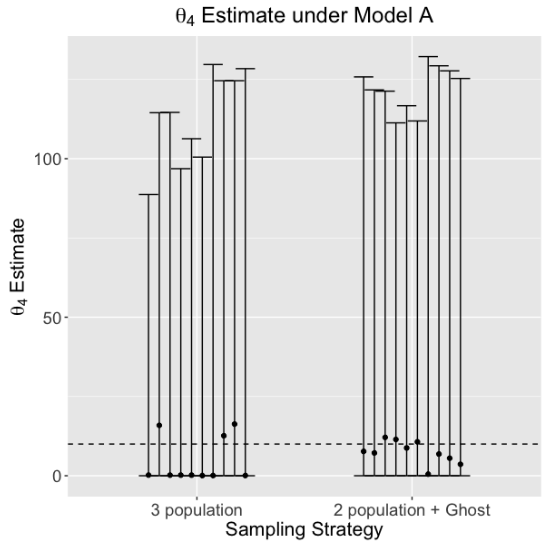


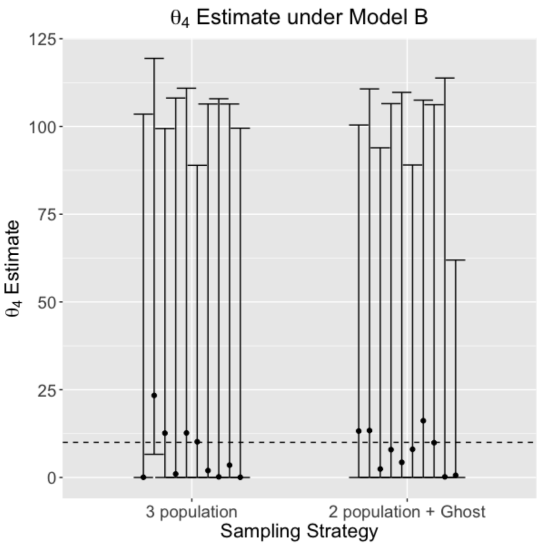


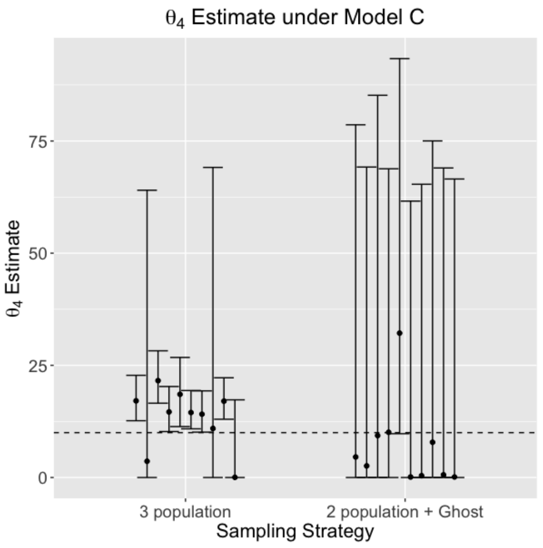


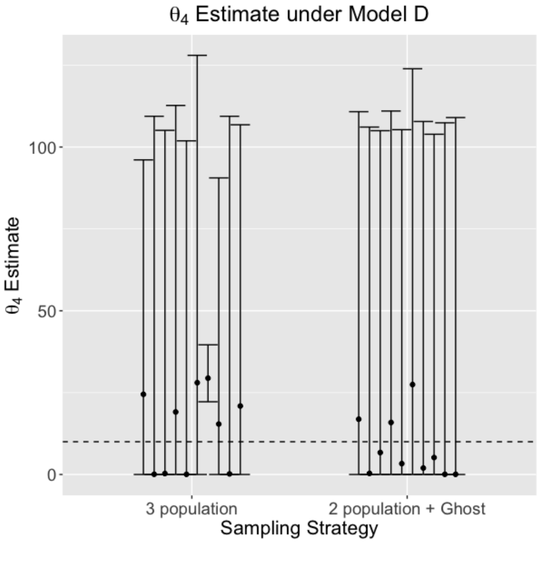


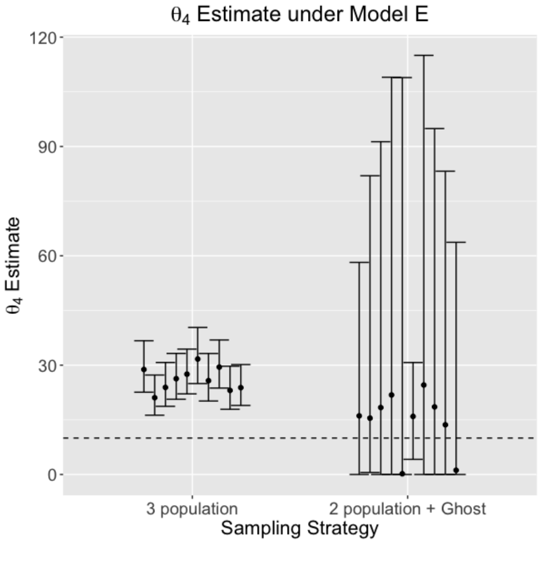

**Figure 16:** Scaled migration rate ($m_{1\to0} = 4N_1u$, where $N_1$ is the effective population size of one of the sampled populations, and $u$ is the mutation rate per site per generation) estimate between the two sampled populations, using 2 genomic loci under models A-E, estimated using a 3-population model, 2-population model (without a 'ghost'), and a 2-population model with a ghost outgroup. True simulated $m_{1\to0} = 0.05$ and is shown with a dotted line. Models with increased gene flow from the ghost (both unidirectional - C, and bi-directional - E) consistently lead to over-estimation of $m_{1\to0}$, as observed by the inflated confidence intervals around the mode.

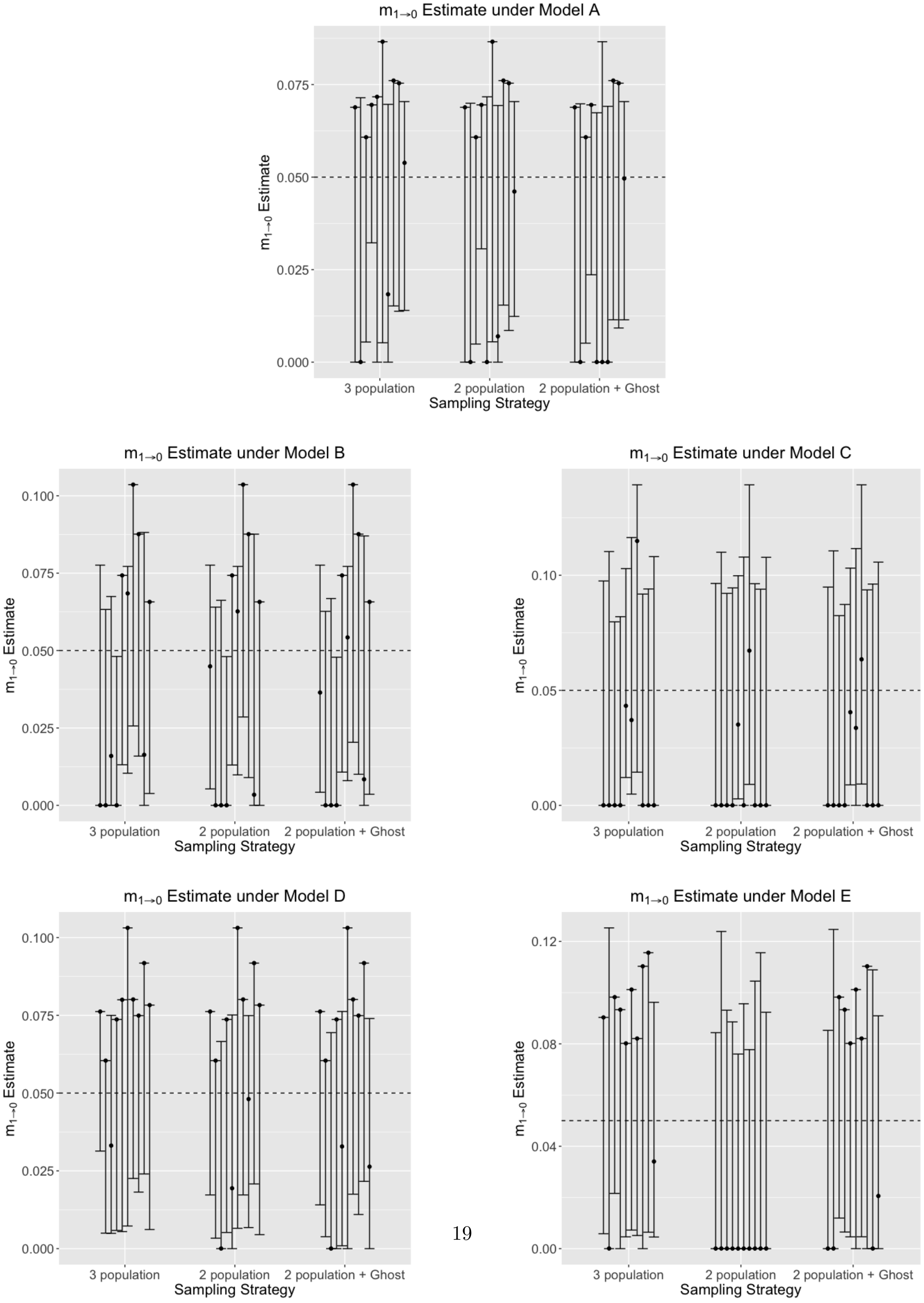

**Figure 17:** Scaled migration rate ($m_{0\to 1} = 4N_0u$, where $N_0$ is the effective population size of one of the sampled populations, and $u$ is the mutation rate per site per generation) estimate between the two sampled populations, using 5 genomic loci under models A-E, estimated using a 3-population model, 2-population model (without a 'ghost'), and a 2-population model with a ghost outgroup. True simulated $m_{0\to 1} = 0.05$, and is shown with the dotted line. Models with increased gene flow from the ghost (both unidirectional - C, and bi-directional - E) consistently lead to over-estimation of $m_{0\to 1}$, as observed by the inflated confidence intervals around the mode. Compared to using 2 genomic loci (Fig. 19), using 5 genomic loci leads to more accurate estimates, as observed by the smaller confidence intervals around the mode.

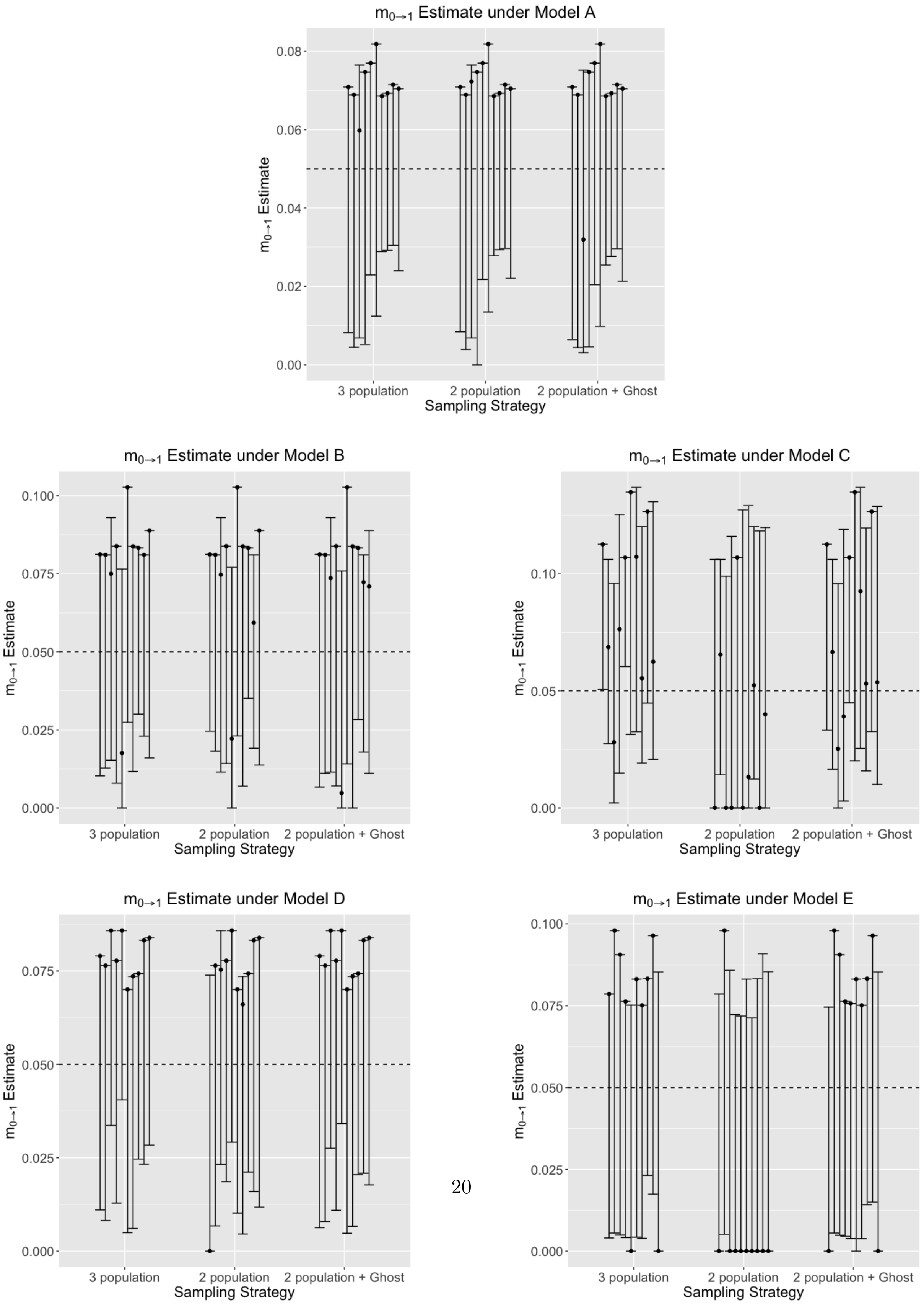

**Figure 18:** Scaled migration rate ($m_{1\to0} = 4N_1u$, where $N_1$ is the effective population size of one of the sampled populations, and $u$ is the mutation rate per site per generation) estimate between the two sampled populations, using 5 genomic loci under models A-E, estimated using a 3-population model, 2-population model (without a 'ghost'), and a 2-population model with a ghost outgroup. True simulated $m_{1\to0} = 0.05$ and is shown with a dotted line. Models with increased gene flow from the ghost (both unidirectional - C, and bi-directional - E) consistently lead to over-estimation of $m_{1\to0}$, as observed by the inflated confidence intervals around the mode. Compared to using 2 genomic loci (Fig. 16), using 5 genomic loci leads to more accurate estimates, as observed by the smaller confidence intervals around the mode.

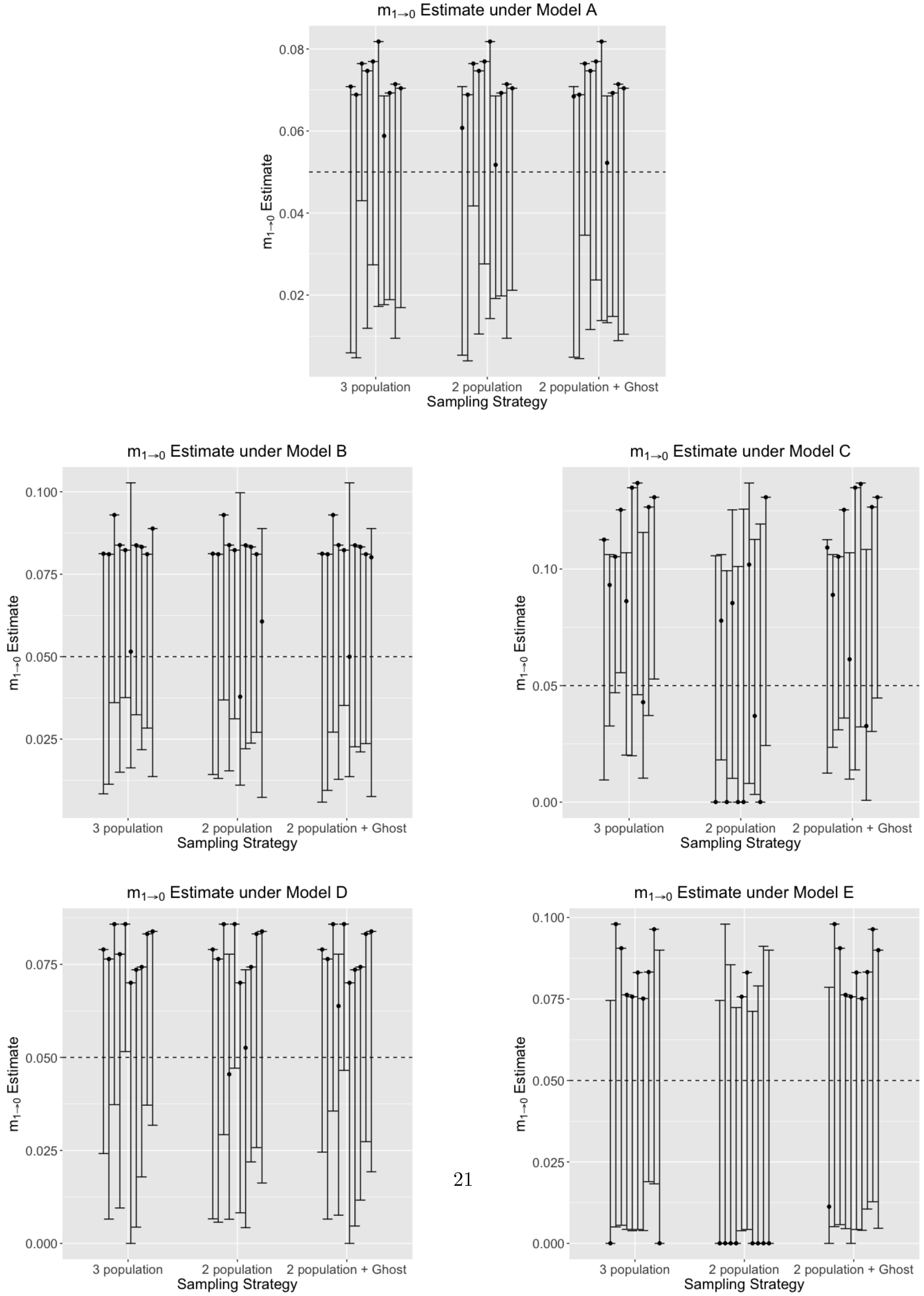

**Figure 19:** Scaled migration rate ($m_{0\to1} = 4N_0u$, where $N_0$ is the effective population size of one of the sampled populations, and $u$ is the mutation rate per site per generation) estimate between the two sampled populations, using 2 genomic loci under models A-E, estimated using a 3-population model, 2-population model (without a 'ghost'), and a 2-population model with a ghost outgroup. True simulated $m_{0\to1} = 0.05$ and is shown with a dotted line. Models with increased gene flow from the ghost (both unidirectional - C, and bi-directional - E) consistently lead to over-estimation of $m_{0\to1}$, as observed by the inflated confidence intervals around the mode.

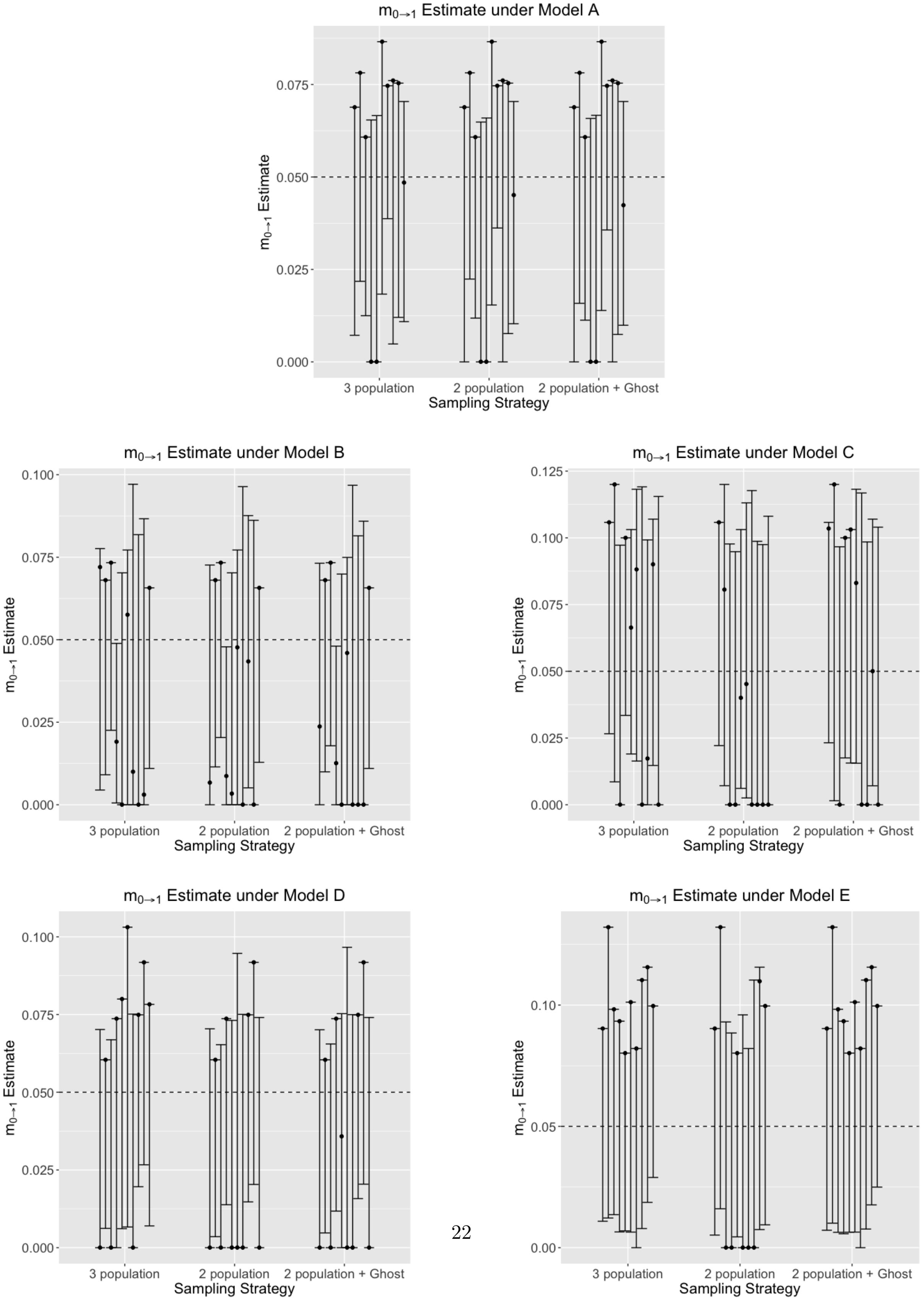

**Figure 20:** Scaled migration rate ($m_{2\to0} = 4N_2u$, where $N_2$ is the effective population size of the 'ghost', and $u$ is the mutation rate per site per generation) estimate between one of the sampled populations and the unsampled 'ghost', using 2 genomic loci under models A-E, estimated using a 3-population model, and a 2-population model with a ghost outgroup. True simulated migration rates from the ghost vary per model, and are shown with the dotted line. Across all models, we consistently under-estimate the migration rate from the 'ghost', especially with increased gene flow (models C and E).

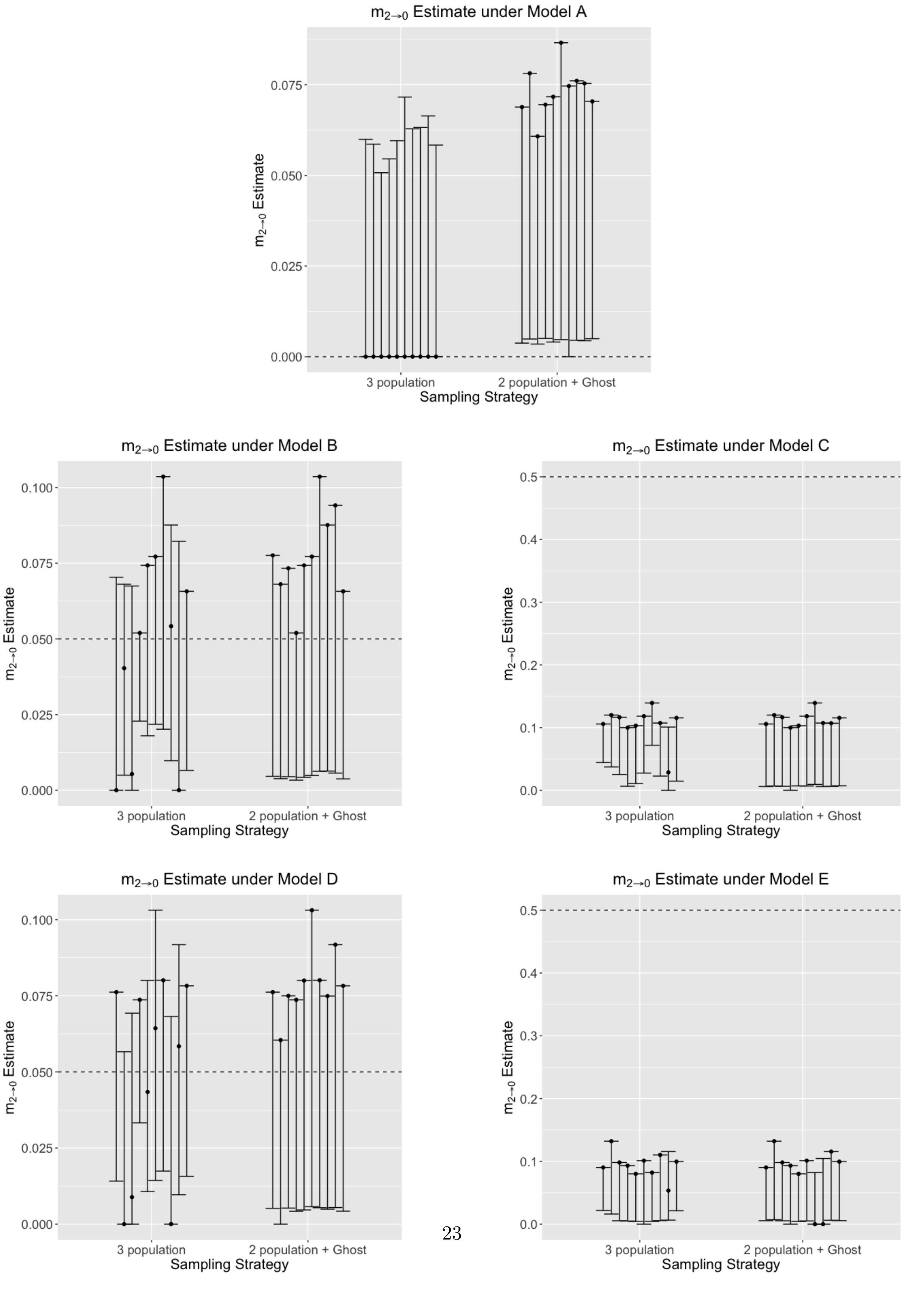

**Figure 21:** Scaled migration rate ($m_{2\to0} = 4N_2u$, where $N_2$ is the effective population size of the 'ghost', and $u$ is the mutation rate per site per generation) estimate between one of the sampled populations and the unsampled 'ghost', using 5 genomic loci under models A-E, estimated using a 3-population model, and a 2-population model with a ghost outgroup. True simulated migration rates from the ghost vary per model, and are shown with the dotted line. Across all models, we consistently under-estimate the migration rate from the 'ghost', especially with increased gene flow (models C and E).

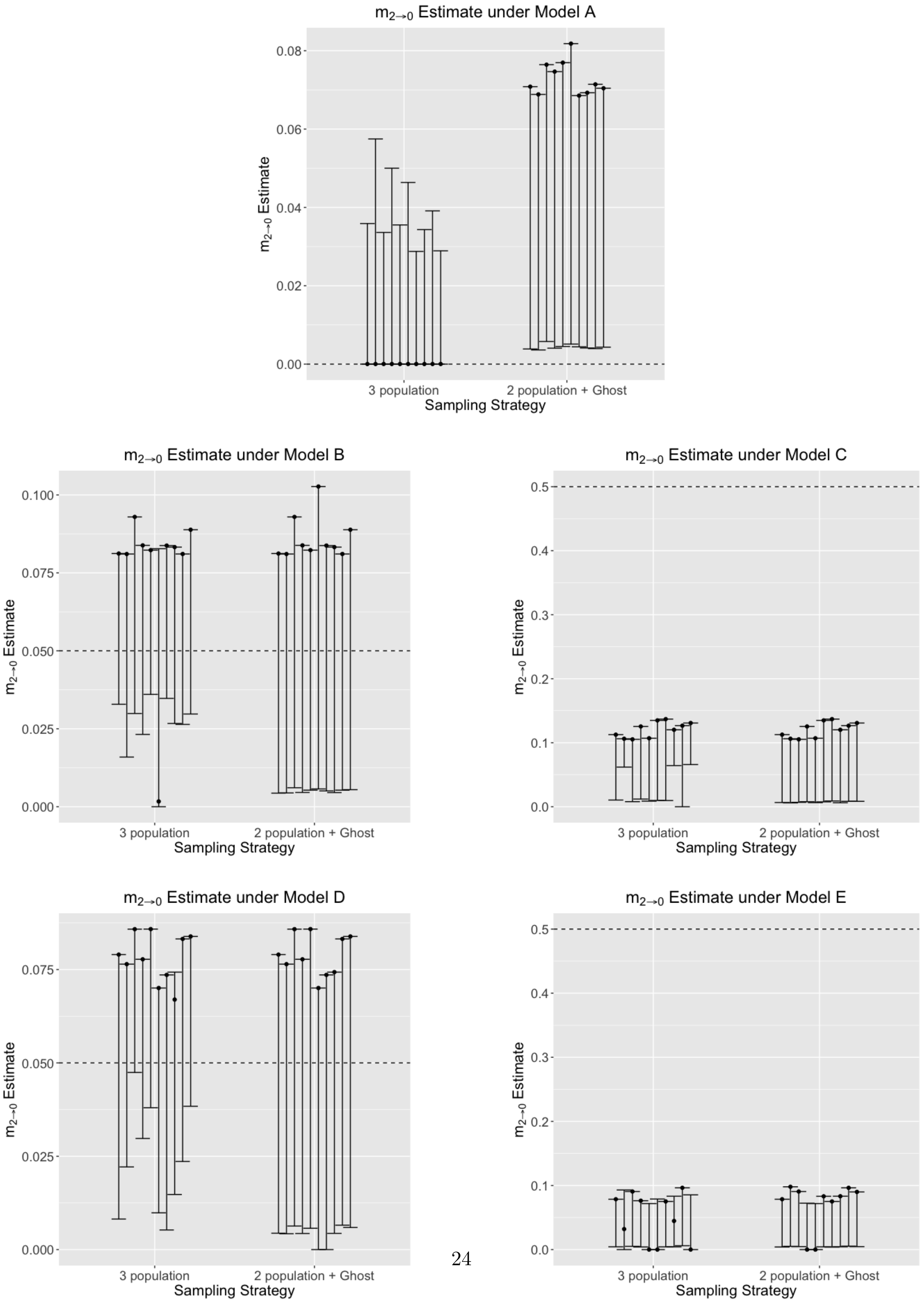

**Figure 22:** Scaled migration rate ($m_{0\to2} = 4N_0u$, where $N_0$ is the effective population size of a sampled population, and $u$ is the mutation rate per site per generation) estimate between one of the sampled populations and the unsampled ‘ghost’, using 2 genomic loci under models A-E, estimated using a 3-population model, and a 2-population model with a ghost outgroup. True simulated migration rates from the ghost vary per model, and are shown with the dotted line. Across all models, we consistently under-estimate the migration rate from the ‘ghost’, especially with increased gene flow (models C and E).

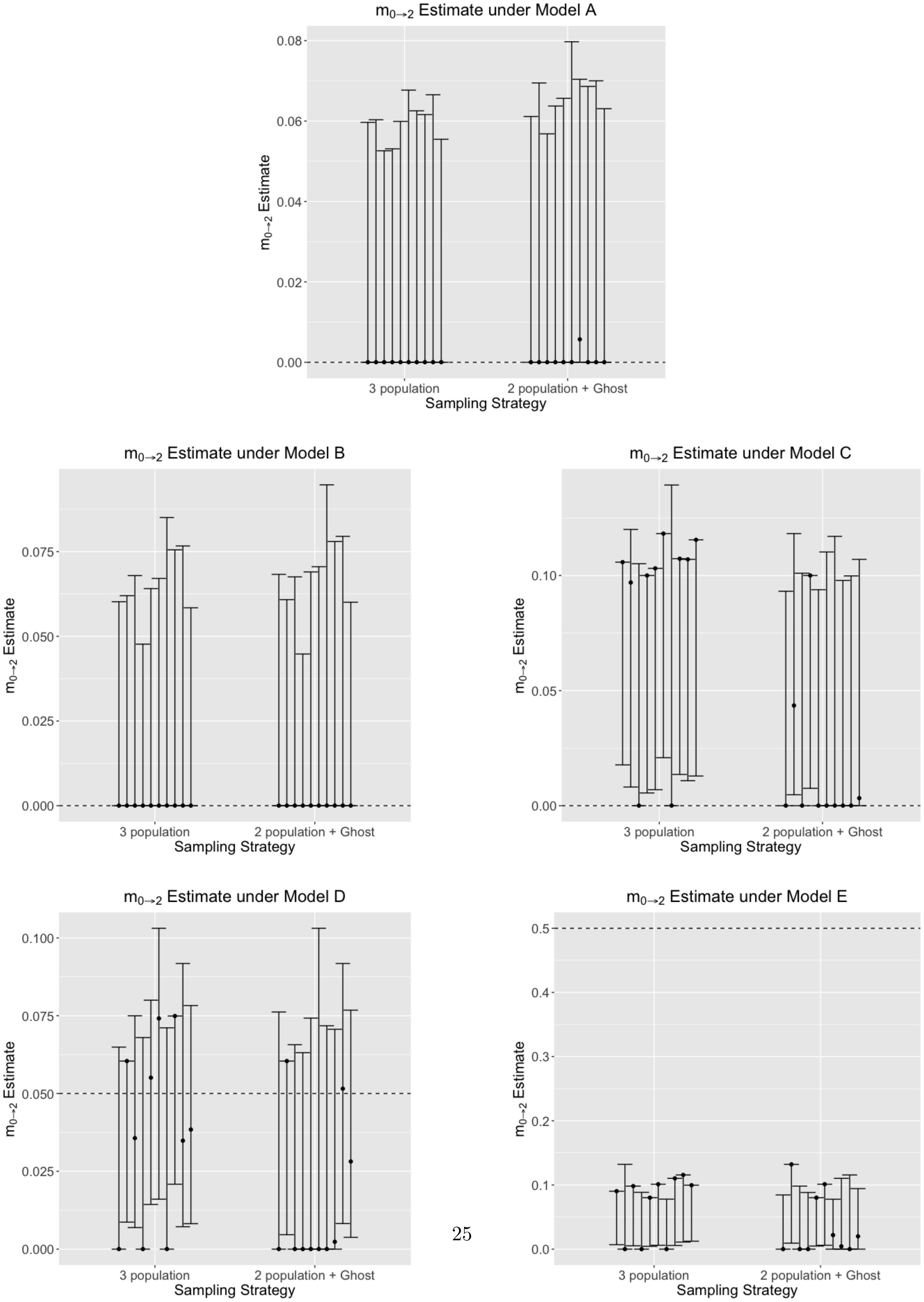

**Figure 23:** Scaled migration rate ($m_{0\to2} = 4N_0u$, where $N_0$ is the effective population size of a sampled population, and $u$ is the mutation rate per site per generation) estimate between one of the sampled populations and the unsampled 'ghost', using 5 genomic loci under models A-E, estimated using a 3-population model, and a 2-population model with a ghost outgroup. True simulated migration rates from the ghost vary per model, and are shown with the dotted line. Across all models, we consistently under-estimate the migration rate from the 'ghost', especially with increased gene flow (models C and E).

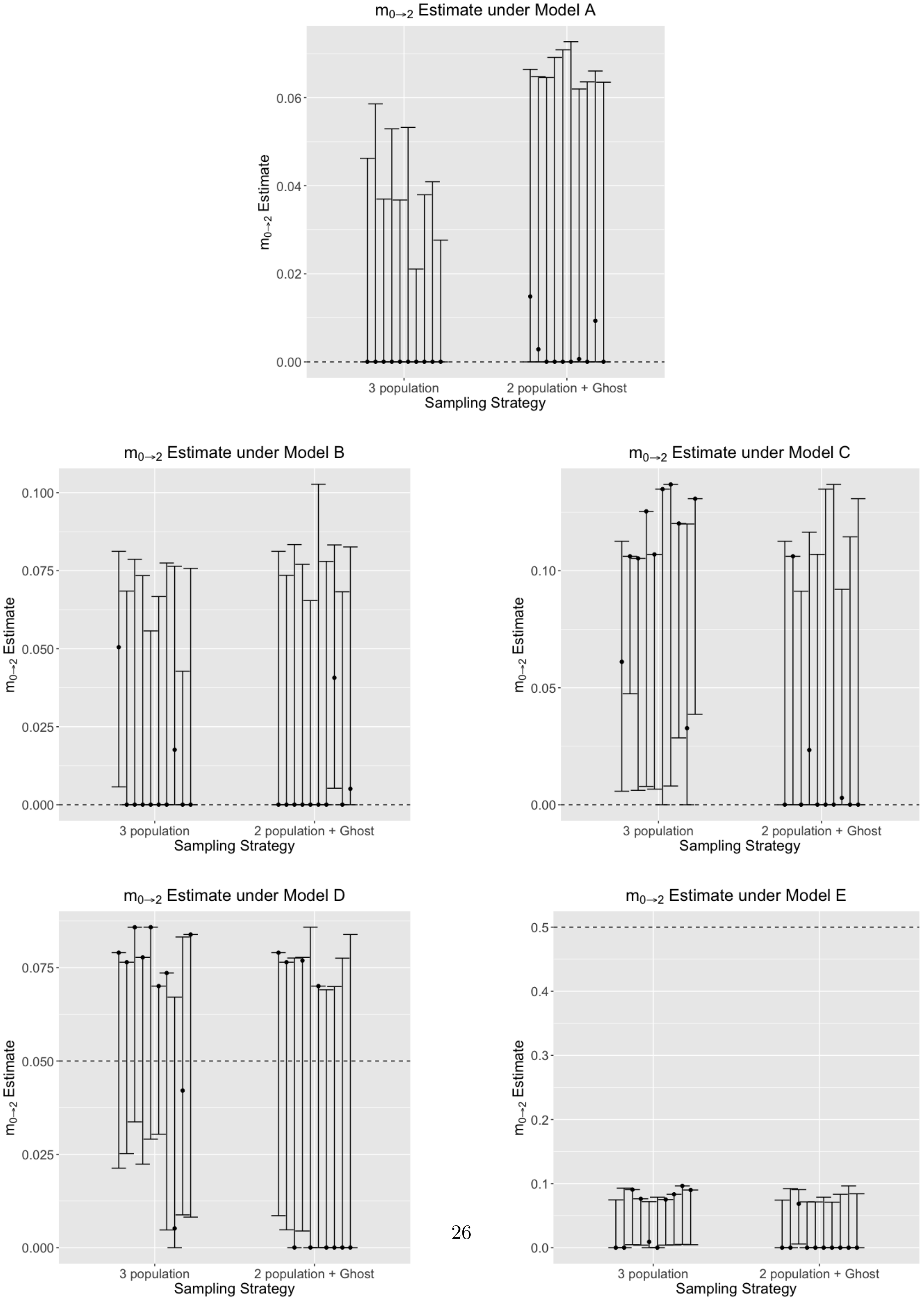

**Figure 24:** Scaled migration rate ($m_{2\to 1} = 4N_2u$, where $N_2$ is the effective population size of the 'ghost', and $u$ is the mutation rate per site per generation) estimate between one of the sampled populations and the unsampled 'ghost', using 2 genomic loci under models A-E, estimated using a 3-population model, and a 2-population model with a ghost outgroup. True simulated migration rates from the ghost vary per model, and are shown with the dotted line. Across all models, we consistently under-estimate the migration rate from the 'ghost', especially with increased gene flow (models C and E).

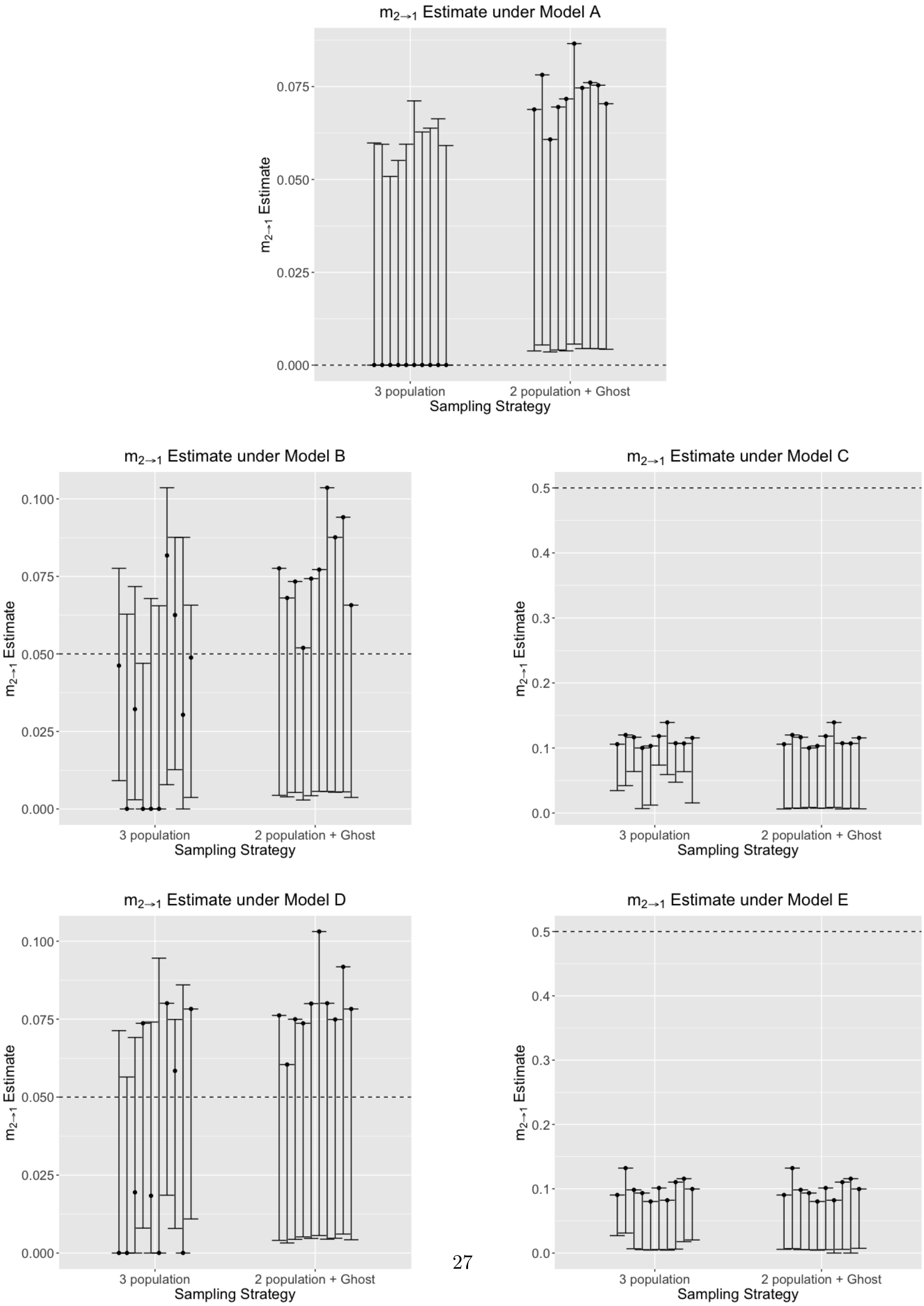

**Figure 25:** Scaled migration rate ($m_{2\to1} = 4N_2u$, where $N_2$ is the effective population size of the 'ghost', and $u$ is the mutation rate per site per generation) estimate between one of the sampled populations and the unsampled 'ghost', using 5 genomic loci under models A-E, estimated using a 3-population model, and a 2-population model with a ghost outgroup. True simulated migration rates from the ghost vary per model, and are shown with the dotted line. Across all models, we consistently under-estimate the migration rate from the 'ghost', especially with increased gene flow (models C and E).

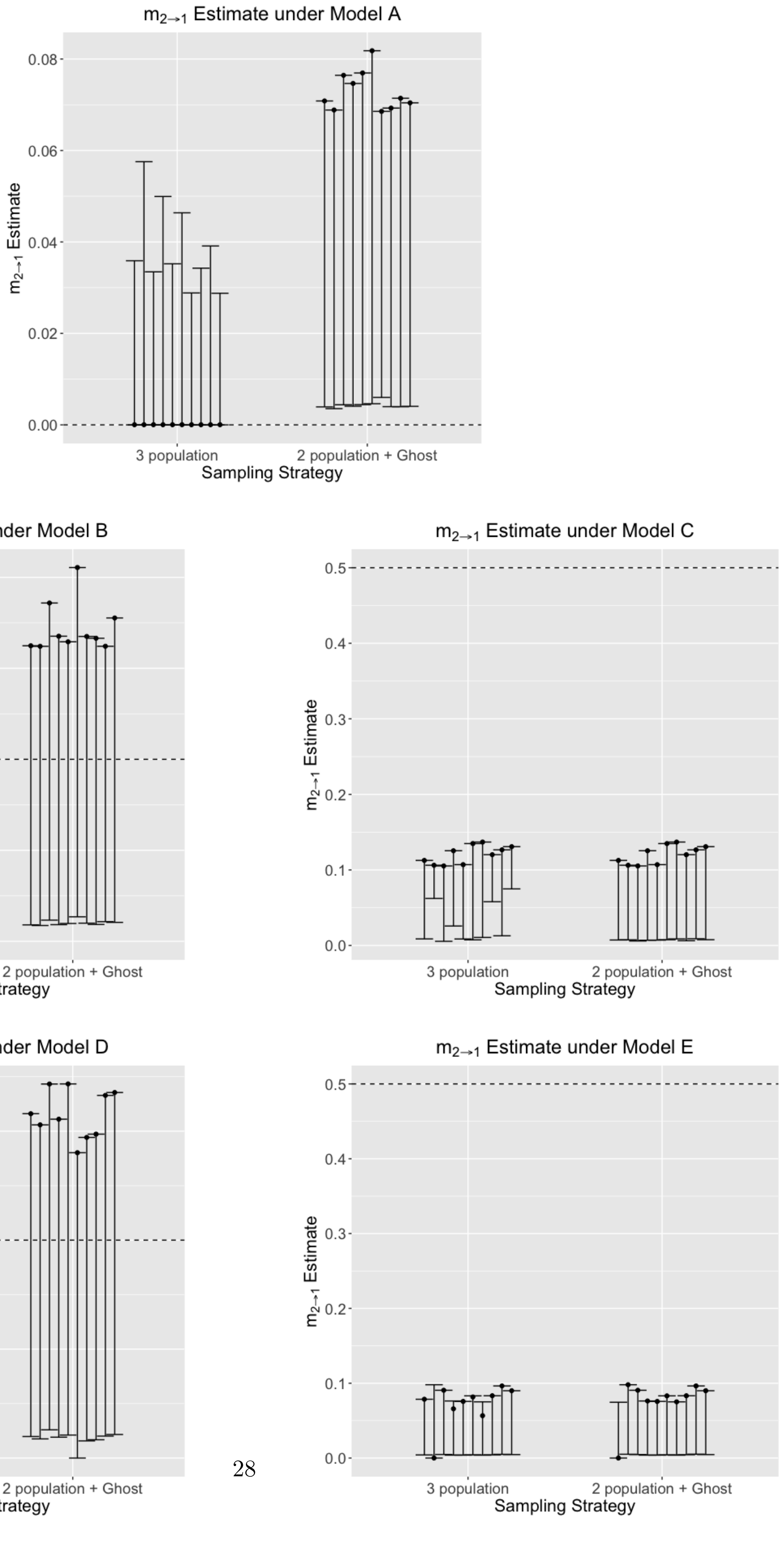

**Figure 26:** Scaled migration rate ($m_{1\to2} = 4N_1u$, where $N_1$ is the effective population size of one of the sampled populations, and $u$ is the mutation rate per site per generation) estimate between one of the sampled populations and the unsampled 'ghost', using 2 genomic loci under models A-E, estimated using a 3-population model, and a 2-population model with a ghost outgroup. True simulated migration rates from the ghost vary per model, and are shown with the dotted line. Across all models, we consistently under-estimate the migration rate from the 'ghost', especially with increased gene flow (models C and E).

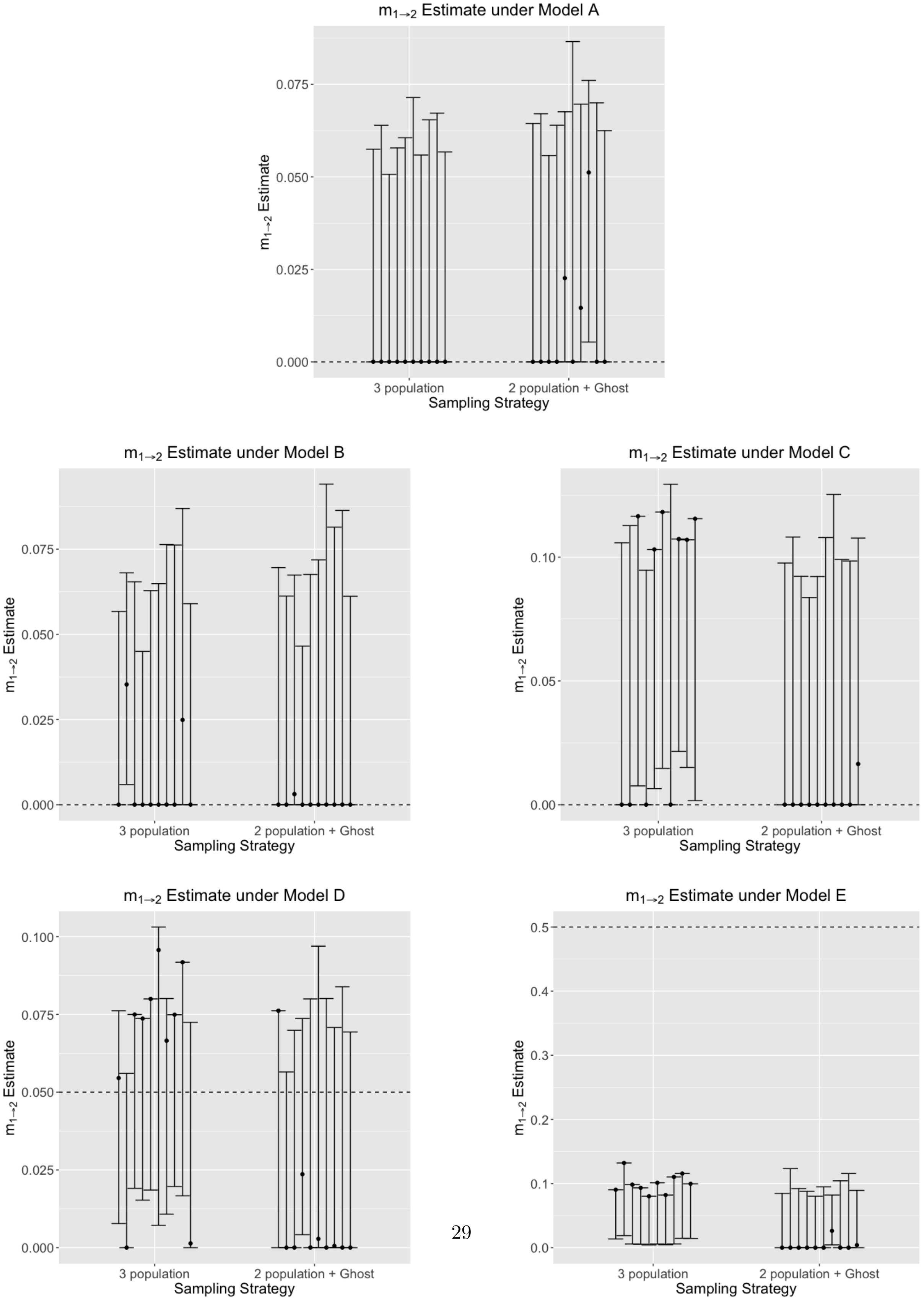

**Figure 27:** Scaled migration rate ($m_{1\to2} = 4N_1u$, where $N_1$ is the effective population size of one of the sampled populations, and $u$ is the mutation rate per site per generation) estimate between one of the sampled populations and the unsampled 'ghost', using 5 genomic loci under models A-E, estimated using a 3-population model, and a 2-population model with a ghost outgroup. True simulated migration rates from the ghost vary per model, and are shown with the dotted line. Across all models, we consistently under-estimate the migration rate from the 'ghost', especially with increased gene flow (models C and E).

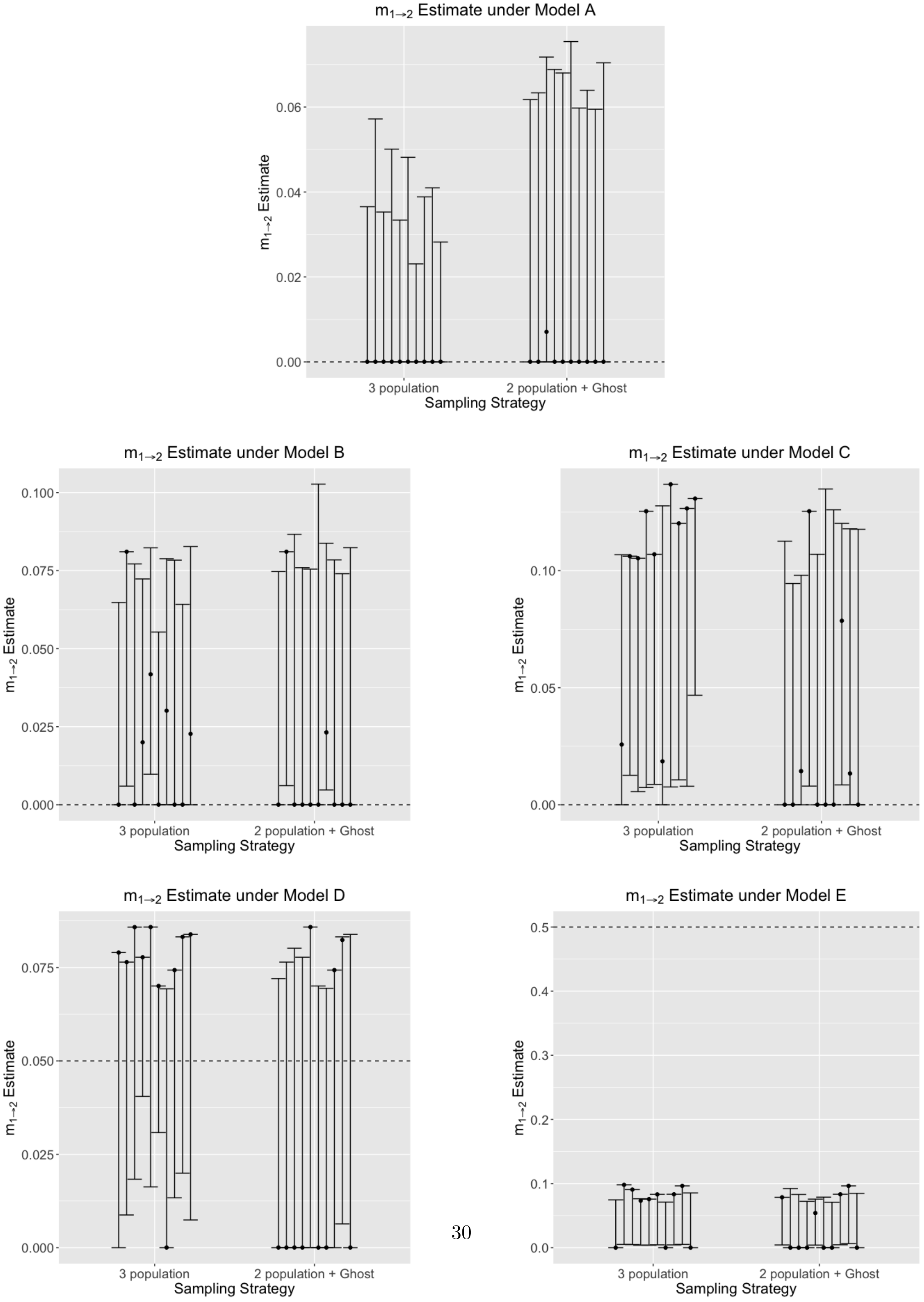

**Figure 28:** Scaled migration rate ($m_{2\to3} = 4N_2u$, where $N_2$ is the effective population size of the 'ghost', and $u$ is the mutation rate per site per generation) estimate between the common ancestor of the two sampled populations and the unsampled 'ghost', using 2 genomic loci under models A-E, estimated using a 3-population model, and a 2-population model with a ghost outgroup. True simulated migration rates from the ghost are 0.0, and are shown with the dotted line. Across all models, we consistently over-estimate the migration rate from the 'ghost', especially with increased gene flow (models C and E).

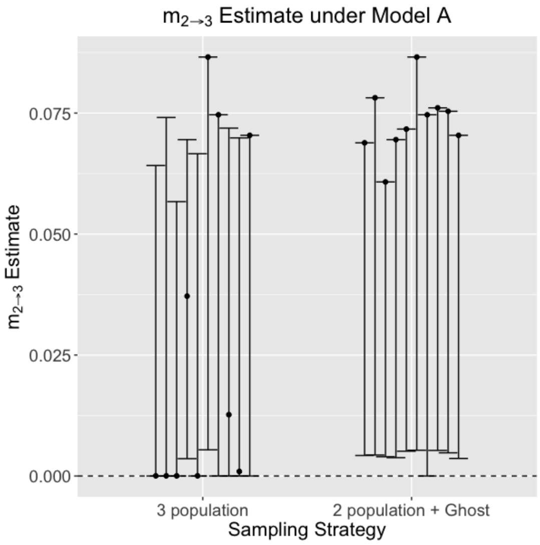


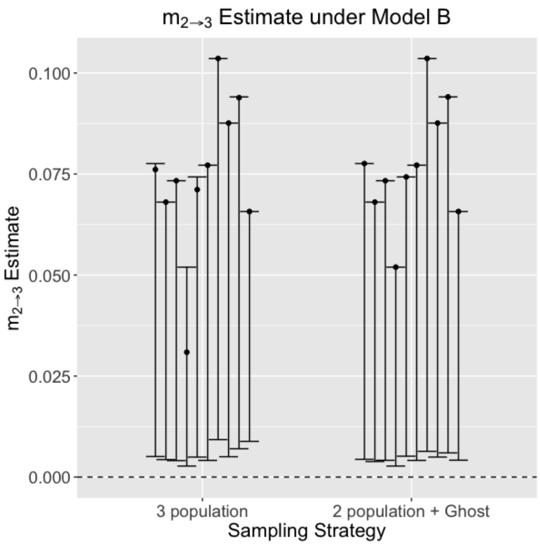


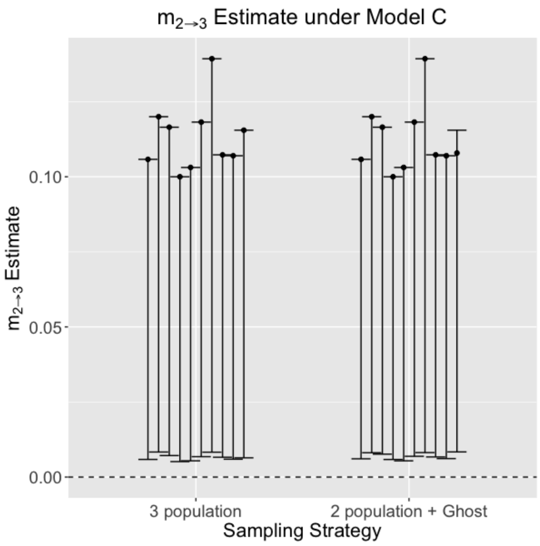


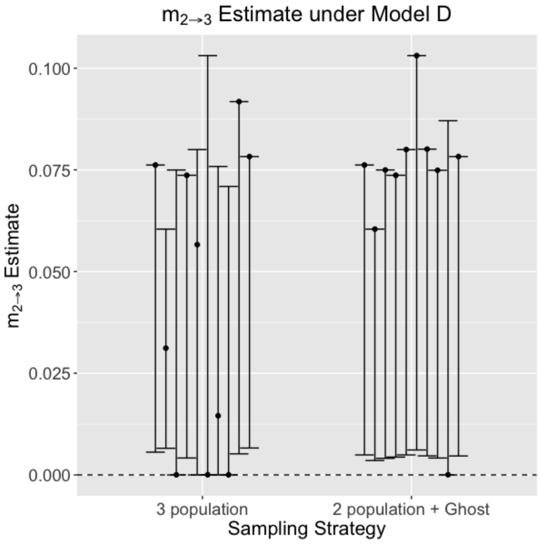


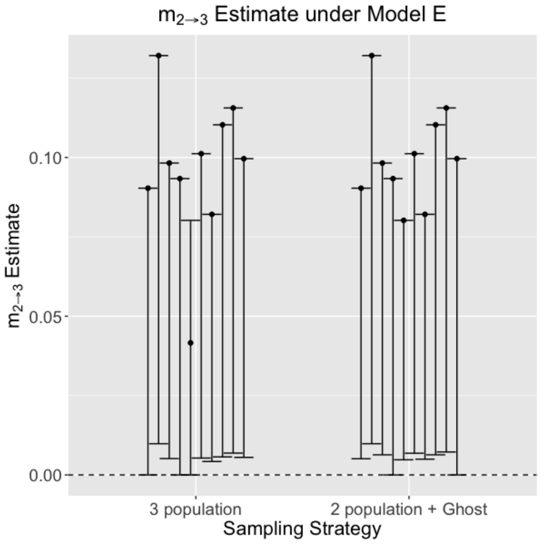

**Figure 29:** Scaled migration rate ($m_{2\to3} = 4N_2u$, where $N_2$ is the effective population size of the 'ghost', and $u$ is the mutation rate per site per generation) estimate between the common ancestor of the two sampled populations and the unsampled 'ghost', using 5 genomic loci under models A-E, estimated using a 3-population model, and a 2-population model with a ghost outgroup. True simulated migration rates from the ghost are 0.0, and are shown with the dotted line. Across all models, we consistently over-estimate the migration rate from the 'ghost', especially with increased gene flow (models C and E).

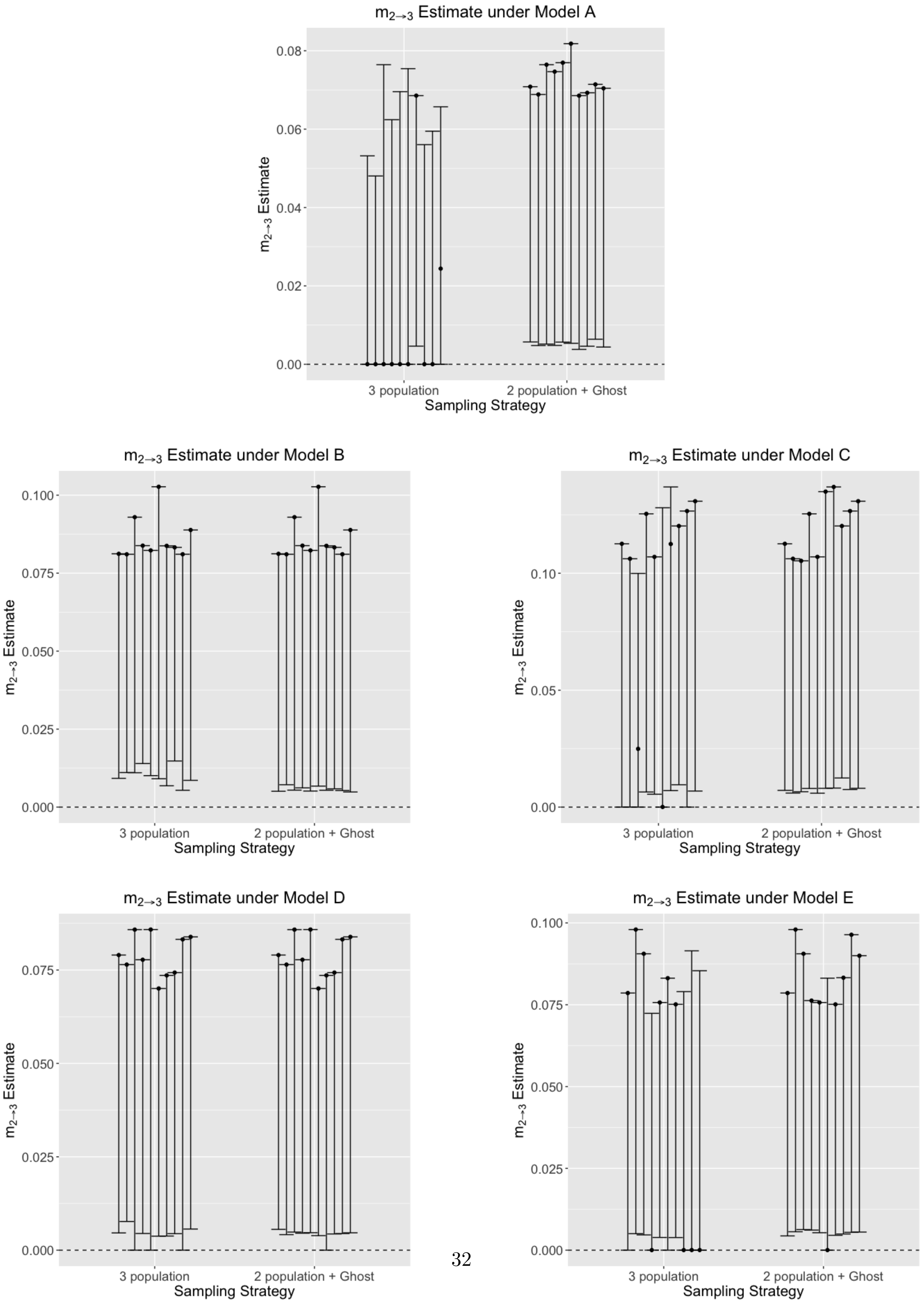

**Figure 30:** Scaled migration rate ($m_{3\to2} = 4N_3u$, where $N_3$ is the effective population size of the common ancestor of the two sampled populations, and $u$ is the mutation rate per site per generation) estimate between the common ancestor of the two sampled populations and the unsampled 'ghost', using 2 genomic loci under models A-E, estimated using a 3-population model, and a 2-population model with a ghost outgroup. True simulated migration rates from the ghost are 0.0, and are shown with the dotted line. Across all models, we consistently over-estimate the migration rate from the 'ghost', especially with increased gene flow (models C and E).

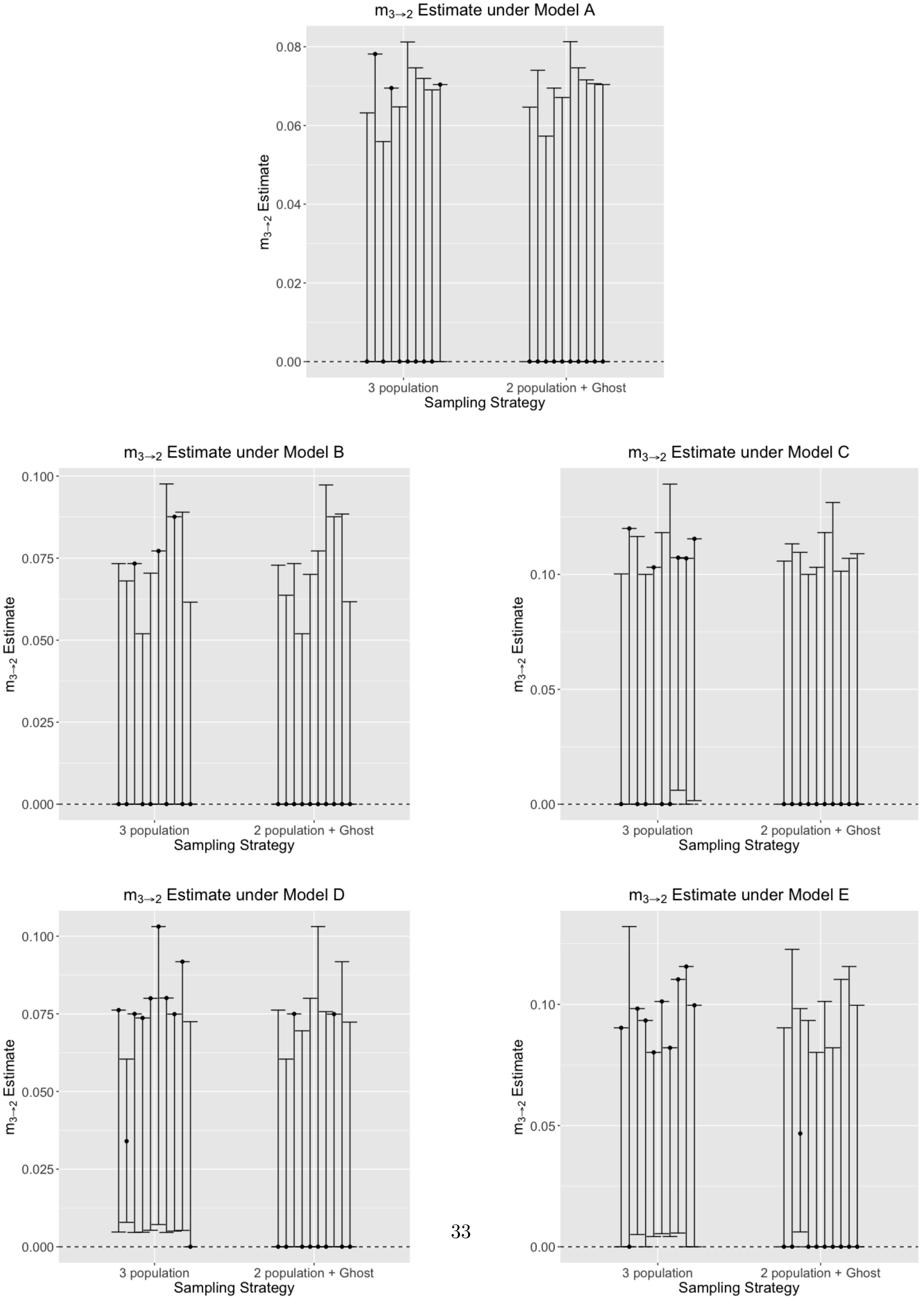

**Figure 31:** Scaled migration rate ($m_{3\to2} = 4N_3u$, where $N_3$ is the effective population size of the common ancestor of the two sampled populations, and $u$ is the mutation rate per site per generation) estimate between the common ancestor of the two sampled populations and the unsampled 'ghost', using 5 genomic loci under models A-E, estimated using a 3-population model, and a 2-population model with a ghost outgroup. True simulated migration rates from the ghost are 0.0, and are shown with the dotted line. Across all models, we consistently over-estimate the migration rate from the 'ghost', especially with increased gene flow (models C and E).

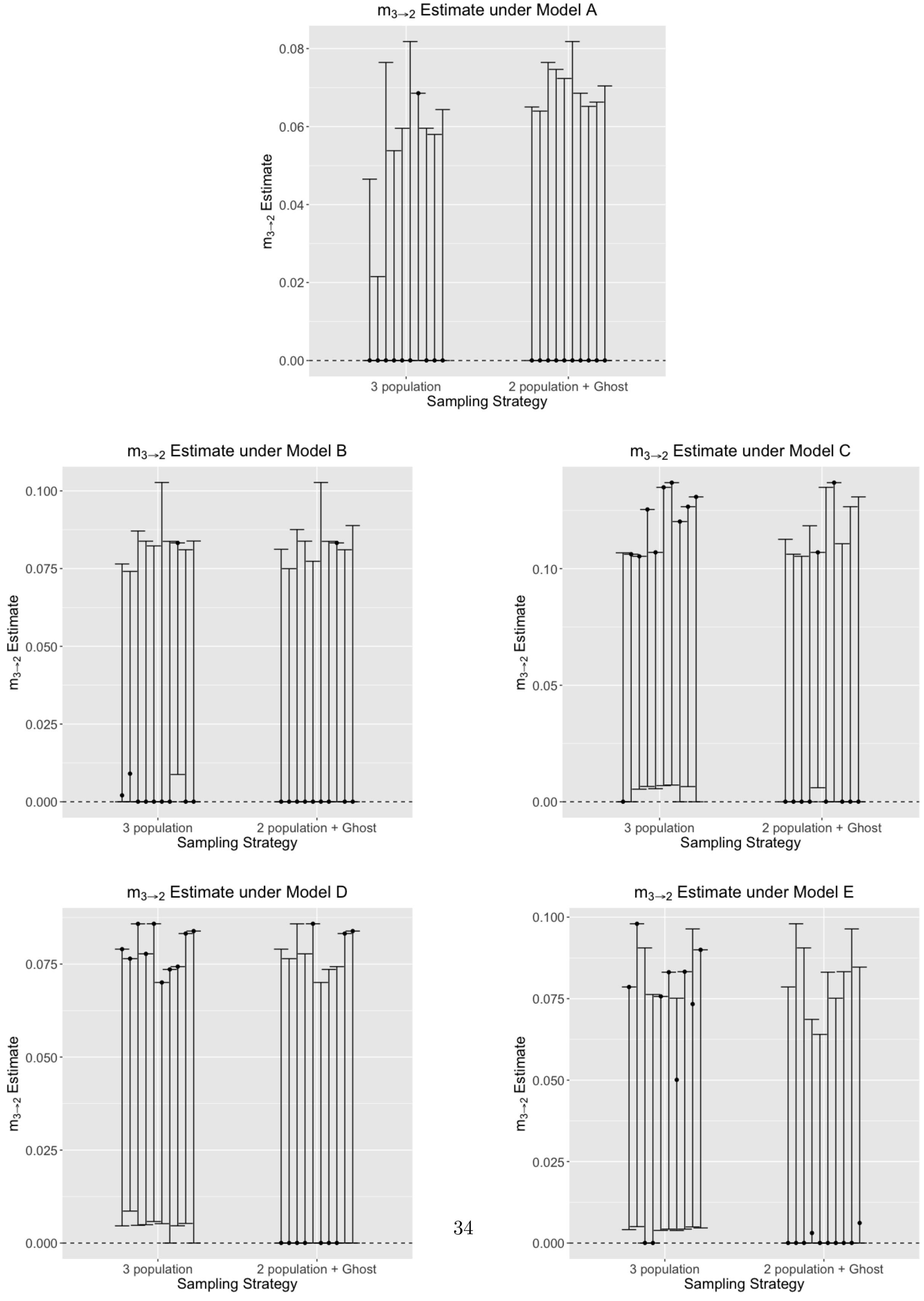

**Figure 32:** Bias in estimates of divergence times $t_0$ between the two sampled populations, under scenarios A-E, estimated under the (a) two population IM model, (b) three population IM model, and (c) two population IM model, with an outgroup ghost. Biases are shown across datasets from 2 loci (black), and 5 loci (red) sampling schemes. Overall bias in estimates is reduced with increasing number of loci. Divergence time is over-estimated in models with zero, or low gene flow from the ghost (A, B, D), but under estimated in models with high gene flow from the ghost (C, E).

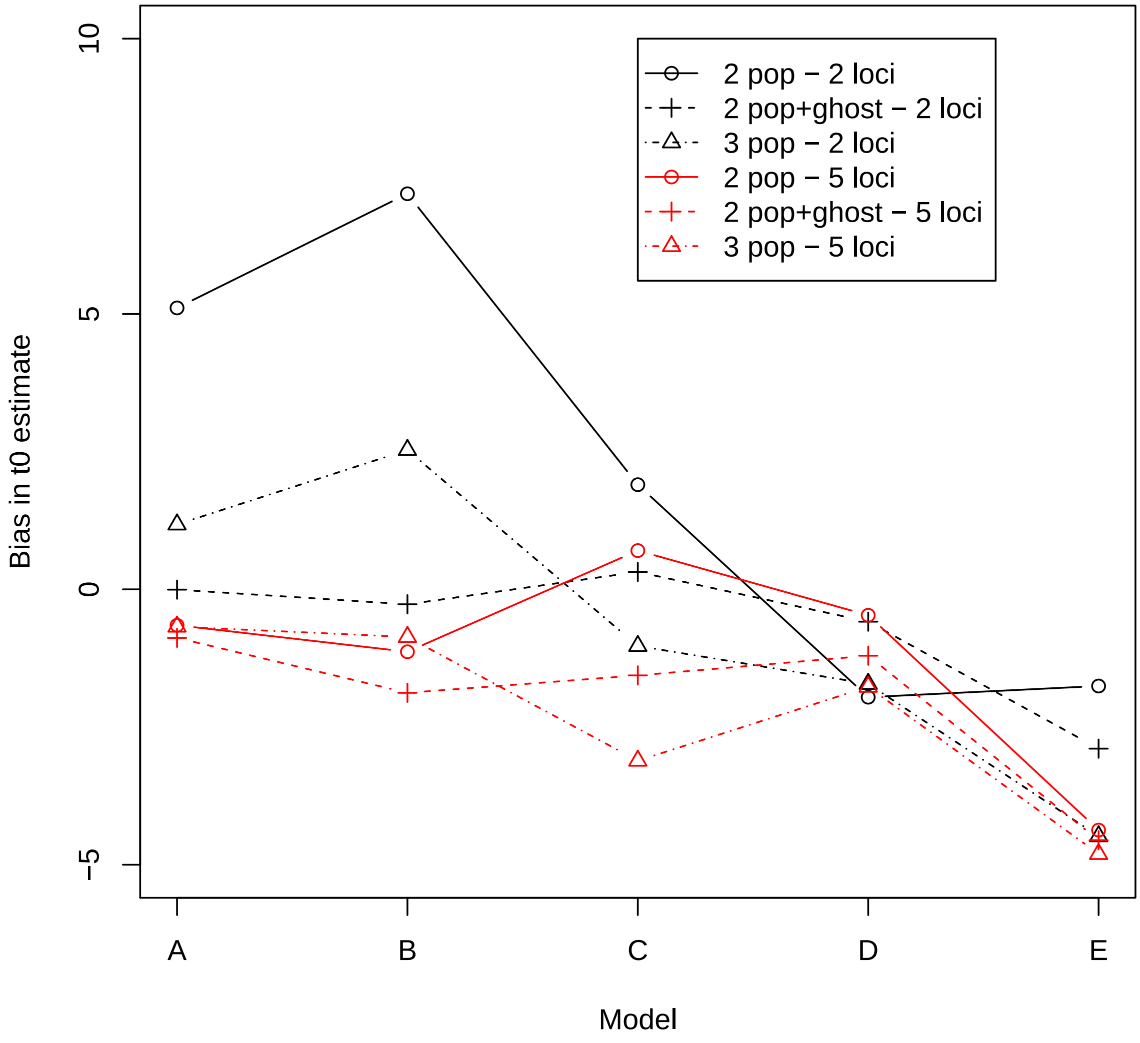

**Figure 33:** Bias in estimates of mutation scaled population size $q_0$ of one sampled population, under scenarios A-E, estimated under the (a) two population IM model, (b) three population IM model, and (c) two population IM model, with an outgroup ghost. Biases are shown across datasets from 2 loci (black), and 5 loci (red) sampling schemes. Mutation scaled population is consistently over-estimated with increased degree of gene flow from the unsampled ghost.

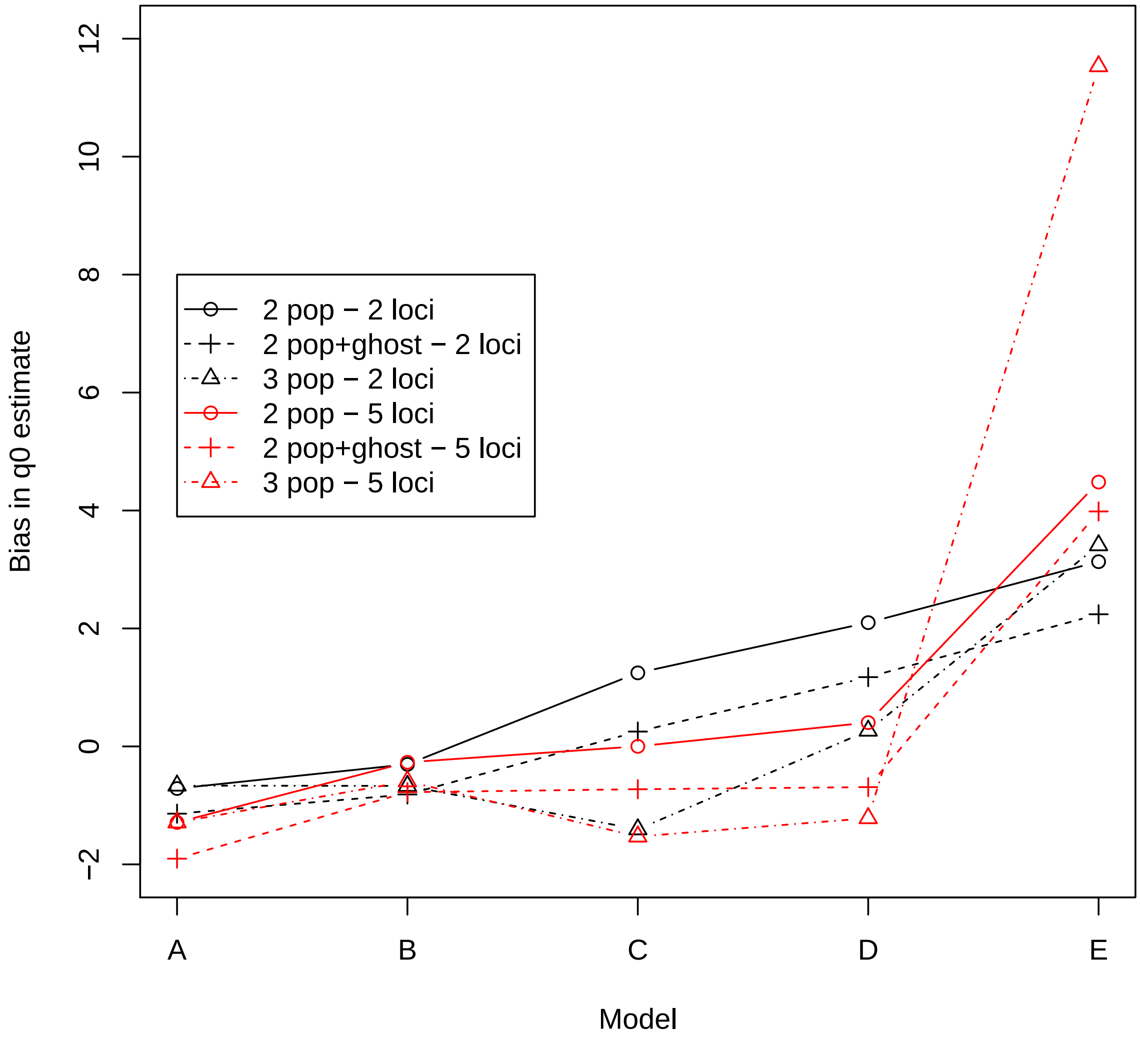

**Figure 34:** Bias in estimates of mutation scaled population size $q_1$ of one sampled population, under scenarios A-E, estimated under the (a) two population IM model, (b) three population IM model, and (c) two population IM model, with an outgroup ghost. Biases are shown across datasets from 2 loci (black), and 5 loci (red) sampling schemes. Mutation scaled population is consistently over-estimated with increased degree of gene flow from the unsampled ghost.

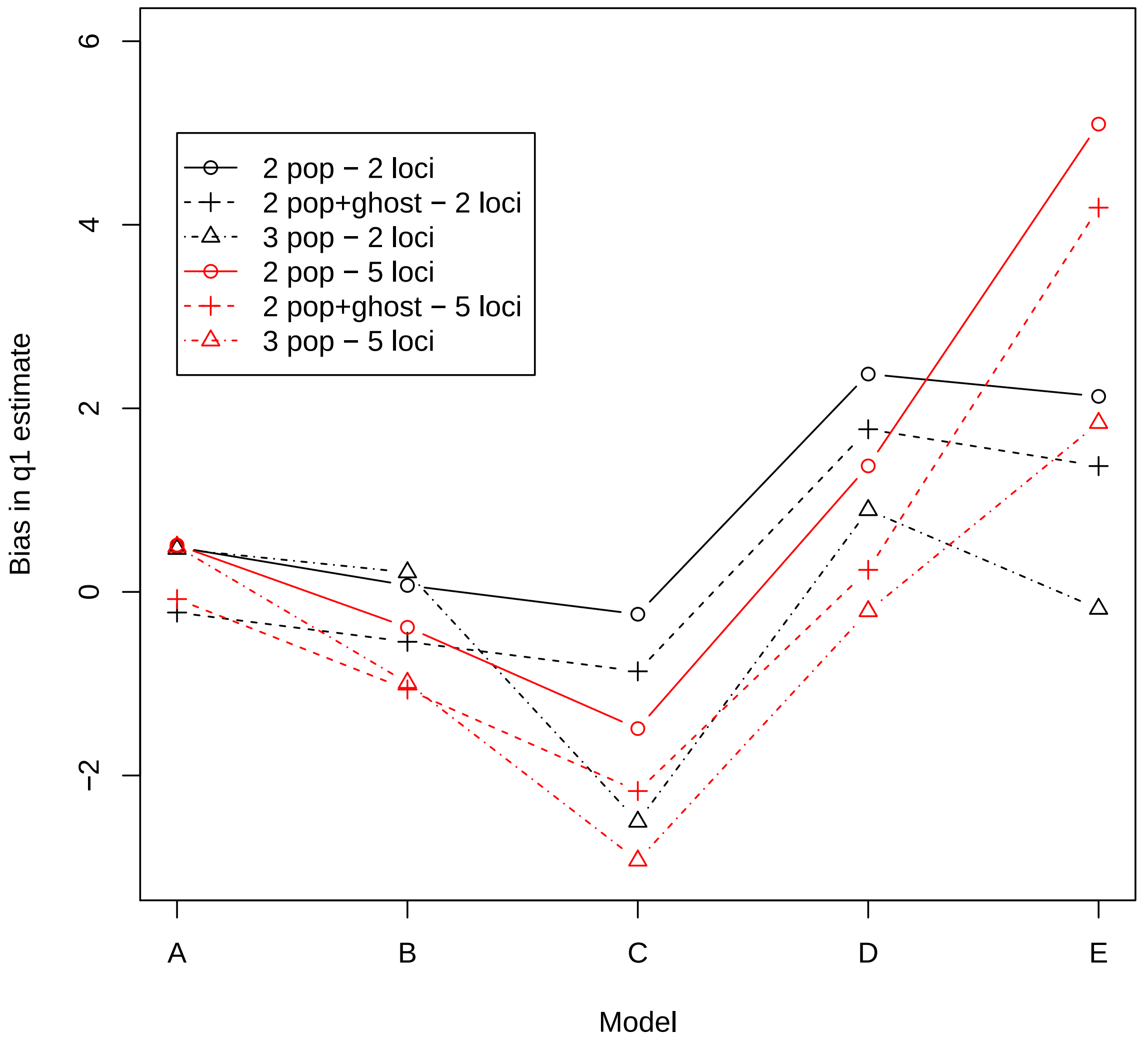

**Figure 35:** Bias in estimates of mutation scaled population size $q_2$ of the common ancestor of the two sampled populations, under scenarios A-E, estimated under the (a) two population IM model, (b) three population IM model, and (c) two population IM model, with an outgroup ghost. Biases are shown across datasets from 2 loci (black), and 5 loci (red) sampling schemes. Mutation scaled population is consistently over-estimated with increased degree of gene flow from the unsampled ghost.

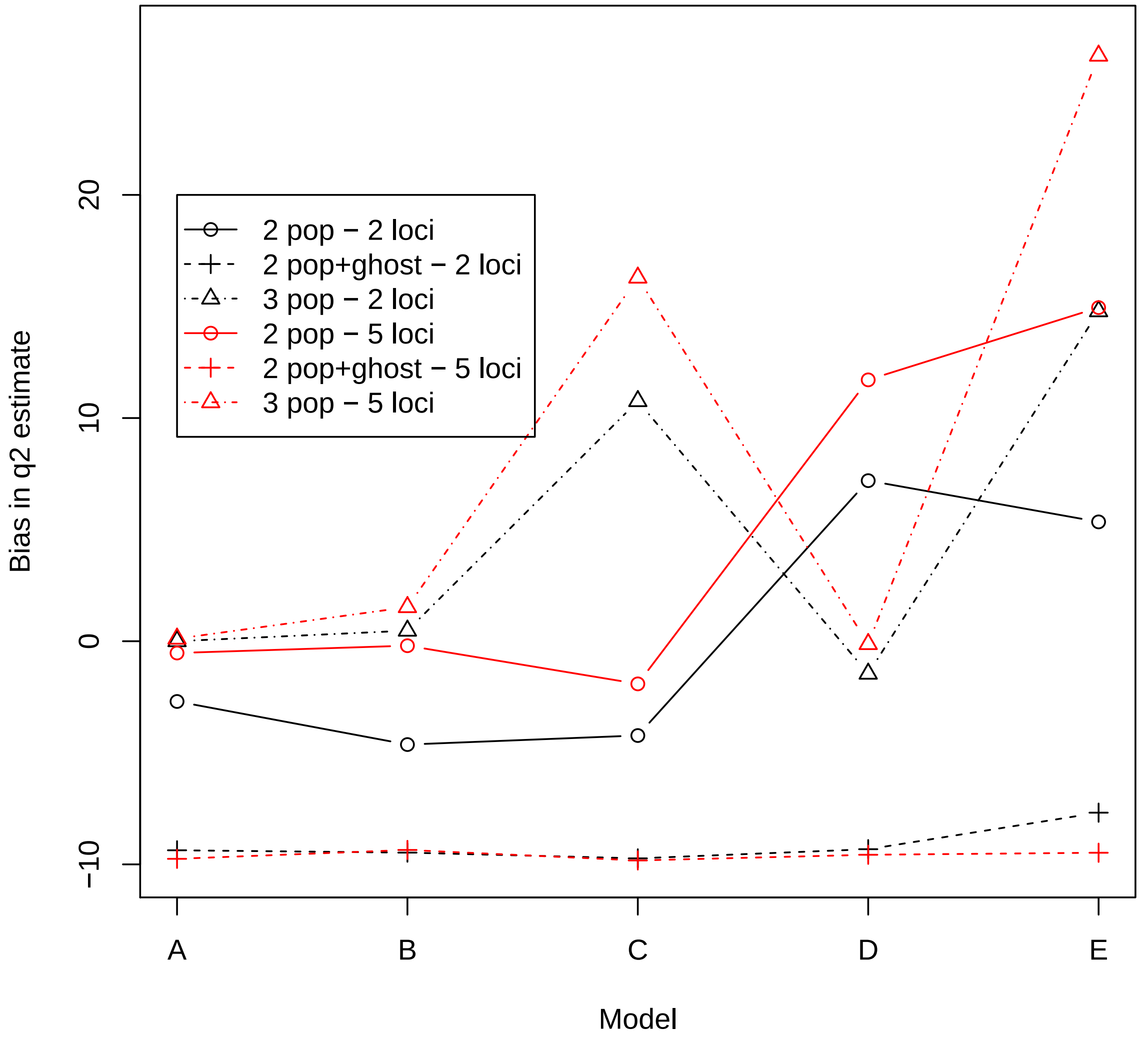

**Figure 36:** Bias in estimates of mutation scaled population size $q_3$ of the ghost population, under scenarios A-E, estimated under the (a) three population IM model, and (b) two population IM model, with an outgroup ghost. Biases are shown across datasets from 2 loci (black), and 5 loci (red) sampling schemes. Mutation scaled population is consistently over-estimated with increased degree of gene flow from the unsampled ghost.

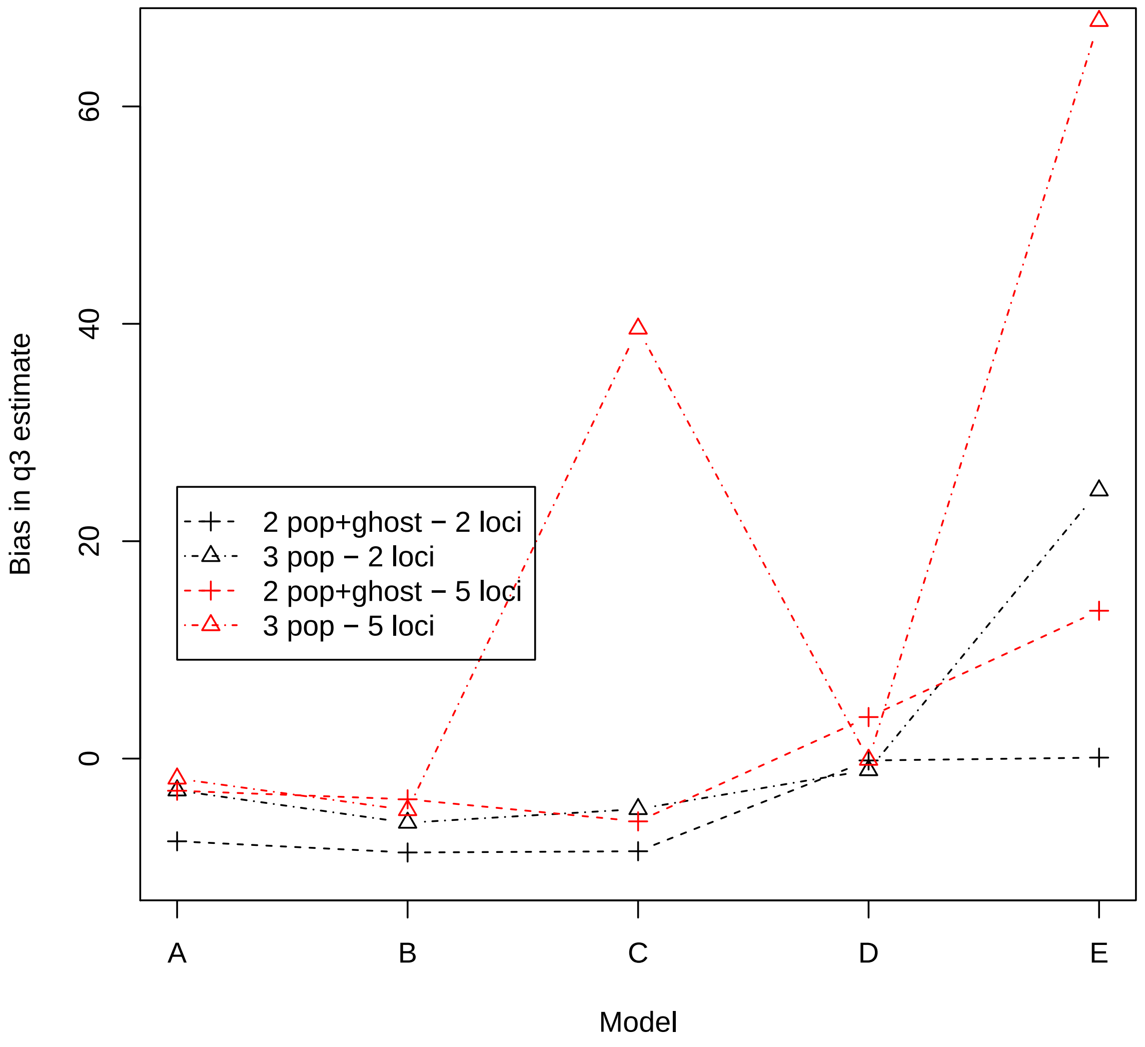

**Figure 37:** Bias in estimates of mutation scaled population size $q_4$ of the common ancestor of the ghost population, and the two sampled populations under scenarios A-E, estimated under the (a) three population IM model, and (b) two population IM model, with an outgroup ghost. Biases are shown across datasets from 2 loci (black), and 5 loci (red) sampling schemes. Mutation scaled population is consistently over-estimated with increased degree of gene flow from the unsampled ghost.

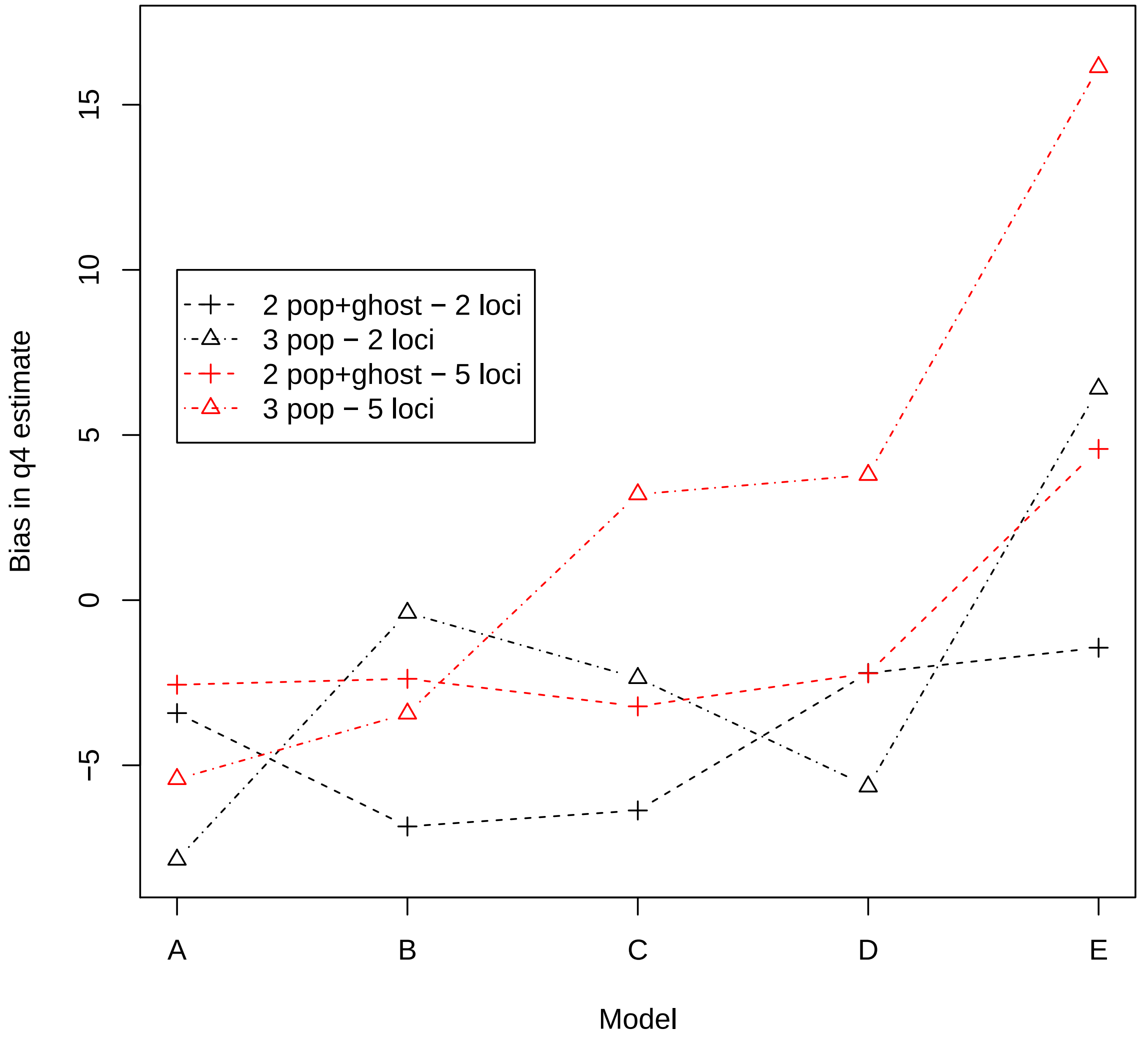

**Figure 38:** Bias in estimates of migration rates $m_{01}$ between the two sampled population, under scenarios A-E, estimated under the (a) two population IM model, (b) three population IM model, and (c) two population IM model, with an outgroup ghost. Biases are shown across datasets from 2 loci (black), and 5 loci (red) sampling schemes.

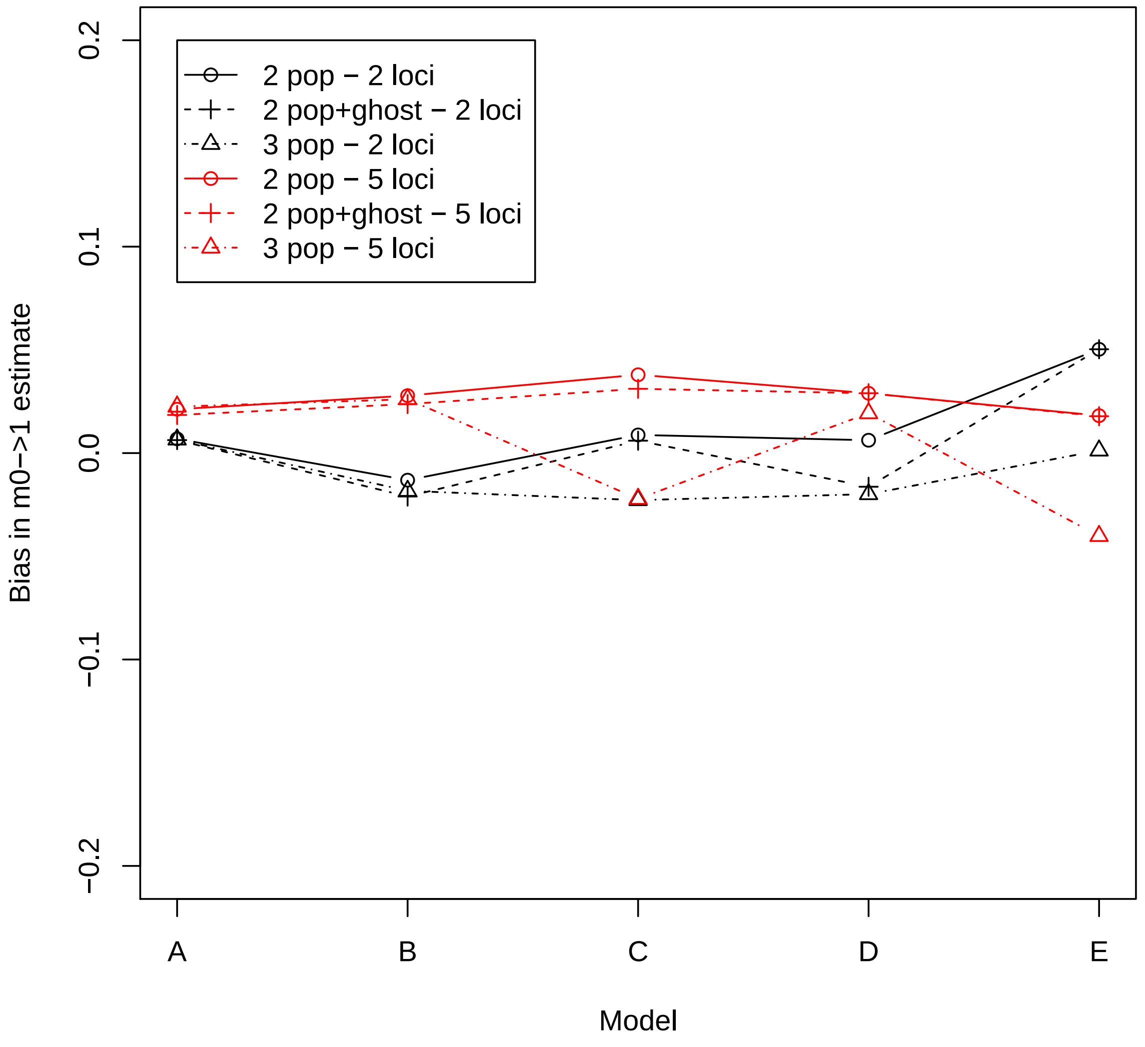

**Figure 39:** Bias in estimates of migration rates $m_{10}$ between the two sampled population, under scenarios A-E, estimated under the (a) two population IM model, (b) three population IM model, and (c) two population IM model, with an outgroup ghost. Biases are shown across datasets from 2 loci (black), and 5 loci (red) sampling schemes.

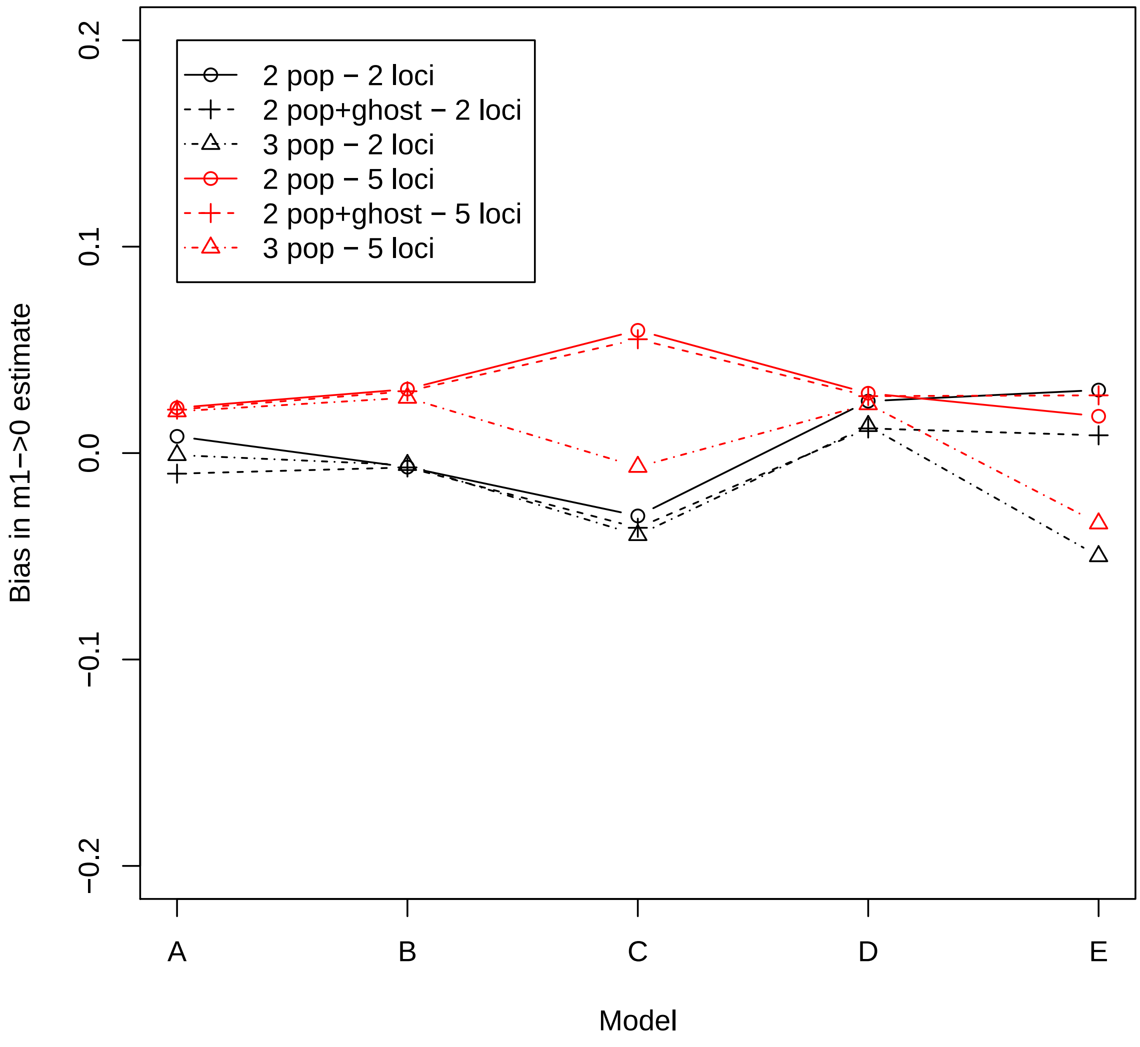

**Figure 40:** Bias in estimates of migration rates $m_{02}$ between the sampled population 0 and the ghost, under scenarios A-E, estimated under the (a) two population IM model, (b) three population IM model, and (c) two population IM model, with an outgroup ghost. Biases are shown across datasets from 2 loci (black), and 5 loci (red) sampling schemes. Migration rates are always under-estimated under scenarios C and E, where there's greater gene flow from the ghost.

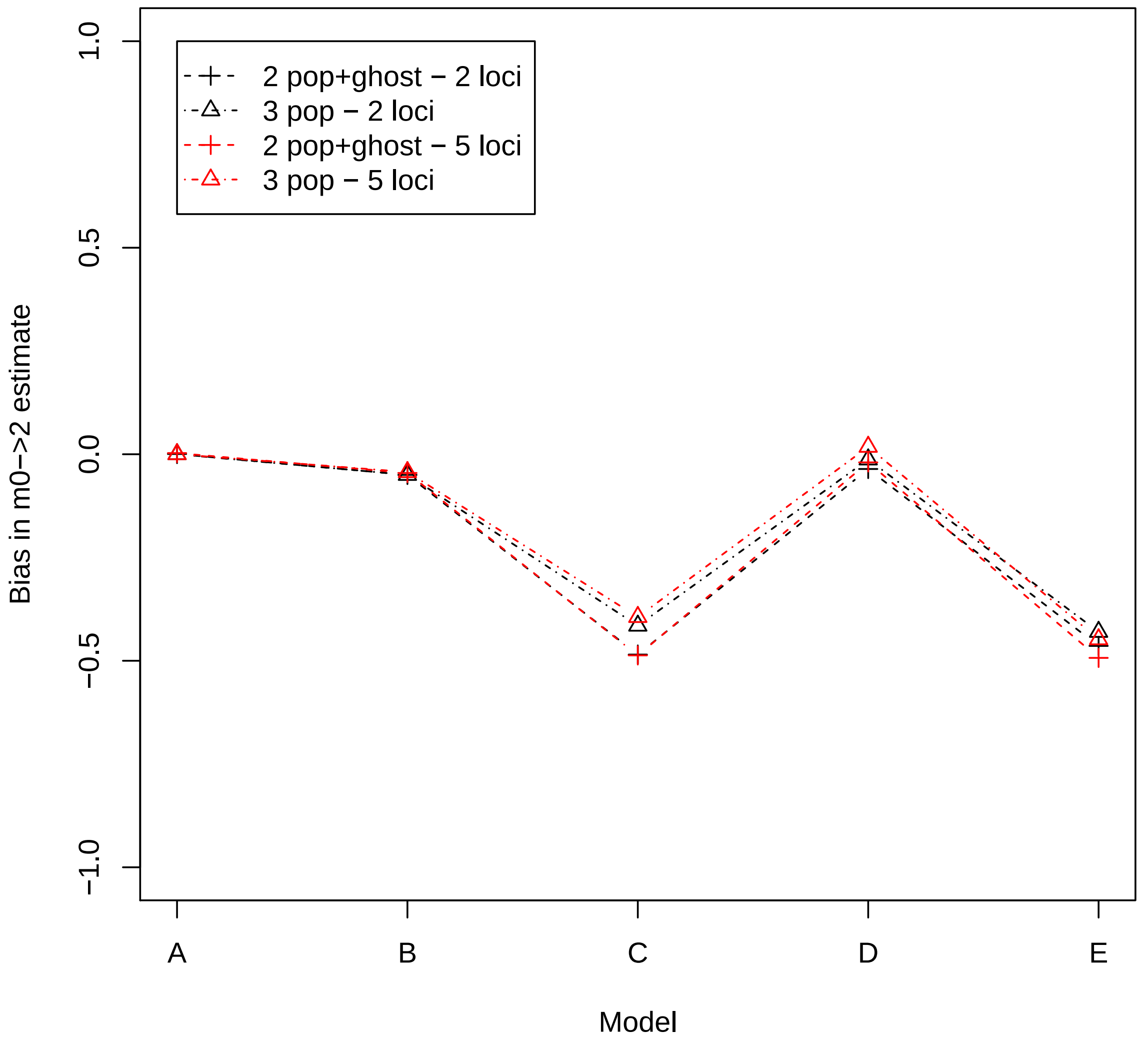

**Figure 41:** Bias in estimates of migration rates $m_{20}$ between the sampled population 0 and the ghost, under scenarios A-E, estimated under the (a) two population IM model, (b) three population IM model, and (c) two population IM model, with an outgroup ghost. Biases are shown across datasets from 2 loci (black), and 5 loci (red) sampling schemes. Migration rates are always under-estimated under scenarios C and E, where there's greater gene flow from the ghost.

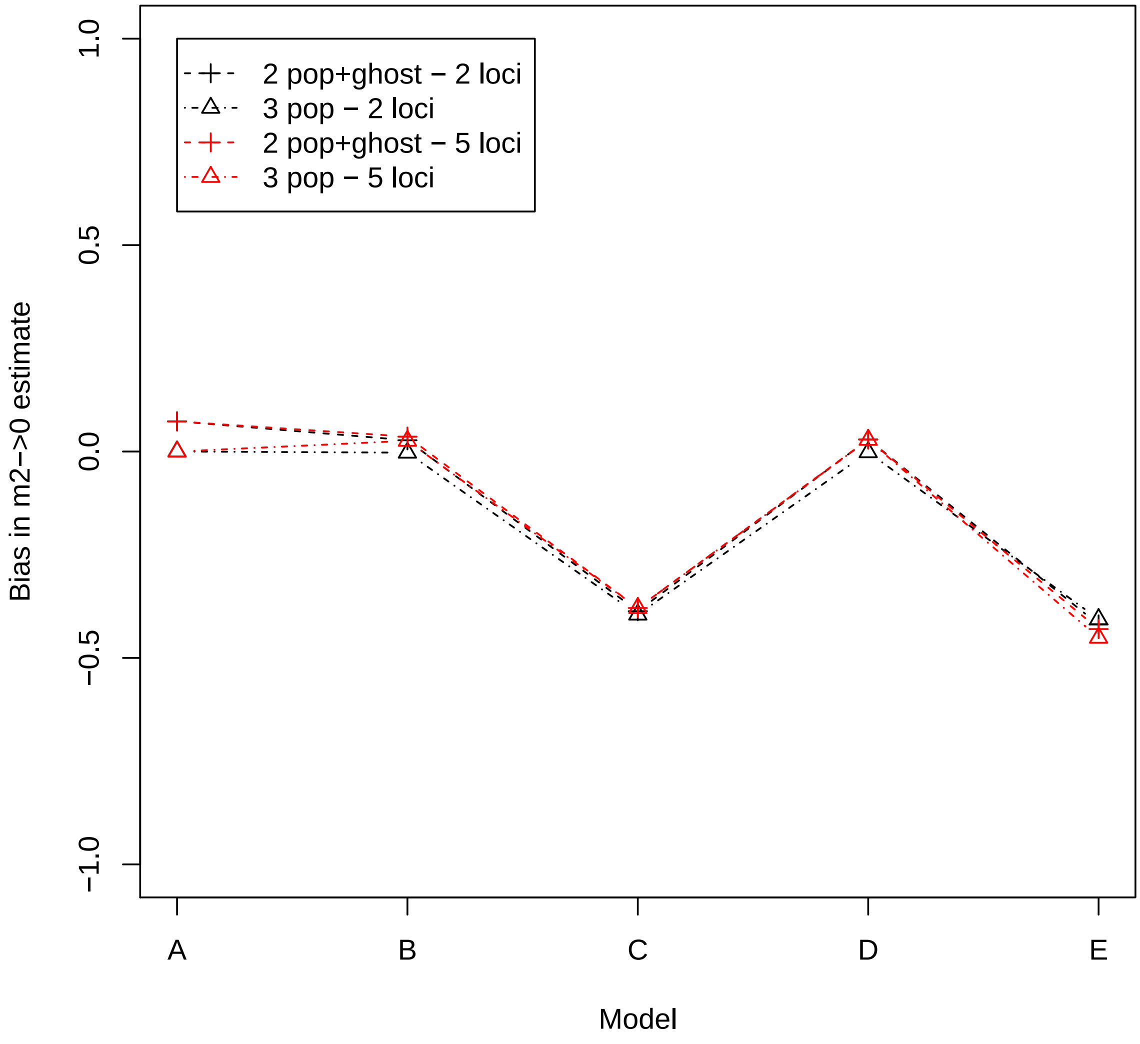

**Figure 42:** Bias in estimates of migration rates $m_{12}$ between the sampled population 1 and the ghost, under scenarios A-E, estimated under the (a) two population IM model, (b) three population IM model, and (c) two population IM model, with an outgroup ghost. Biases are shown across datasets from 2 loci (black), and 5 loci (red) sampling schemes. Migration rates are always under-estimated under scenarios C and E, where there's greater gene flow from the ghost.

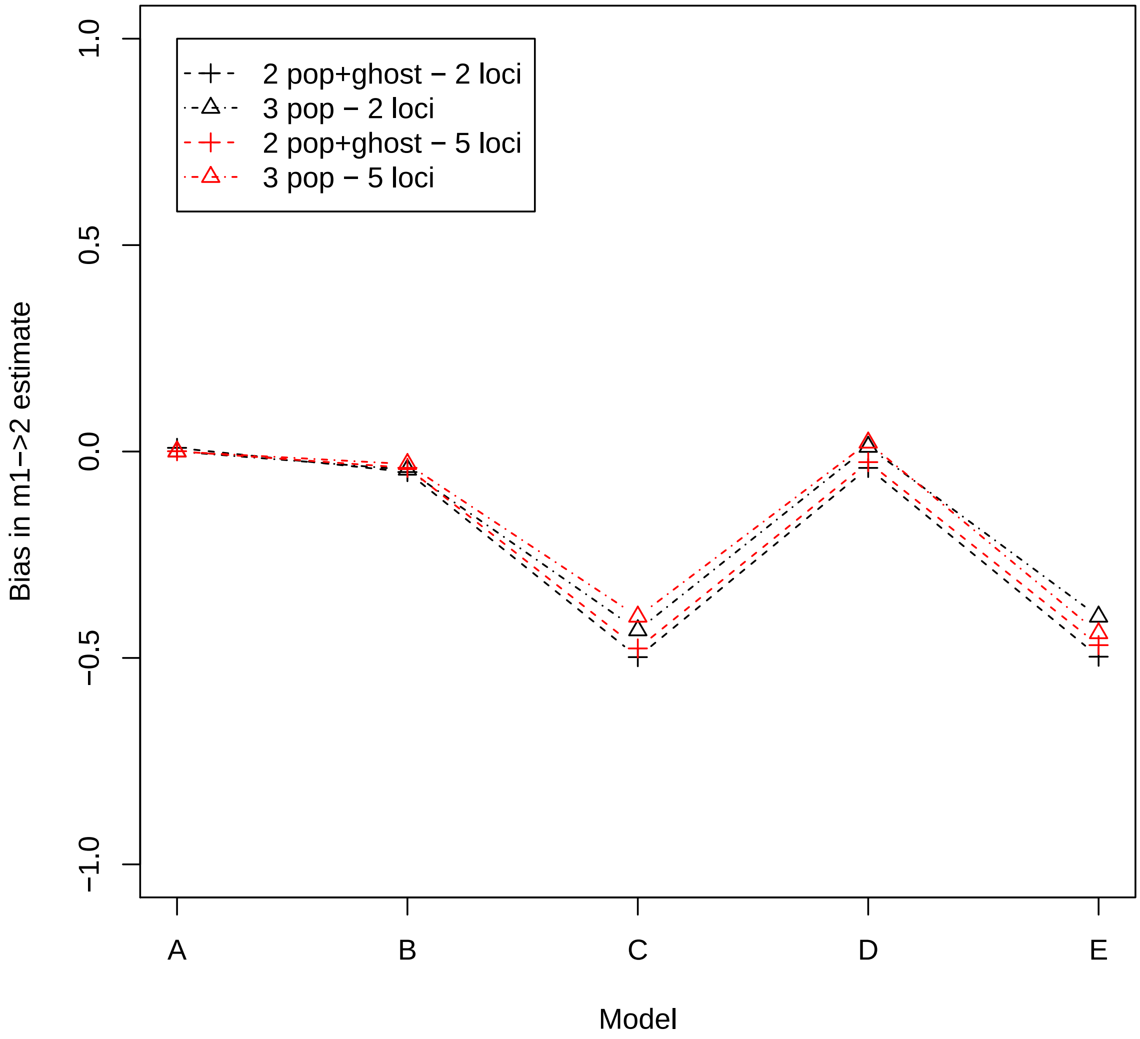

**Figure 43:** Bias in estimates of migration rates $m_{21}$ between the sampled population 1 and the ghost, under scenarios A-E, estimated under the (a) two population IM model, (b) three population IM model, and (c) two population IM model, with an outgroup ghost. Biases are shown across datasets from 2 loci (black), and 5 loci (red) sampling schemes. Migration rates are always under-estimated under scenarios C and E, where there's greater gene flow from the ghost.

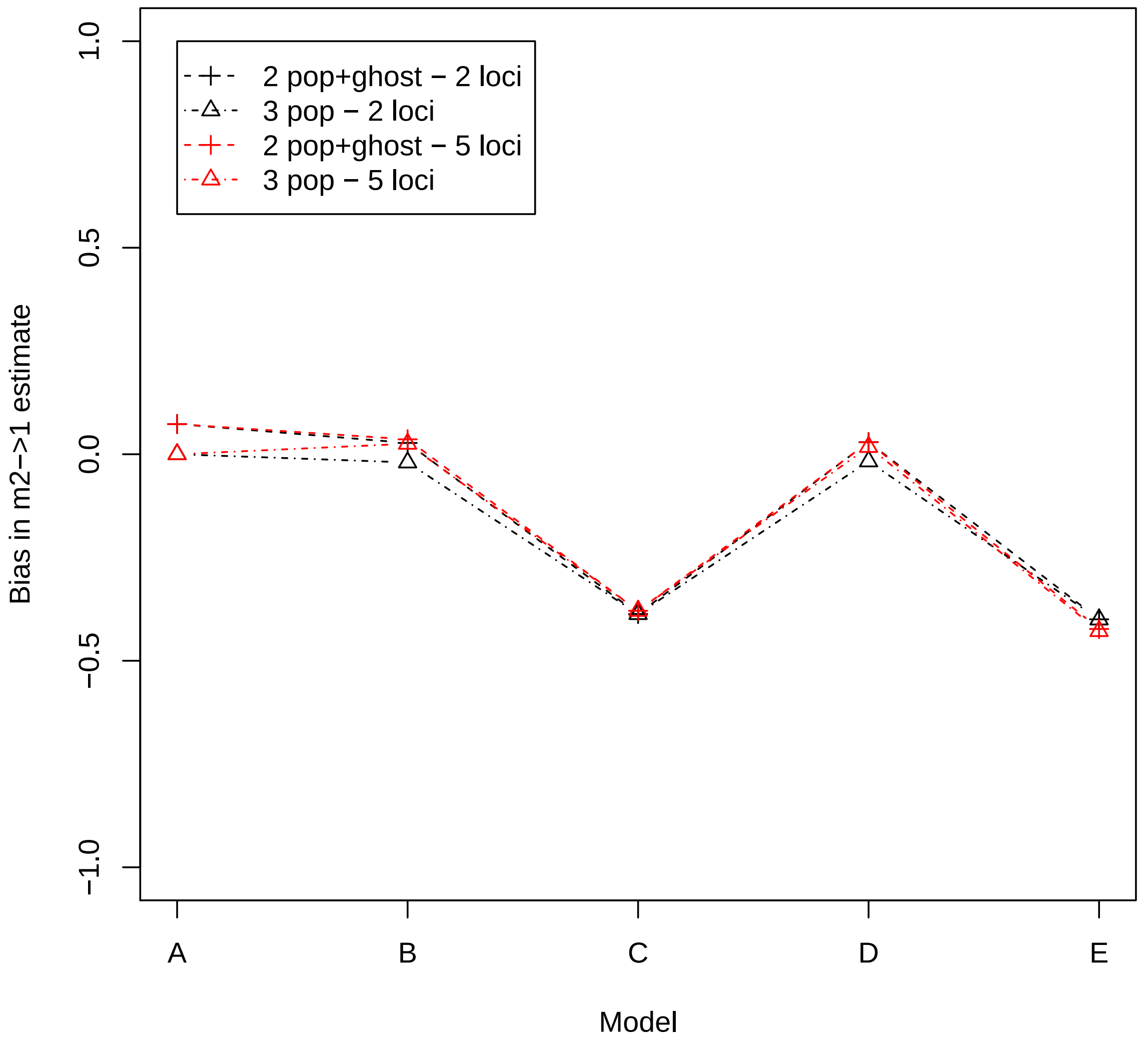

**Figure 44:** Bias in estimates of migration rates $m_{23}$ between the common ancestor of sampled populations and the ghost, under scenarios A-E, estimated under the (a) two population IM model, (b) three population IM model, and (c) two population IM model, with an outgroup ghost. Biases are shown across datasets from 2 loci (black), and 5 loci (red) sampling schemes. Migration rates are always under-estimated under scenarios C and E, where there's greater gene flow from the ghost.

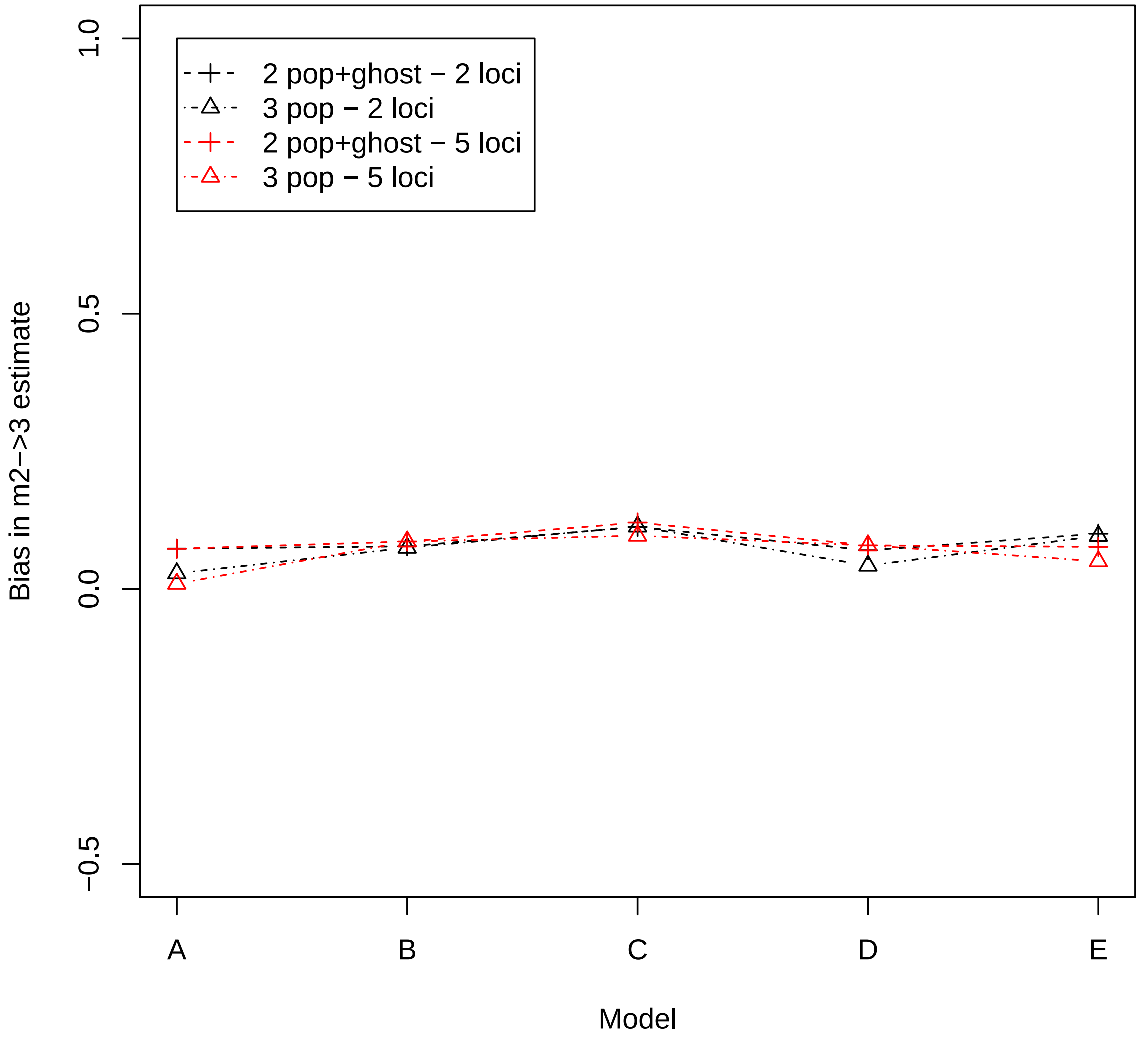

**Figure 45:** Bias in estimates of migration rates $m_{32}$ between the common ancestor of sampled populations and the ghost, under scenarios A-E, estimated under the (a) two population IM model, (b) three population IM model, and (c) two population IM model, with an outgroup ghost. Biases are shown across datasets from 2 loci (black), and 5 loci (red) sampling schemes. Migration rates are always under-estimated under scenarios C and E, where there's greater gene flow from the ghost.

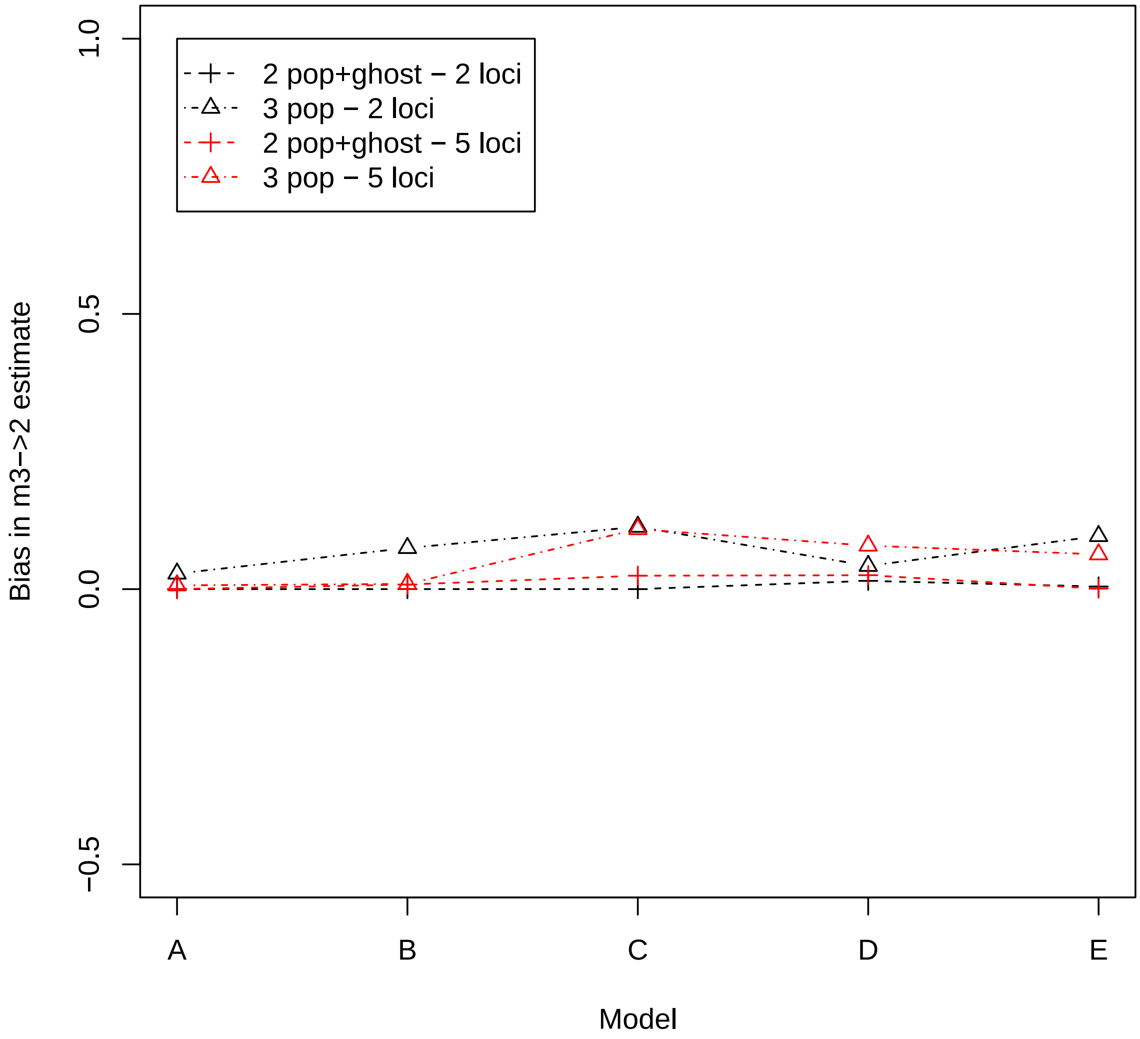
